# Spectral Phenotyping Reveals Time-Specific QTLs in Field-Grown Lettuce

**DOI:** 10.64898/2026.03.16.711173

**Authors:** Sarah Mehrem, Anne Zijl, Myrthe de Haan, Guido van den Ackerveken, Basten L. Snoek

## Abstract

Lettuce (*Lactuca sativa*) is an important field crop, but our understanding of its phenotypic variation and underlying genetics under natural field conditions remains limited, posing challenges for identifying effective crop breeding targets. Longitudinal hyperspectral phenotyping allows for non-invasive monitoring of crop performance under diverse agricultural conditions. In this study, we used hyperspectral imaging to assess the phenotypic variation of almost 200 different field-grown lettuce varieties, following the same plants from just after seedling- to flowering-stage. With automated image processing, we extracted a wide range of spectral phenotypes related to metabolite content, growth efficiency, and environmental stress responses, creating a multi-dimensional time-resolved data set. Principal component analysis (PCA) revealed the major axes of spectral variation over time, and highlighted differences in spectral patterns among lettuce genotypes. Integrating on-site weather data, we modelled G×E interactions of reflectance, revealing regions of the lettuce vegetation spectrum that are primarily shaped by genotype and/or environment. We estimated phenotypic plasticity in response to time, temperature and rainfall using best linear unbiased predictions (BLUPs), capturing genotype-specific developmental trajectories and responses to the environment. We used genome-wide association studies (GWAS) to identify quantitative trait loci (QTLs) of PC-based, single and BLUP-based phenotypes, disentangling the genetic architecture of spectral lettuce phenotypes from major axes of variation down to single wavelength spectral plasticity. These findings provide new insights into the genome-wide genetic regulation and dynamics of spectral phenotypes in field grown lettuce.

## Introduction

Crops grown in an open field face environmental stressors, like fluctuating temperatures, drought and heavy rainfall, posing a serious risk to production. Extreme weather phenomena occur more frequently and with greater intensities due to anthropogenic climate change [1]. To secure stable yields under such highly variable conditions, crop breeding programs must focus on producing more resilient varieties. Lettuce (*Lactuca sativa*), a major field crop, provides a relevant case study in which genetic and phenotypic dynamics under field conditions can reveal breeding targets for more resilient, high-performing varieties.

Resilience, yield, and crop architecture are dynamic agronomic traits of lettuce, changing over time and with the environment [2–4]. Their manifestation reflects a complex interplay of genotype, development, and environment, which complicates the detection of underlying quantitative trait loci (QTLs) using mapping approaches including genome-wide association studies (GWAS) [5]. These developmental trajectories also vary among horticultural types of lettuce (Butterhead, Cos, Latin, Cutting, Stalk and Oilseed), each with distinct growth and morphological features. Moreover, QTLs identified under lab or greenhouse conditions often fail to translate into improved field performance [2]. Together, this highlights the need to study trait expression and genetic architecture of lettuce in a temporal, field-relevant context.

Longitudinal phenotyping is a critical part of this effort, but such studies are demanding, both in terms of labor and logistics. Invasive phenotyping methods, like tissue sampling or destructive biomass measurements, additionally complicate repeated measurements of the same individuals over time [6]. Hyperspectral imaging circumvents some of these caveats, enabling high-resolution, non-destructive phenotyping. It has been successfully applied across various crops to capture both biochemical and morphological traits of the canopy with high precision [7]. Combined with weather monitoring and genetic information, spectral longitudinal phenotyping provides valuable insights into the mechanisms underlying trait variation in a field context [8–10]. Beyond raw reflectance, hyperspectral data is typically transformed into vegetation indices (e.g. NDVI) that serve as proxies for traits such as biomass, pigment composition, photosynthetic efficiency, and stress responses [7]. In lettuce, field-based hyperspectral phenotyping has been applied to estimate and monitor nutrient and plant pigment content, as well as for early detection of biotic stresses [11–13].

Field studies in lettuce have combined GWAS with hyperspectral phenotyping or drone-based RGB/multispectral imaging, yet both approaches provided limited to no temporal resolution [5, 14, 15]. These studies consistently identified major QTLs for anthocyanin and chlorophyll content (*RLL1-4*, *ANS*, *LsGLK*), as well as regulators of plant height and architecture such as *PhyC* and *LsTCP4*, but limited temporal resolution and narrow spectral ranges left trait dynamics underexplored [5, 16–18].

Although previous work identified major QTLs for pigments and architecture, limited temporal resolution left trait dynamics unexplored. Here, we integrated longitudinal spectral phenotyping with GWAS in 194 field-grown lettuce accessions. Spectral data was captured at 10 time points spanning development from seedling to flowering. We extracted several spectral phenotypes, including vegetation indices related to metabolite content, growth efficiency, and environmental stress responses. This high-dimensional dataset provided a rich resource to study how genetics and environment interact to shape trait dynamics in field-grown lettuce. Using dimensionality reduction, we described the major axes of spectral variation over time, linked to water content, horticultural type, and plant pigmentation. We used mixed linear models to estimate the effects of genetics, time, and environment on spectral phenotypes, revealing developmental and environmentally driven plasticity. Using GWAS, we uncovered the genetic architecture underlying spectral lettuce phenotypes, identifying both known and novel QTLs, including time-specific loci linked to developmental and environmental plasticity of anthocyanin pigmentation.

Altogether, these results demonstrate how longitudinal hyperspectral phenotyping can be used to investigate the genetic basis of phenotypic dynamics in field-grown lettuce across development and variable environmental conditions.

## Methods

### Field experiment

For this experiment 194 *Lactuca sativa* accessions (**Table S1**) were grown under field conditions near Maasbree in the Netherlands. Plants were germinated on the 25^th^ and 26^th^ of March 2021 and planted in the field in early April. Each accession was grown on two plots, where each plot contained 30 to 40 individual plants. Plots were separated by a row of red lettuce for clear visual distinction and automated image processing. The entire field was imaged over the course of one month on 10 non-consecutive dates: 2021-05-21, 2021-05-28, 2021-06-01, 2021-06-04, 2021-06-08, 2021-06-11, 2021-06-15, 2021-06-18, 2021-06-22,2021-06-25. This was done by the Netherlands Plant Eco-phenotyping Centre (NPEC) using a manned imaging robot “TraitSeeker” (https://www.npec.nl/tool/traitseeker-and-uavs/) over the plots. For more information see also Dijkhuizen et al. 2025 [14]. During the same period, hourly weather data was measured with an on-site weather station, which recorded air and soil temperature in °C, rainfall in mm and vapor pressure deficit in kPa.

### Data acquisitions and pre-processing

The TraitSeeker was equipped with sensors that measure hyperspectral reflectance and height profiles at the same time. The images were taken within a closed box equipped with Tungsten halogen lamps for controlled lighting conditions and removal of ambient daylight. The hyperspectral images were taken by two spectral cameras that cover the visible and near infrared range (VNIR) (397.66nm - 1003.81nm, stepsize: 2.62nm, Specim FX10, SPECIM, Oulu, Finland) and the shortwave infrared range (SWIR) (935.61nm-1720.23nm, stepsize:3.58nm, Specim FX17, SPECIM, Oulu, Finland) of the electromagnetic spectrum. Height images were taken with a 2D LIDAR sensor (Sick Ranger, St Minneapolis, MN, USA) calibrated for each spectral camera such that each spectral image and corresponding height image of a plot can be combined. The raw data collected by the TraitSeeker was pre-processed by NPEC with an in-house pipeline. This pipeline segments spectral and height images per spectral camera for each timepoint and plot in the field. Each pixel in an image corresponds with the reflected light intensity for each nm band and a height value. In addition, NDVI images were generated per plot, where each pixel corresponded to the calculated NDVI value from spectral bands taken by the FX10 camera. For each timepoint, daily weather data were aggregated by calculating mean soil and air temperature, mean vapor pressure deficit, and total rainfall (**Table S3**).

### Image processing

Spectral data was extracted for VNIR (FX10) and NIR (FX17) images separately. Spectral data between cameras could not be directly combined due to different resolutions and dimensions of both cameras. Per plot, two data cubes were collected, one per camera. All image processing was performed in R (Version 4.4.1) [19]. NDVI and height images were loaded into R using Imager (Version 1.0.2) [20]. Per image, pixels were first classified as either belonging to soil or plant using the corresponding spectral indices and height pixel values. For FX10 images, pixels with a NDVI pixel value > 0.6 and a height pixel value of > 0.02 were classified as “plant”. For FX17, 3-channel greyscale images were created using the log2-transformed ratio of pixel values at wavelengths of 967nm and 1457nm for all channels. Fake RGB images were converted to greyscale with increased contrast using the command RGB2gray from R-package SpatialPack, with method= “weighted” and weights = c(5,-5,10) [21]. Using this method, soil pixels appear much darker than plant pixels (**Figure S1+S2**). Spectral images were loaded using caTools (Version 1.18.2) [22]. Using the coordinates of all plant pixels in a plot, the mean plant height and mean reflectance per nm step was determined using the mean() function in R. Per plot this produced a representation of the mean reflectance spectrum across VNIR and NIR for a given accession. This dataset provided the basis for all further phenotyping.

### Spectral phenotyping

We defined the averaged plant pixel height and averaged plant pixel reflectance per wavelength as the most basal phenotypes for each replicate. The measured spectra from the FX10 and FX17 cameras overlapped between 935 nm and 1003 nm. To avoid redundancy, we used the FX10 data and excluded the corresponding measurements from the FX17 camera, as overlapping spectra showed strong correlation between cameras (**Figure S3**). To identify outliers, we plotted the full reflectance spectrum for each replicate and timepoint and flagged any replicates whose spectral curves deviated from the typical vegetation reflectance pattern (**Figure S4A**). Before downstream analyses, including analysis of variance, vegetation index (VI) calculation, and genome-wide association studies (GWAS), we applied a Standard Normal Variate (SNV) transformation on the spectral data. SNV is used to reduce variability in spectral reflectance due to scattering and spectral pathlength, allowing comparison of spectral curves between accessions [23]. SNV transformation was performed on the mean reflectance per wavelength per replicate per timepoint per camera. The mean reflectance of a given wavelength is subtracted with the mean reflectance of all wavelengths measured for that replicate. This is then divided by the standard deviation of all wavelengths. Outlier values were removed if the spectral shape before and after transformation did not follow the expected shape (**Figure S4B**). These extreme outliers were only observed in the FX17 camera data, so we limited outlier removal to that dataset (**Figure S4C**). For all further analyses using SNV-transformed spectral values, a constant of +5 was added to shift the data and remove negative values.

VI were calculated from SNV transformed wavelength values using the Index Data Base (IDB) (**Table S2**) [24]. To calculate a given VI, the closest (in nm) wavelength measured by either camera was used for the required wavelength of a VI. Heatmaps were created using the R-package ComplexHeatmaps (Version 2.14.0) [25].

### Filtering of red and green accessions

To separate red and green accessions using hyperspectral data, we defined a wavelength ratio that can be used for thresholding. The ratio was computed per replicate per timepoint as such:

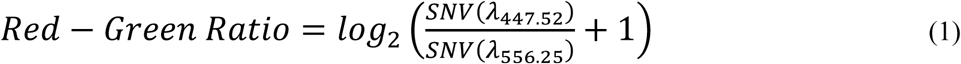

where *SNV*(*λ*) is the standard normal variate–transformed mean reflectance for a given genotype at wavelength *λ*; *λ*_447.52_ corresponds to reflectance at 447.52 nm near the chlorophyll b absorption peak in the blue region, where reflectance is typically lower for both red and green plants; and *λ*_556.25_ corresponds to reflectance at 556.25 nm near the green reflectance peak, which is high in green plants but reduced in red plants due to anthocyanin absorption. A pseudo count of +1 was added before logarithmic transformation to avoid undefined values. This ratio utilizes the distinct spectral shape of red versus green plants, where red accessions will show a higher Red-Green Ratio compared to green accessions, as their reflectance of green light is lower. We determined the threshold to classify green and red accessions by visually inspecting ranked Red-Green Ratio values across all sampling points. A conservative threshold of Red-Green Ratio < 0.95 was selected just before the inflection point, classifying accessions as green (**Figure S5**).

### Dimensionality reduction of wavelength phenotypes

To explore patterns of variation from both spectral and biological perspectives, we conducted principal component analysis (PCA) on wavelength phenotypes across accessions, and reciprocally, on accessions across wavelength phenotypes. For PCA on wavelength phenotypes we used the prcomp() function on SNV transformed wavelengths, which were first averaged across the two replicate per genotype, with scale=TRUE. For PCA on accessions, we first scaled the averaged SNV-transformed values using the scale function (center = TRUE, scale = TRUE) to correct for differences in relative brightness across wavelengths. No additional scaling was applied within the prcomp() function.

### Analysis of variance of wavelength phenotypes

To determine the variance contributed by replicate-plots on the field to total spectral variance, we modeled SNV-transformed reflectance at each wavelength using a linear mixed-effects model, with genotype, and replicate nested within genotype as random effects:

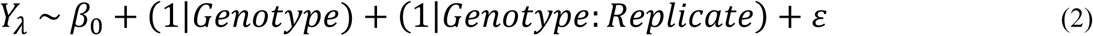

where *Y*_*λ*_ stands for the reflectance at a given wavelength *λ*, *β*_0_is the fixed intercept, (1|*Genotype*) is the random effect for genotype, (1|*Genotype*: *Replicate*) is the random effect of replicate nested within genotype, and *ɛ* is the residual error. The lmer() function of the lme4 R-package (Version 1.1-37) was used to fit the model [26]. The VarCorr() function of the lme4 R-package was used to estimate proportion of variance explained by genotype and replicate, as the fraction of total variance. Total variance was estimated as the sum of variances of all variance components, i.e. genotype, replicate and residual variance.

To understand how genotype, time and weather influence the spectral shape of lettuce accessions we fitted linear mixed models (LMMs) for each wavelength using replicate-level SNV transformed reflectance. The lmer() function of the lme4 package (Version 1.1-37) was used, modelling reflectance as the outcome with time, air temperature, and rainfall as fixed effects and genotype as a random effect:

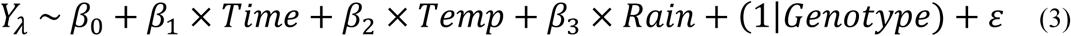

where *Y*_*λ*_ stands for the reflectance at a given wavelength *λ*, *β*_0_ is the fixed intercept, *β*_1_, *β*_2_, *β*_3_ are the fixed effect coefficients of *Time* (scaled measurement day), *Temp* (scaled average air temperature of that day), and *Rain* (scaled cumulative rainfall of that day), (1|*Genotype*) is the random effect of genotype, and *ɛ* is the residual error. We evaluated model performance using marginal and conditional R² values, where marginal R² represents the proportion of variance explained by fixed effects, and conditional R² represents the variance explained by both fixed and random effects. We used the r2_nakagawa() function of the performance R-package (Version 0.14.0), with ci=TRUE, ci_method=”boot” and iterations=100, to estimate confidence intervals [27].

To estimate the contribution of single fixed effects, namely time, temperature and rainfall per wavelength, we performed marginal effect prediction. For each fixed effect of a fitted model, we set all other fixed effects constant at 0, and computed variance of the newly formulated model, using the predict() and var() function of base R. We computed the total variance as the sum of variances of genotype, residual and the estimated variances from partitioning. We defined the contribution of each variance component, fixed, random and residual as their fraction from the total variance.

### Estimation of genotype-by-Environment BLUPs

To quantify phenotypic plasticity of individual lettuce accessions, genotype-by-environment-specific best linear unbiased predictors (BLUPs) were estimated from linear mixed effect models, as described by Arnold et al. (2019) [28]. Using lme4, separate models were fitted for each wavelength or specific VIs, with scaled time, mean air temperature, and rainfall as fixed effects, and genotype as random effect. To capture plasticity, each environmental variable was added as a random slope respectively:

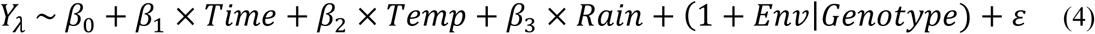

Where *Y*_*λ*_ stands for the reflectance at a given wavelength *λ*, *β*_0_ is the fixed intercept, *β*_1_, *β*_2_, *β*_3_ are the fixed effect coefficients of *Time* (scaled measurement day), *Temp* (scaled average air temperature of that day), and *Rain* (scaled cumulative rainfall of that day), (1 + *Env*|*Genotype*) is the random effect of genotype with a random slope *Env* for either environmental variable, and *ɛ* is the residual error. BLUPs for genotype-by-environment slopes were only extracted for wavelengths where the models converged successfully without singular fits or zero variance components, ensuring reliable variance component estimation. BLUPs were extracted from the fitted models using the ranef() function of lme4. These BLUPs represent genotype-specific sensitivity of reflectance to the respective environmental variable.

### Heritability estimation

Broad-sense heritability *H*^2^ of wavelength and VI phenotypes was estimated using two types of mixed-effect models, a base model defined in Equation 3, and an advanced model defined in Equation 4 with *Env* being *Time*. The advanced model was chosen to better account for genotype-by-time spectral variance, such that the genotypic variance component was estimated per timepoint. Per phenotype performance between the base and advanced models were compared using a likelihood ratio test with the anova() R-function, and the Akaike Information Criterion (AIC) using the AIC() R-function. The complex model was chosen for heritability estimate if the model converged successfully, the Benjamini-Hochberg adjusted p-value of the likelihood test was p < 0.05 and the AIC was lower than the base model. For the complex model, *H*^2^was estimated with the following equation:

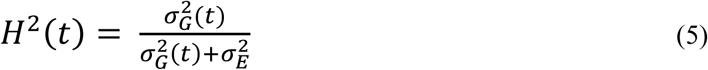

Where 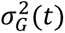 is the genetic variance per timepoint *t*, and 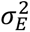 is the residual variance, including environmental and random. For the base model, the same equation is used, but genetic variance is estimated as constant across timepoints.

### SNP matrix

The SNP data for *L. sativa* accessions included in this study was previously published by Dijkhuizen et al. (2024), where the methodology for sequencing, SNP calling and SNP filtering is described [14, 29]. In short, SNPs were called against the Salinas V8 Lettuce reference genome, downloaded from NCBI (GCF_002870075.3) using the GATK HaplotypeCaller embedded in the Nextflow Illumina Analysis Pipeline (https://github.com/UMCUGenetics/NF-IAP/) [30]. Only bi-allelic SNPs were kept for further analysis. SNPs were clumped in batches of 2000 neighboring SNPs, with a clumping criterion of differing in less than 5 accessions. The SNP with the highest MAF was chosen as the SNP clump representative. SNP alleles were coded with 0 for homozygous reference allele, 1 for heterozygous and 2 for homozygous alternative allele.

### GWAS

Per phenotype SNPs with a MAF < 5% were excluded. GWAS was performed in R (Version 4.2.2) using the R-packages lme4QTL and pbmcapply, and GNU parallel [31–33]. *L. sativa* accessions LK087, LK196, LK197, LK198, LK199, LK200 were excluded from GWAS to exclude possible hybrids with wild species and oilseed varieties. GWAS was performed as described by Mehrem et al. (2024) [34]. A linear mixed model was used (relmatLmer and matlm) with the kinship matrix set as a random effect. Kinship was calculated per species as the covariance matrix of the respective SNP set using the cov() fuction in R. For all GWAS the Bonferroni method was used to correct for multiple testing (0.05/Number of tested SNPs), where SNPs that were excluded in the correlation-based filtering were included in the total number of tested SNPs. GWAS results were processed per phenotype and timepoint, generating plots and summary statistics tables automatically. Across GWAS results, QTLs were defined as genomic intervals spanning consecutive significant SNPs that were no more than 3 Mbp apart, with the peak position corresponding to the SNP with the lowest p-value within the interval. Genes falling within QTL boundaries were considered candidate genes. Candidates were investigated using the genome annotation by Van Workum et al. (2024) [29, 35]. For selected phenotypes, an iterative GWAS approach was used to dissect multiple QTLs, as described in Mehrem et al. (2025) [36].

### Data and code availability

Scripts used in this study can be found at https://github.com/SnoekLab/Hyperspec_Mehrem_etal_2025. The data is available upon request.

## Results

### Spectral- and Vegetation Index-Based phenotyping of *Lactuca sativa* in the field

To characterize phenotypic variation in *Lactuca sativa* under field conditions, we collected hyperspectral images of 194 accessions at 10 timepoints across the growing season. Images were captured at the plot level, with each plot containing 30-40 plants of the same accession, and two replicate plots were included per accession. The field layout is described in more detail by Dijkhuizen *et al.* (2025) [14]. This resulted in a high-dimensional dataset including reflectance at individual wavelengths, canopy height, and vegetation indices (VIs) (**Table S4**). The hyperspectral sensor captured the visible, near-infrared (VNIR), and shortwave infrared (SWIR) regions of the light spectrum, ranging from 380 to 1700 nm, in approximately 2 nm increments, resulting in 428 single-wavelength traits per plot. We examined this dataset across different dimensions. Comparing many accessions at a single timepoint revealed between-accession differences (**Figure 1A**), while tracking a single accession over time revealed temporal dynamics in spectral traits (**Figure 1B**). Across accessions and timepoints, we observed variation in reflectance patterns, including shifts in the red edge (680 to 730 nm), associated with chlorophyll content and canopy structure, and differences around 550 nm, where red lettuce varieties showed lower reflectance due to anthocyanin pigmentation (**Figure 1A, S6+S7**).

**Figure 1:**
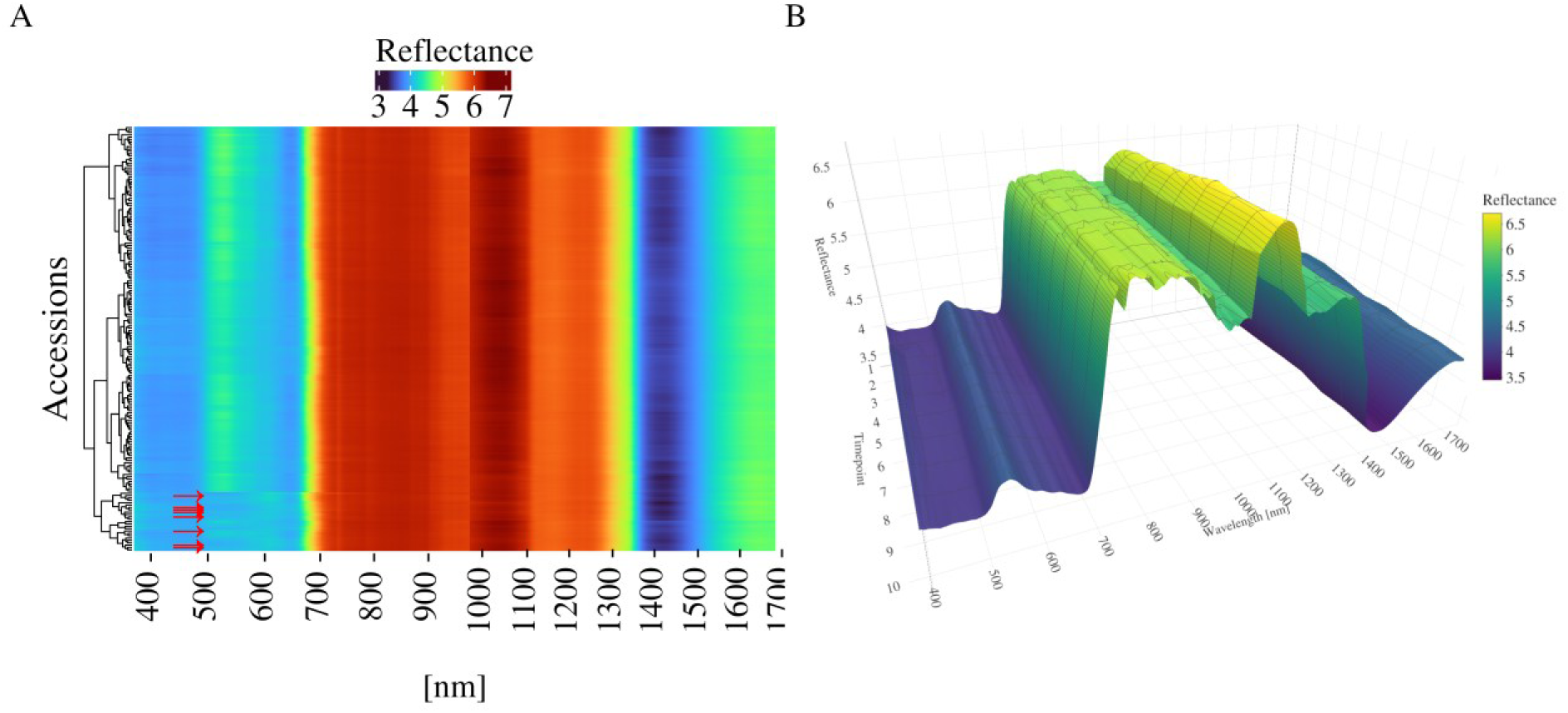
Spectral phenotypes of a single timepoint and single lettuce accession. **A:** Heatmap of averaged reflectance across measured wavelengths for all accessions for the first timepoint. Color indicates the transformed reflectance value. Accessions (rows) were clustered using the cluster_rows=T command of the Heatmap() function. Red arrows indicate the 10 most red accessions based on the Red-Green Ratio. **B**: 3D surface plot showing the spectral shape of one accession (LK123) across all timepoints. Color indicates transformed reflectance value.

To complement single-wavelength phenotypes, we calculated 135 different VIs at each timepoint, capturing a broad range of physiological and biochemical traits derived from combinations of multiple wavelengths. Most VIs were related to chlorophyll content and general vegetation detection (n = 89), with additional indices estimating water content (n = 13), water stress (n = 8), red-edge position and slope (n = 14), vegetation stress (n = 6), carotenoid content (n = 3), anthocyanin content (n = 1), and nitrogen content (n = 1).

Vegetation indices such as NDWI (water content), SIPI (carotenoid to chlorophyll ratio), ARI (anthocyanin accumulation), and height showed dynamic patterns across time, reflecting both developmental progression and environmental responses (**Figure 2A–D**) [37–39]. The weather conditions during the trial were characterized by low rainfall together with periods of elevated temperatures, resulting in a high vapor pressure deficit (VPD) (**Figure 2E**). NDWI gradually increased over time but plateaued between timepoints 4 and 6, increased again between timepoints 6 and 7, and then remained stable until timepoint 10, indicating reduced water availability or uptake (**Figure 2B**). Increases in ARI and SIPI suggested shifts in stress-related pigment accumulation that coincided with periods of elevated temperature and low rainfall, particularly at timepoints 4 to 8 (**Figure 2C-E**). Irrigation was limited, with a substantial watering event via sprinklers recorded between timepoints 7 and 8. These conditions likely imposed moderate to significant drought and heat stress on the plants.

**Figure 2:**
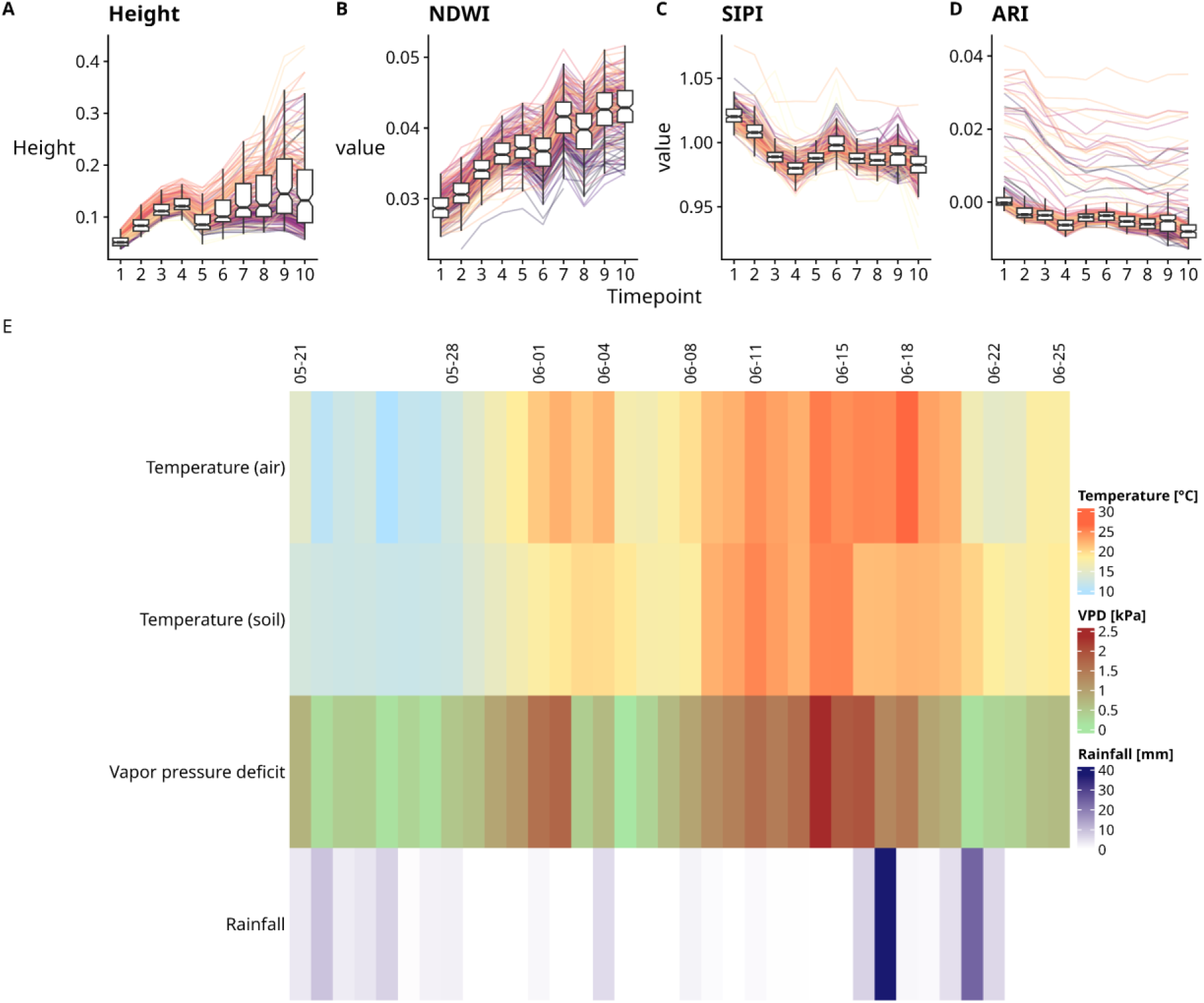
Temporal dynamics of selected spectral traits across *Lactuca sativa* accessions. **A**: Canopy height measured as LIDAR-derived pixel intensity values. **B**: Normalized Difference Water Index (NDWI). **C**: Structure Insensitive Pigment Index (SIPI) per accession. **D**: Anthocyanin Reflectance Index (ARI). Each panel shows a time series of mean trait values per accession, visualized as line plots overlaid by boxplots summarizing the distribution per timepoint. Boxplots represent interquartile range and median; outliers are not shown in boxplots but are reflected in the individual trajectories. **E**: Daily averages of air temperature, soil temperature in °C, and vapor pressure deficit (VPD) in kPa, along with cumulative rainfall in mm between the first and last timepoint. Columns represent days, timepoints are marked by dates on top. Note that the high rainfall on 17-06 is due to additional watering.

### PCA reveals major axis of spectral variation and wavelength-specific contributions

Spectral phenotypes capture known variation, such as morphological differences in horticultural type, pigment levels, as well as other unknown underlying patterns. We used principal component analysis (PCA) to identify major patterns of variation in the hyperspectral data of each timepoint, revealing both known and novel phenotypic signatures embedded in the spectral phenotypes. Across all time points, the first two principal components (PCs) explained a substantial proportion of the spectral variance, between 75% and 80% (**Figure 3**). Despite this high proportion, there was limited dispersion of accessions along these PCs, suggesting that major components of spectral variation reflected common features, like overall reflectance intensity or color, rather than traits distinguishing individual genotypes. Variation captured by subsequent PCs was initially low but gradually increased over time, indicating that new sources of variation emerged as the plants developed (**Figure S8**). While overall separation was limited, accessions belonging to the same horticultural type clustered moderately, suggesting that spectral phenotypes captured broad morphological or physiological similarities associated with horticultural type.

**Figure 3:**
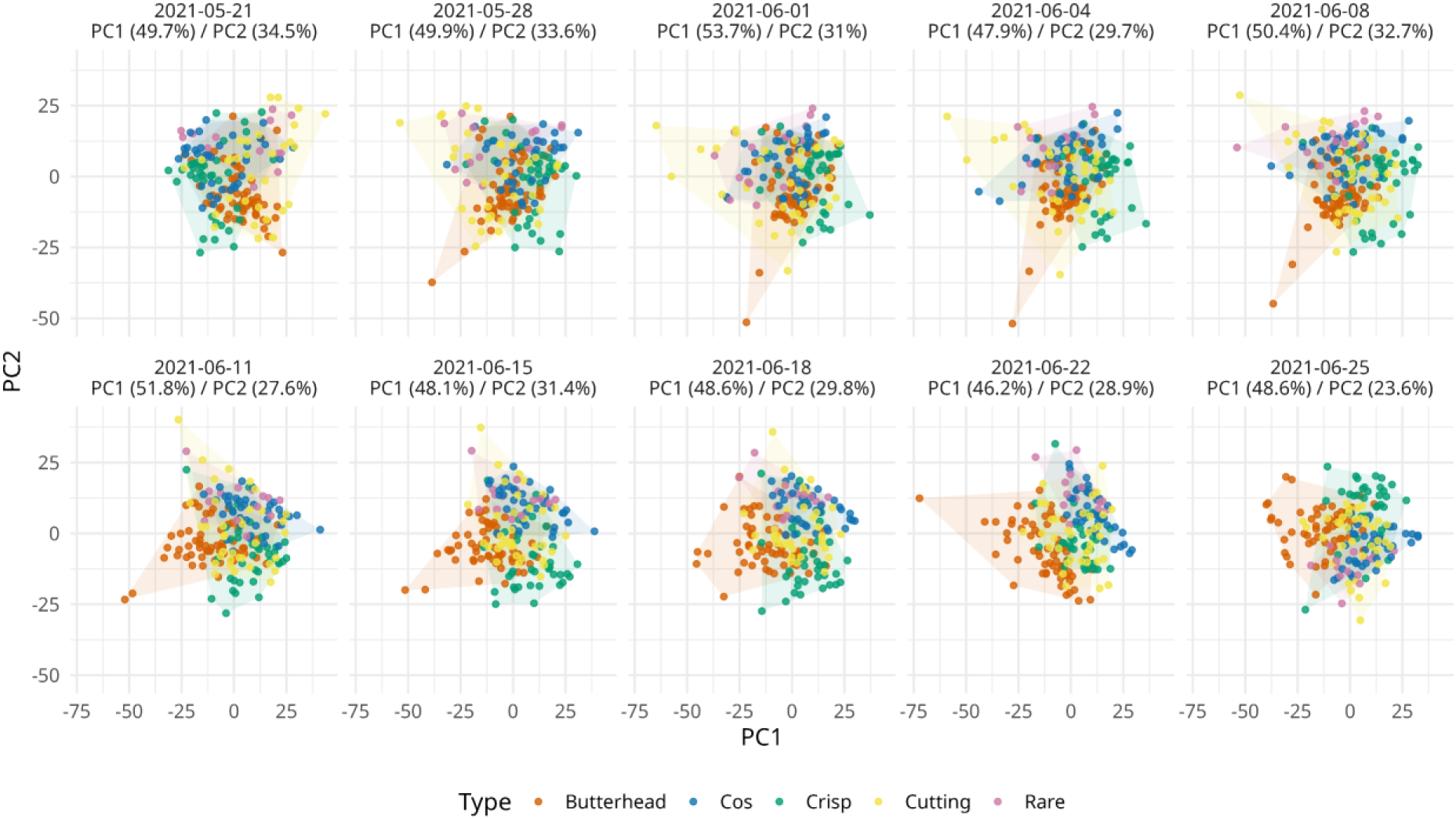
Principal component analysis (PCA) of spectral phenotypes across ten timepoints during the field trial. Each panel represents a separate timepoint, with PC1 and PC2 capturing most of the variation (percentage of variance explained indicated per plot). Points represent individual genotypes, colored by horticultural type: Butterhead (orange), Cos (blue), Crisp (green), Cutting (yellow), and Rare (purple). Hulls illustrate the spread of each type.

Accessions with high anthocyanin levels, and therefore red leaves, were clearly distinguishable based on their spectral profiles, suggesting a strong contribution to the variation captured by hyperspectral imaging (**Figure S7**). To evaluate the effect of red pigmentation, we repeated the analysis after filtering out red accessions using Red-Green Ratio thresholding applied to their spectral shape. To reduce the likelihood of still including strongly red accessions when repeating the PCA, we applied a conservative threshold that excluded some accessions that would be classified as green (**Figure S5**). Removal of red accessions weakly increased the variance explained of both PC1 and PC2 but did not change the overall outcome of the PCA which included the red accessions (**Figure S9+S10**). Red accessions likely contributed more to the variation captured by later principal components, but the primary pattern captured by the first two PCs did not appear to be dominated by anthocyanin pigmentation.

We examined the contribution of single wavelengths to the separation along the first two PCs over time using loadings, indicating which traits are captured by those PCs. Distinct peaks in absolute loading values across the VNIR and SWIR ranges highlighted spectral regions that contributed strongly to the explained variation (**Figure 4**). PC1 was driven by SWIR features beyond the red edge, with strong loadings near water absorption bands, pointing to traits like water content and cell structure. In the VNIR region, elevated loadings in the blue and green wavelengths likely reflect chlorophyll variation. Loadings in the visible range increased over time, even after excluding red accessions, while SWIR-associated peaks remained stable (**Figure S11**). PC2 was dominated by wavelengths in the blue and green regions of the visible spectrum, along with peaks near the red edge and the first water absorption band. Temporal dynamics in PC2 were more pronounced in the SWIR region, while loadings in the VNIR remained stable over time (**Figure 4 + S11**).

**Figure 4:**
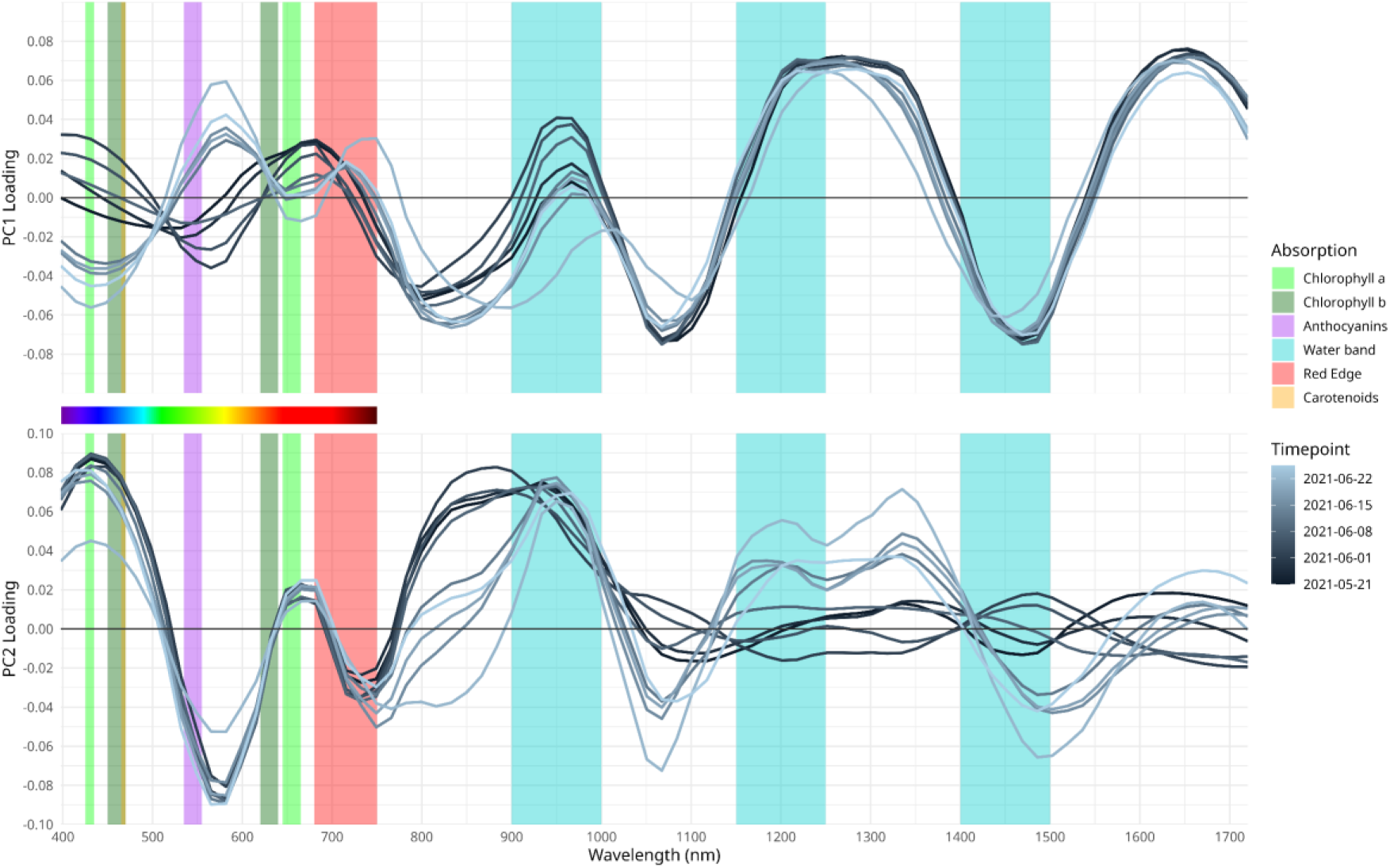
Loadings of wavelengths along the first two PCs over time. Loadings of PC1 (Top) and PC2 (Bottom) are plotted against the respective wavelength (x-axis) using a geom_smooth with the “loes” method with a span=0.2. Line color indicates time progression, with darker lines representing earlier timepoints and lighter lines later ones. Colored background bars indicate absorption spectra of known plant pigments and spectral features of plants. The visible light spectrum is shown as a color gradient between the panels to indicate approximate wavelength positions.

The effect of filtering out red accessions was most strongly observed in PC3, where wavelengths in the green and red region showed a high absolute loading before filtering, which was reduced after filtering, suggesting that anthocyanin content contributed considerably to the variation captured by this component (**Figure S12**).

### Reciprocal PCA illustrates growth-dependent spectral differences of horticultural type

We demonstrated that distinct spectral regions drove the observed spectral variation over time, and that horticultural type was clearly captured along the major axes of spectral variation. Reciprocal PCA, which swaps variables and samples to examine variation from the perspective of wavelengths, was applied per timepoint to assess how horticultural type drives spectral shape. Across PC1 and PC2, reciprocal PCA revealed a pattern of two angled perpendicular ring-like structures in principal component space (**Figure 11A+C**). One ring comprised primarily of VNIR-dominated observations, while the other reflected variation driven by SWIR wavelengths. The elliptical arrangement and clustering by wavelengths reflected strong internal correlations within each spectral region. Compared to the initial PCA, a similar overall pattern emerged with respect to cumulative variation explained over time, with the first two PCs capturing most variation, and later timepoints showing more variation in higher PCs (**Figure S13+S14**). These patterns were present regardless of filtering out red accessions (**Figure S15+S16**). While the initial PCA already indicated clustering by horticultural type, the reciprocal PCA vector fields provided a clearer visualization of how the first two major PCs capture these horticultural-specific reflectance patterns (**Figure 5B+D)**. This separation also revealed temporal dynamics, with horticultural types becoming more distinct as plant development progressed (**Figure S13+S14**). Vector fields showed that Butterhead accessions primarily separated along PC1, followed by Cos, Rare, and Crisp varieties, which separated along PC2. Cutting varieties did not show clear separation along the first two major PCs, consistent with their minimal dispersion observed in the initial PCA (**Figure 3**, **Figure 5 B+D**). Along subsequent PCs, horticultural types showed variable separation across timepoints (**Fig S17+S18**).

**Figure 5:**
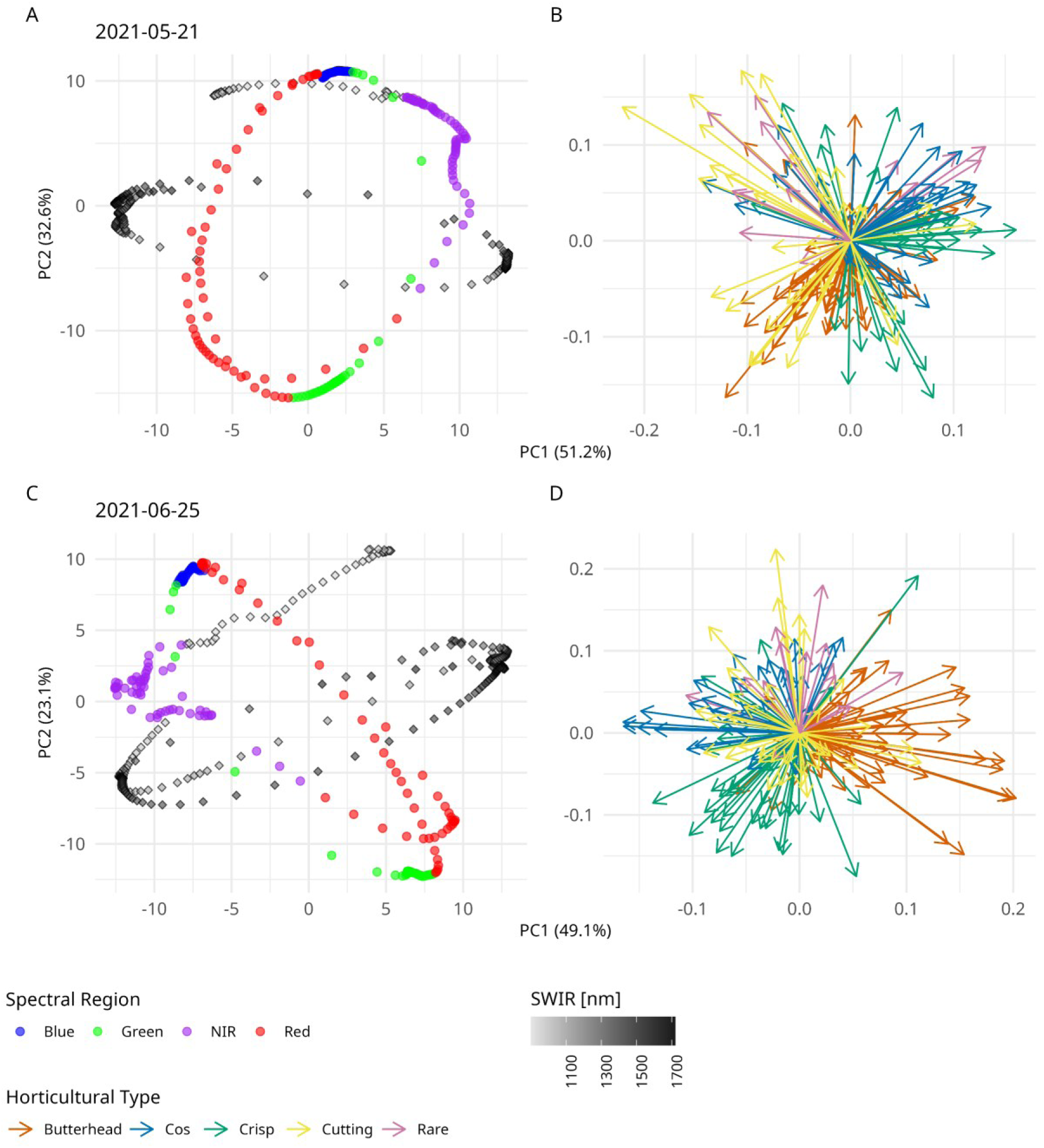
Reciprocal PCA plots of spectral phenotypes for all *>L. sativa* accessions of the first and last timepoint. PCA results based on the first two principal components are shown for the first (A) and last (C) timepoint, with the corresponding vector field (B: First timepoint, D: Last timepoint). Axes represent the first (x-axis) and second (y-axis) principal components, with the percentage of variance explained indicated in parentheses. Data points of the PCA are color-coded by spectral region (Blue, Green, Red, purple:VNIR, grey to black: SWIR). Vectors are colored by horticultural type: Butterhead (orange), Cos (blue), Crisp (green), Cutting (yellow), and Rare (purple).

### Estimating genotypic, temporal, and environmental effects on spectral variation

Dimensionality reduction revealed temporal and horticultural type variations in spectral phenotypes. Variation in reflectance was linked to prolonged high temperatures and low rainfall, conditions likely imposing drought and heat stress on plants. We used linear mixed modeling and variance decomposition to estimate the relative influence of genotype, time, and weather on spectral variation per wavelength. The full mixed-effects model included time, average air temperature, and cumulative rainfall per timepoint as fixed effects, and genotype as a random effect. Possible covariates of plot replicates and horticultural type were excluded from the model. Plot replicates did not show a strong contribution to the explained spectral variance (**Figure S19**). Although horticultural type accounted for significant spectral variance, it was omitted to avoid collinearity with genotype and confounding genotype-level estimates. Model performance (Nakagawa’s R²) varied across the VNIR and SWIR spectrum (**Figure S20**). Marginal R² values (fixed effects only) ranged from 0.031 to 0.777 (mean 0.391 ± 0.256), while conditional R² values (fixed + random effects) ranged from 0.301 to 0.917 (mean 0.755 ± 0.118), highlighting the substantial contribution of genotype (random effect) to overall spectral variation (**Table S5**). Decomposition of variance components showed that genotype and time contributed more strongly than temperature and rainfall (**Figure 6**). Residual variance within the model was consistently present across all wavelengths, representing variance captured by the model but not attributable to individual fixed or random effects. The influence of time was particularly pronounced in the SWIR region, possibly capturing changes in plant structure and water content during development. Air temperature explained more variance than rainfall and showed variable importance across the spectrum. In the visible range (450–600 nm), associated with pigments like chlorophylls and anthocyanins, the model explained a substantial portion of variance. In contrast, performance declined around 760 nm, just beyond the red edge, and near 1100 nm, prior to a water absorption band, indicating lower explanatory power in these regions (**Figure S20**). Interestingly, although the visible spectrum showed lower marginal R² values, implying a stronger influence of genotype, an exception occurred around 510 nm, where fixed effects explained over 60% of the variance (**Figure 6**). This sharp increase was primarily driven by temporal change, suggesting that 510 nm captures dynamic shifts in pigment content, possibly linked to greenness or chlorophyll degradation.

**Figure 6:**
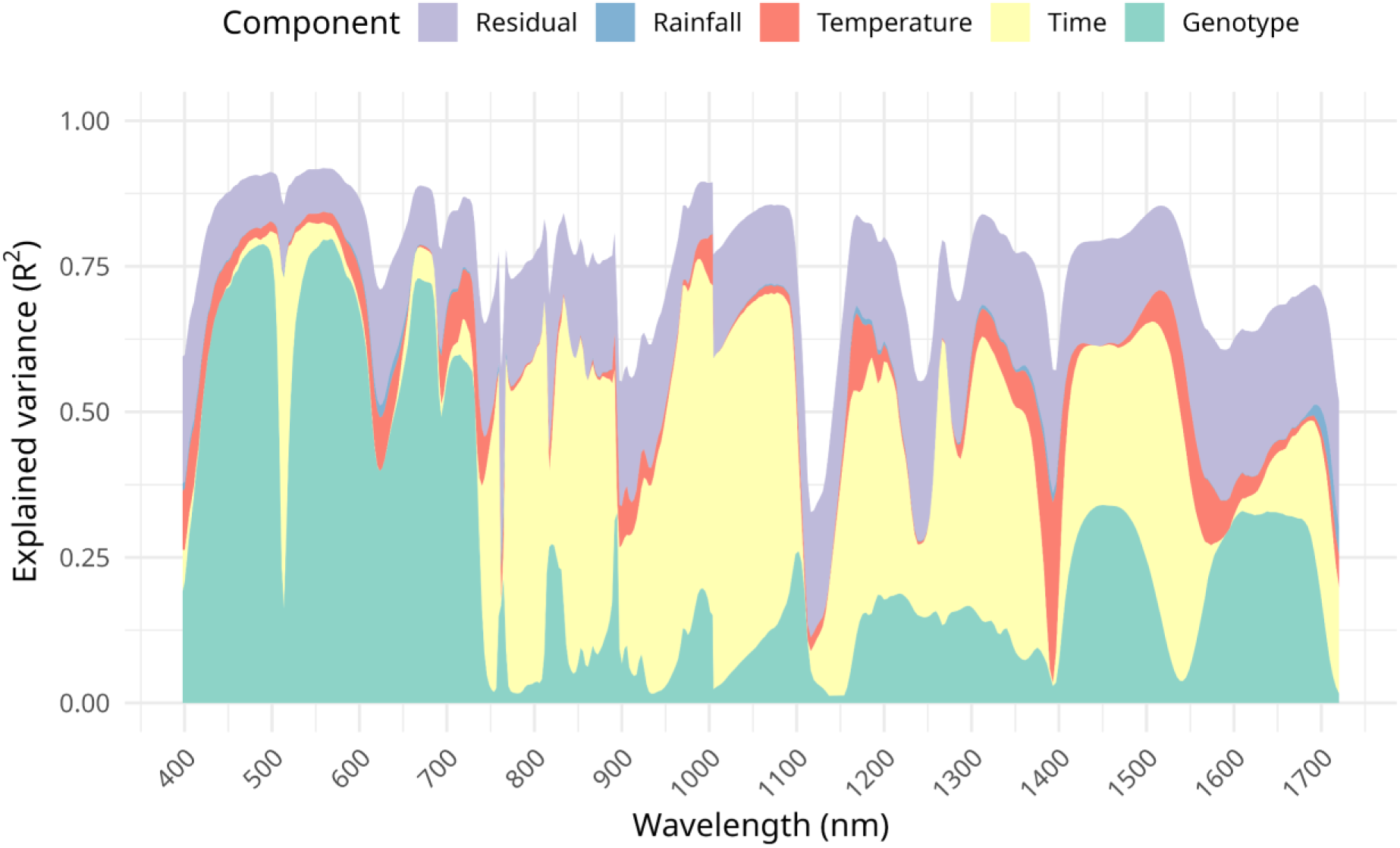
Decomposition of variance components from the full mixed-effects model across wavelengths. The stacked area plot shows the proportion of explained variance (R^2^) attributed to each component: Genotype (green), time (yellow), temperature (red), rainfall (blue), and residual variance of the model (purple), across the spectral range. The residual component reflects variation not captured by the fixed or random effects but still explained by the model. The space above the stacked area represents unexplained variance beyond the model.

### Estimating phenotypic plasticity using best linear unbiased prediction

Reflectance at different wavelengths is influenced variably by genotype, time, and weather, as demonstrated previously. Non-genetic factors contribute to phenotypic variation and can also reveal genotype-specific responses, a property known as phenotypic plasticity. To assess this plasticity in lettuce, we estimated best linear unbiased predictors (BLUPs) in relation to time, temperature, and rainfall. These BLUPs reflect the genotype-specific sensitivity to a change in each environmental factor respectively. We assumed that each factor affected phenotypic variation at least separately and therefore modeled each as a random interaction with genotype. BLUPs were estimated only for wavelengths with stable model fits. To account for the strong spectral signal of red varieties, we repeated the analysis using only green accessions (**Figure S21**). Across VNIR and SWIR wavelengths, lettuce accessions showed diverse patterns of phenotypic plasticity, especially in response to time and temperature, reflecting genotype-specific developmental trajectories and responses to environmental cues (**Figure 7**). Phenotypic plasticity also varied across horticultural types. For instance, Oilseed, Stalk, and Cos types showed stronger temporal shifts in the 400-500 nm range, while Butterhead, Crisp, and Latin types exhibited weaker responses in this region (**Figure 7A**). In response to temperature, Oilseed accessions had elevated reflectance shifts to around 550 nm, and Stalk types responded more strongly near to 1450 nm, a known water absorption band (**Figure 7B**). Model fits for rainfall were unstable across all accessions. However, when restricting the analysis to green varieties, stable fits emerged at 11 wavelengths, primarily in the water absorption regions around 900 nm and 1400 nm (**Figure S21C**). Consequently, BLUPs for rainfall responses were estimated only in this subset.

**Figure 7:**
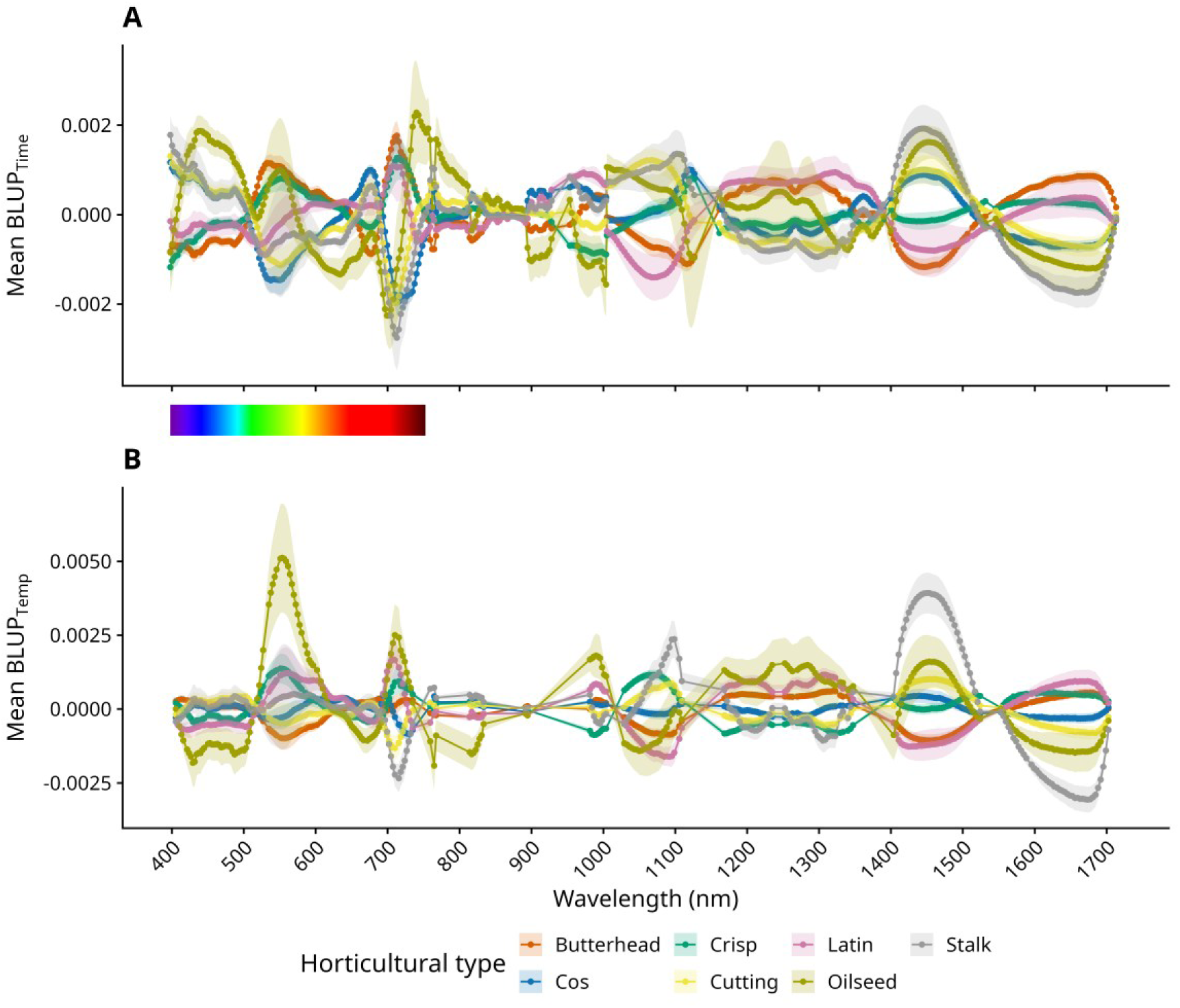
Mean BLUPs (Best Linear Unbiased Predictiors) across wavelengths for genotype-by-environment interaction components time and air temperature. BLUPs were extracted from linear mixed models with stable fits (x-axis), then rescaled to reflect the estimated change in reflectance per day (**A**) or per °C (**B**) (y-axis). Each colored line represents a different horticultural type of lettuce: Butterhead (red), Crisp (green), Latin (pink), Stalk (grey), Cos (blue), Cutting (yellow), and Oilseed (dark yellow), with shaded areas indicating standard deviation. The visible light spectrum is shown as a color gradient between the panels to indicate approximate wavelength positions.

### Estimating heritability of wavelength and vegetation index phenotypes over time

We estimated broad-sense heritability (H²) of wavelength and vegetation index traits to assess the total proportion of phenotypic variance attributable to genetic factors (**Table S6**). H² was calculated using a linear mixed model incorporating a random slope for time, enabling quantification of genetic variance while accounting for variable developmental changes over time. Patterns of H² across wavelengths revealed both stable and dynamic regions of heritability, broadly consistent with the genotype-driven variance estimated from the variance partitioning (**Figure 7+8**). Wavelengths in the visible range up to approximately 725 nm exhibited high and temporally stable H². In contrast, certain regions of the NIR and SWIR spectra, particularly from 900-950 nm and 1100-1150 nm, showed consistently low heritability. Additionally, H² shifted over time for specific regions, such as between 1200 and 1300 nm, suggesting growth-stage-dependent genetic control or changing environmental sensitivity. Vegetation indices, derived from combinations of wavelengths, similarly displayed a broad range of heritability patterns, both stable and dynamic over time (**Figure 9**). The anthocyanin index exhibited the highest H² throughout the trial, consistent with strong genetic control of red versus green pigmentation across varieties. In contrast, the heritability of plant height increased over time, reflecting variation in bolting or flowering time, developmentally timed traits with strong genetic regulation. Altogether, the high and dynamic heritability observed across spectral traits and VIs indicates a strong genetic signal.

**Figure 8:**
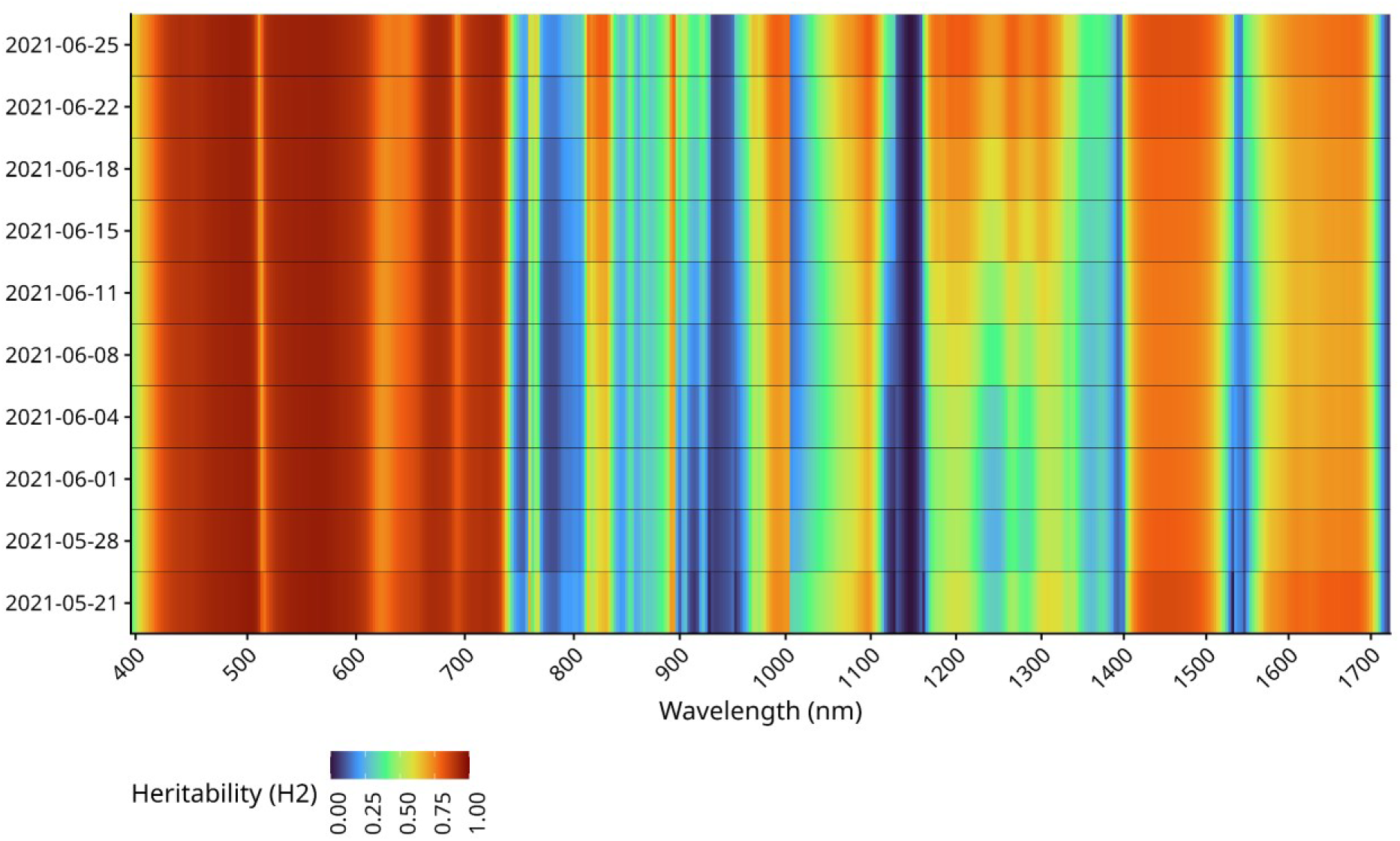
Temporal variation in spectral reflectance heritability (H^2^) across wavelengths. This heatmap illustrates the broad-sense heritability (H²) of spectral phenotypes (x-axis) per timepoint (y-axis) from all *L. sativa* accessions. The color gradient, from blue (low heritability) to red (high heritability), indicates the magnitude of genetic contribution to the observed spectral variation at each wavelength and timepoint.

**Figure 9:**
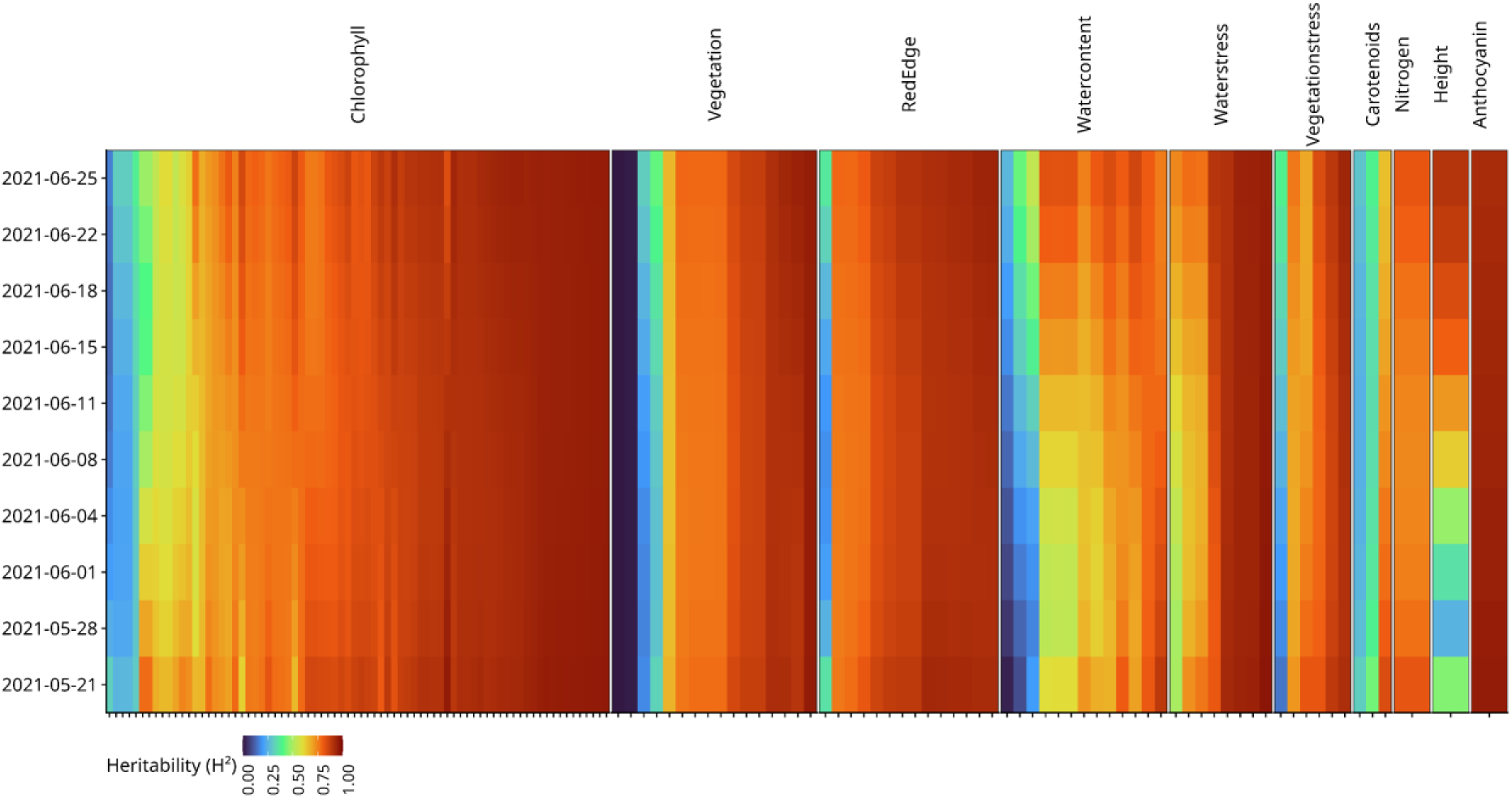
Temporal variation in spectral reflectance heritability (H2) across vegetation indices (VIs). This heatmap illustrates the broad-sense heritability (H²) of VI phenotypes (x-axis) per timepoint (y-axis) from all *L. sativa* accessions. The color gradient, from blue (low heritability) to red (high heritability), indicates the magnitude of genetic contribution to the observed spectral variation at each wavelength and timepoint. VIs are grouped by category.

### GWAS based on Principal Components over time

To investigate the genetic architecture underlying hyperspectral phenotypes, we performed GWAS on the first 10 PCs of spectral variation at each timepoint, using each accession’s PC scores as phenotypes (**Figure 3, Tables S7-S9**). This approach allowed a global scan for QTLs influencing dominant and minor axes of spectral variation and revealed both stable and transient QTLs with distinct temporal dynamics (**Figure 10**). A broad QTL on chromosome 3 (QTL_Chr3) was associated with PC1 of the first four timepoints, spanning 100.7 to 111.8Mbp and containing 178 candidate genes. Since PC1 was strongly influenced by SWIR wavelengths linked to water absorption, this QTL was could be associated with variation in water content of field-grown lettuce and linked to earlier developmental stages. For PCs 2 and 3, GWAS detected major QTLs on chromosomes 4 and 5, QTL_GLK and QTL_MYB, that co-localize with known regulators *LsGLK2* and *LsMYB113*, which control chlorophyll and anthocyanin variation in *L. sativa* [16, 18]. An additional QTL (QTL_ANS) on chromosome 9, mapping to *LsANS* and also linked to anthocyanin production in *L. sativa*, showed different behavior in green accessions [16]. It was captured by PC2 at later timepoints of all accessions but appeared in PC5 and PC6 during early timepoints of only green accessions.

**Figure 10:**
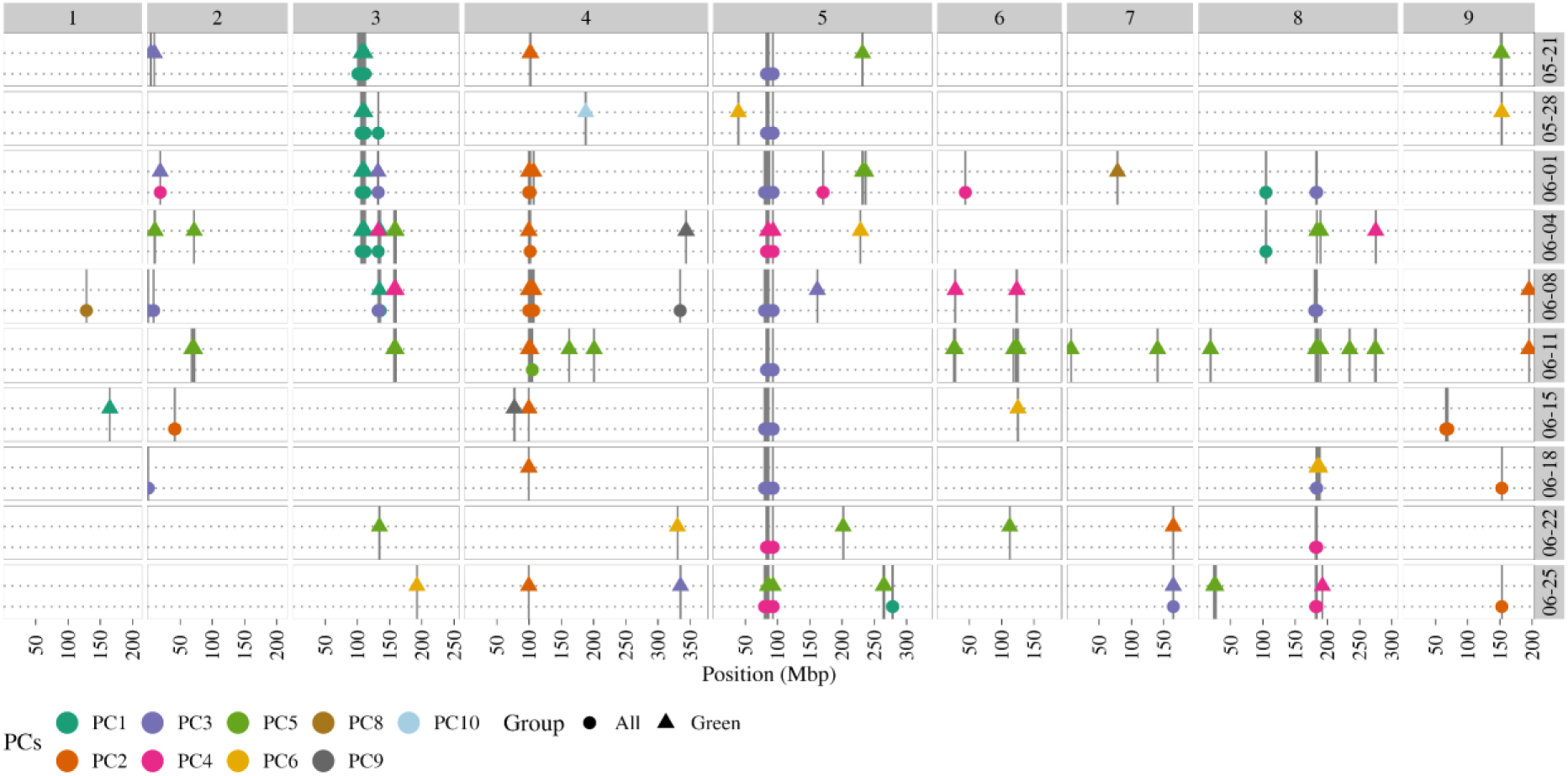
Multi-Manhattan plot of spectral principal components per timepoint. GWAS results for the first ten spectral principal components across timepoints. The x-axis shows genomic position in mega base pairs, with chromosome numbers indicated above. Significant QTLs (-log₁₀(P) > 7.69, Bonferroni-corrected) are plotted, and grouped by timepoint on the right. Points represent QTLs identified using all accessions, while triangles indicate QTLs detected in the subset of green accessions. Colors correspond to the principal component used as the phenotype. Vertical lines highlight the precise genomic position of each QTL within each timepoint.

Towards the final timepoints of the trial, PC2 and PC3 revealed a QTL on chromosome 7 (QTL_PHYC) mapping to *LsPhyC*, a key locus controlling flowering time in lettuce [16]. This shift highlighted that major axis of spectral variation increasingly reflected bolting or flowering-related traits as plants developed, as was reflected by time-driven differences in loadings of SWIR wavelengths (**Figure 4**).

Building on these PC-driven findings, we examined single-trait GWAS results for individual wavelengths and vegetation indices to validate the genetic signals identified through dimensionality reduction (**Tables S10+S11**). QTL hotspots from single-trait analyses overlapped with those detected via PCs, with similar temporal patterns and shared captured variation (**Figure S22**). Interestingly, a novel QTL was identified for PC9 at timepoint 06-15 in green accessions on chromosome 4, mapping to 76.3-77.6Mbp (**Figure 10, Table S9**). This QTL was not detected in any other GWAS. A single gene maps within this QTL, a VQ motif-containing protein. VQ proteins are transcriptional co-regulators that modulate responses to both biotic and abiotic stresses [40].

### GWAS on plastic phenotypes shaped by time and environment

To investigate the genetic basis of phenotypic plasticity, we performed GWAS with time, temperature and rainfall-linked BLUPs as phenotypes (**Tables S12-S14**). Between rainfall, temperature and time-linked BLUPs, temperature-linked BLUPs showed the clearest QTL-hotspots on chromosome 1 (QTL_1Temp), chromosome 5 (QTL_MYB), and chromosome 9 (QTL_ANS), and several additional QTLs (**Figure 11, Table S14**). QTL_1Temp was predominantly found in GWAS with all accessions and linked to BLUPs of wavelengths in the SWIR spectrum only, reflecting cell structure or water content properties. In single wavelength GWAS, this locus was linked specifically to timepoints 06-11 and 06-15, where plants underwent prolonged heat and drought conditions (**Figure S22**). The locus spanned from 57.2 to 58.6 Mbp, harboring 37 genes. A possible candidate gene is LOC111905038, a predicted vacuolar ATPase subunit A (VHA-A1). V-ATPases, including VHA-A1, support multiple stress-adaptive functions in plants, such as stabilizing cellular osmotic balance through vacuolar acidification. Under simultaneous heat stress and water deficit, this acidification could help cells maintain osmotic homeostasis [41].

**Figure 11:**
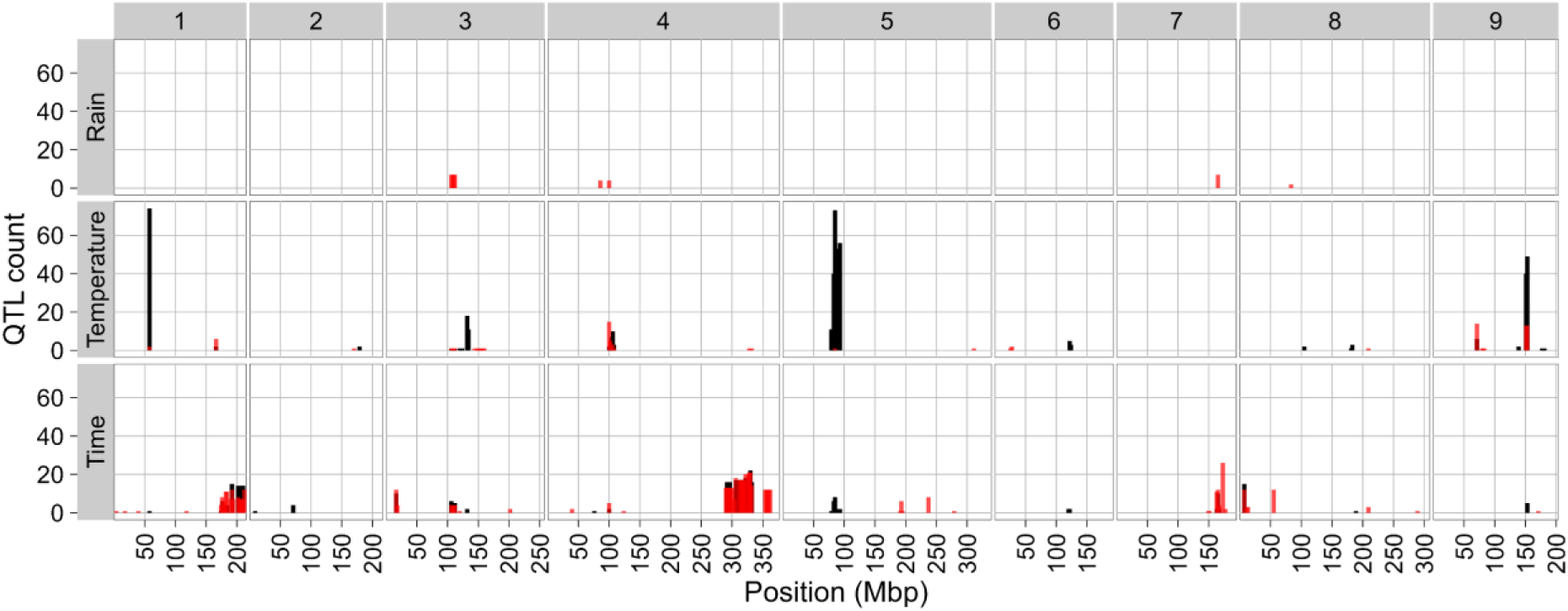
Histogram of QTLs found for time, rain and temperature-linked BLUPs. QTLs were defined as the most significant SNPs per 2 Mb window, per trait (–log₁₀P > 7.69, Bonferroni-corrected). The x-axis represents genomic position in mega base pairs, with chromosome numbers shown above, while the y-axis indicates the QTL count. Bar colors distinguish QTLs detected using all accessions (black) from those identified in green accessions only (red). QTLs are grouped by variance component, labeled on the left. Note that GWAS on rain-linked BLUPs was conducted exclusively for green *L. sativa* accessions.

GWAS on rain-linked BLUPs produced limited QTLs, identifying QTL_Chr3, QTL_GLK and QTL_PhyC, and additional minor QTLs (**Figure 11**, **Table S14**). GWAS on time-linked BLUPs identified several QTLs, with a broad region at the end of chromosome 1, spanning 173.9 to 211.3Mbp. This large QTL is exclusively found with time-linked wavelengths in the SWIR spectrum, specifically wavelengths between 970 to 1000nm, suggesting a link with temporal change in water content or cell architecture.

### Disentangling QTLs underlying plastic anthocyanin variation

Including red accessions in GWAS caused most associations to be dominated by major anthocyanin-associated QTLs (**Figure S22**). Meanwhile, anthocyanin content, estimated by the ARI, varied over time across all accessions, potentially influenced by heat stress or developmental timing (**Figure 2D**). To explore the genetics underlying these dynamics, GWAS was performed on ARI phenotypes across timepoints and on BLUPs of ARI to capture plastic responses, identifying multiple QTLs and candidate genes (**Table 1)**. GWAS including red accessions yielded static QTLs driven by major pigmentation loci, whereas excluding them uncovered transient QTLs, suggesting dynamic genetics underlying anthocyanin regulation (**Figure 12**+**S23**). In green accessions, QTL patterns shifted significantly after the third timepoint, with QTL_ANS absent until the final timepoint, while QTL_MYB60 on chromosome 2 and QTL_MYB113 became more prominent.

**Figure 12:**
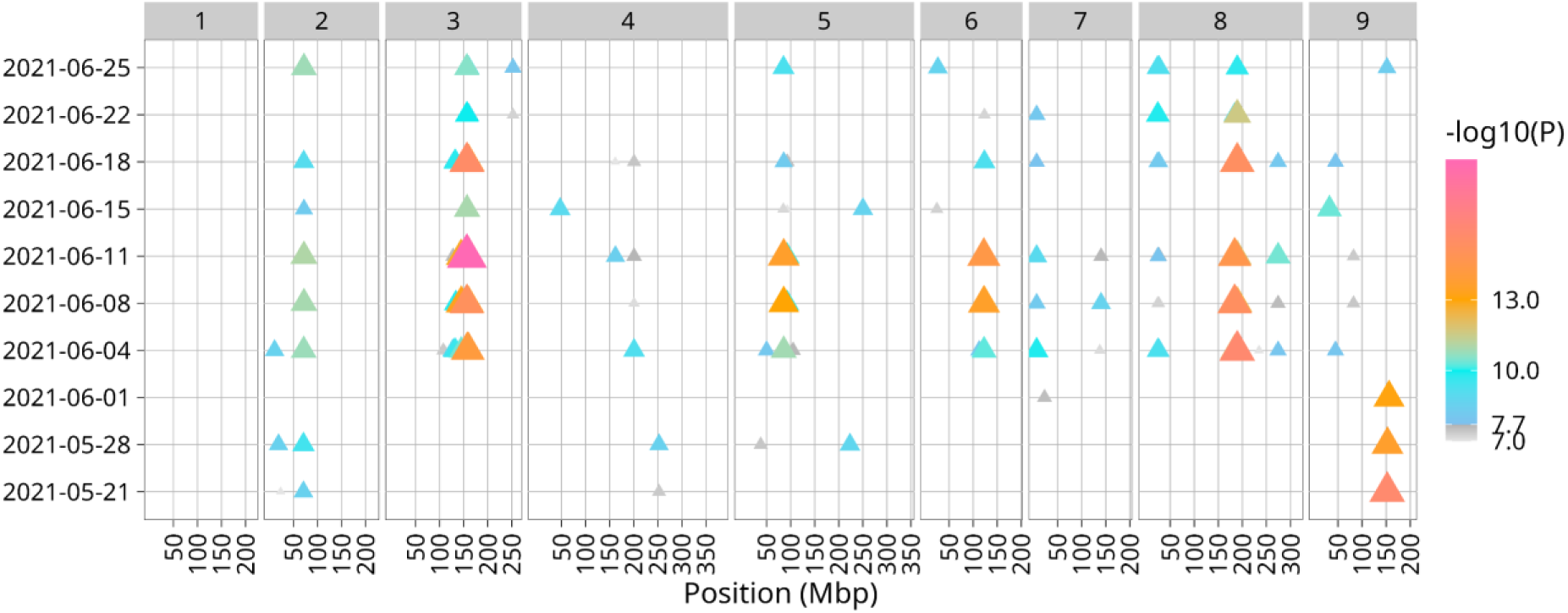
GWAS of anthocyanin reflectance index (ARI) in green *L. sativa* accessions over time. Multi-manhattan depicts QTLs associated with ARI variation per timepoint (left). The x-axis shows genomic position in mega base pairs, with chromosome numbers indicated above. Significant QTLs (–log₁₀(P) > 7.69, Bonferroni-corrected) are marked as triangles, where color and size correspond to the -log_10_(P) value.

**Table 1:**
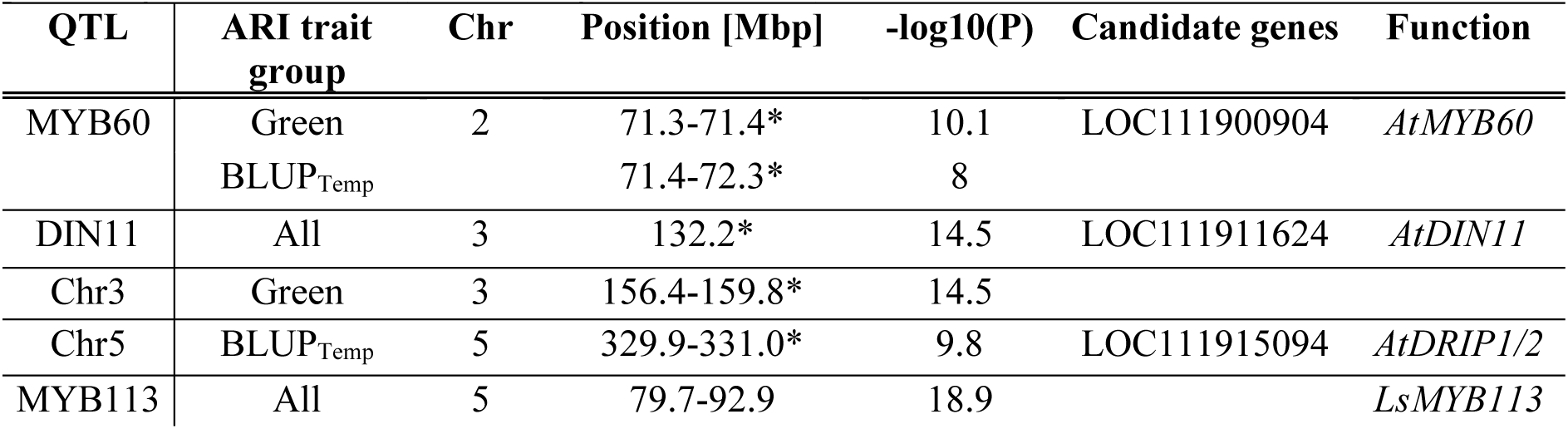
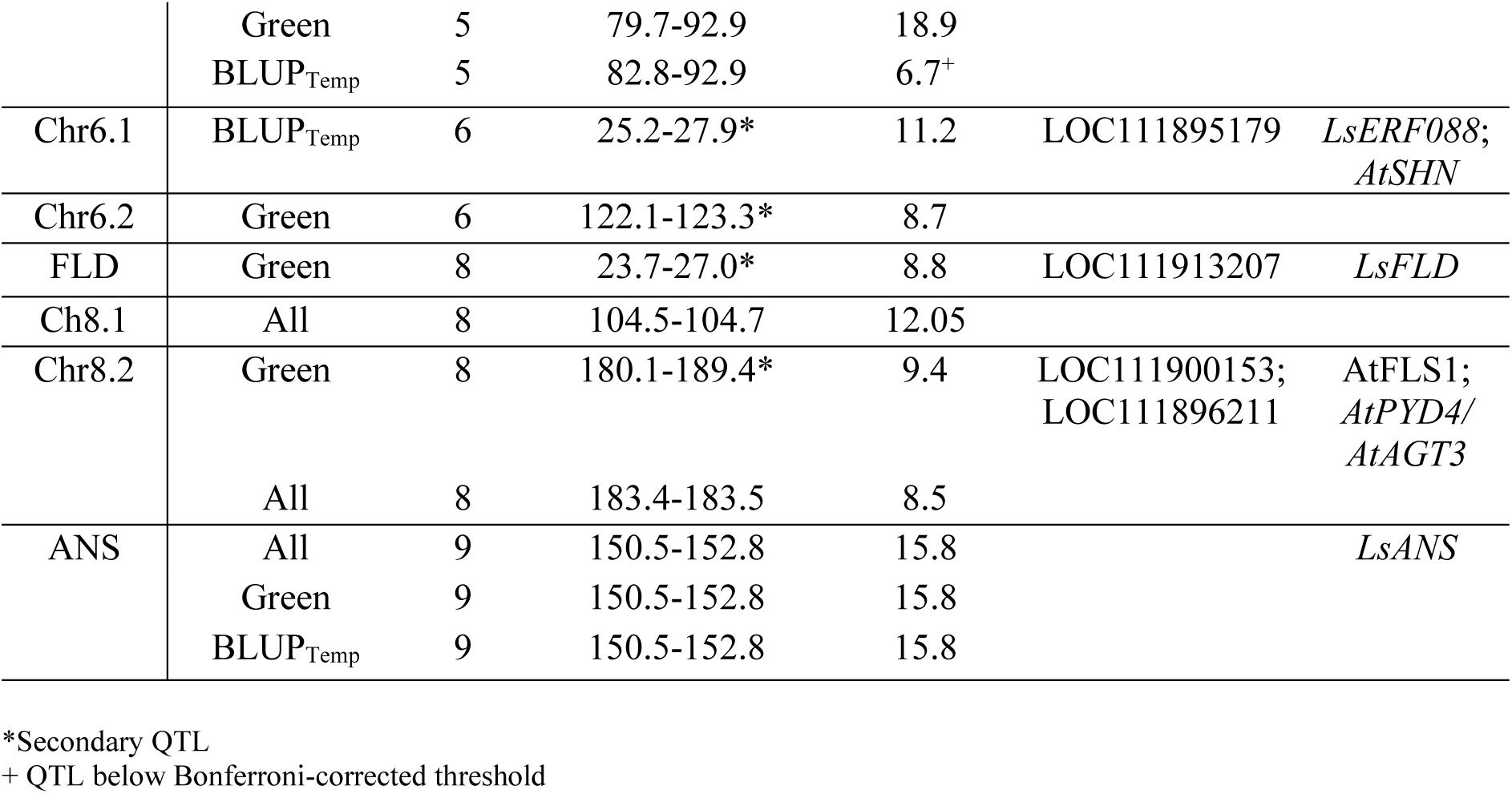
QTLs identified by GWAS on ARI. The table lists the name of the QTL (QTL), the GWAS it was detected in (ARI trait group), the genomic location (Chr, Position), the p-value (-log10(P)) and possible genes of interest (Candidate genes, Function).

| QTL | ARI trait group | Chr | Position [Mbp] | $-\log_{10}(P)$ | Candidate genes | Function |
| --- | --- | --- | --- | --- | --- | --- |
| MYB60 | Green | 2 | 71.3-71.4* | 10.1 | LOC111900904 | *AtMYB60* |
|  | BLUP<sub>Temp</sub> |  | 71.4-72.3* | 8 |  |  |
| DIN11 | All | 3 | 132.2* | 14.5 | LOC111911624 | *AtDIN11* |
| Chr3 | Green | 3 | 156.4-159.8* | 14.5 |  |  |
| Chr5 | BLUP<sub>Temp</sub> | 5 | 329.9-331.0* | 9.8 | LOC111915094 | *AtDRIP1/2* |
| MYB113 | All | 5 | 79.7-92.9 | 18.9 |  | *LsMYB113* |
|  | Green | 5 | 79.7-92.9 | 18.9 |  |  |
|  | BLUP<sub>Temp</sub> | 5 | 82.8-92.9 | 6.7<sup>+</sup> |  |  |
| Chr6.1 | BLUP<sub>Temp</sub> | 6 | 25.2-27.9* | 11.2 | LOC111895179 | *LsERF088*;<br>*AtSHN* |
| Chr6.2 | Green | 6 | 122.1-123.3* | 8.7 |  |  |
| FLD | Green | 8 | 23.7-27.0* | 8.8 | LOC111913207 | *LsFLD* |
| Ch8.1 | All | 8 | 104.5-104.7 | 12.05 |  |  |
| Chr8.2 | Green | 8 | 180.1-189.4* | 9.4 | LOC111900153;<br>LOC111896211 | *AtFLS1*;<br>*AtPYD4*/<br>*AtAGT3* |
|  | All | 8 | 183.4-183.5 | 8.5 |  |  |
| ANS | All | 9 | 150.5-152.8 | 15.8 |  | *LsANS* |
|  | Green | 9 | 150.5-152.8 | 15.8 |  |  |
|  | BLUP<sub>Temp</sub> | 9 | 150.5-152.8 | 15.8 |  |  |

\*Secondary QTL
+ QTL below Bonferroni-corrected threshold

With GWAS on BLUP_Time_ and BLUP_Temp_ phenotypes derived from the ARI of green varieties, we attempted to identify QTLs that are more likely explained by a developmental or environmental change. GWAS on BLUP_Time_ did not yield any significant QTLs, while GWAS on BLUP_Temp_ revealed several clear QTLs (**Figure 13, Figure S24**). Both QTL_ANS and QTL_MYB60 were significantly associated with temperature-linked variation of anthocyanin. Interestingly, GWAS on BLUP_Temp_ phenotypes of single wavelengths near the anthocyanin absorption peak (∼556 nm) also identified a strong association with QTL_ANS in green accessions, whereas simple mean reflectance values did not recover this signal (**Figure S25+S26**). BLUP_Time_ of this wavelength also did not recover this QTL (**Figure S27**). Modeling this phenotype as temperature-sensitive therefore improved the detection of this locus.

**Figure 13:**
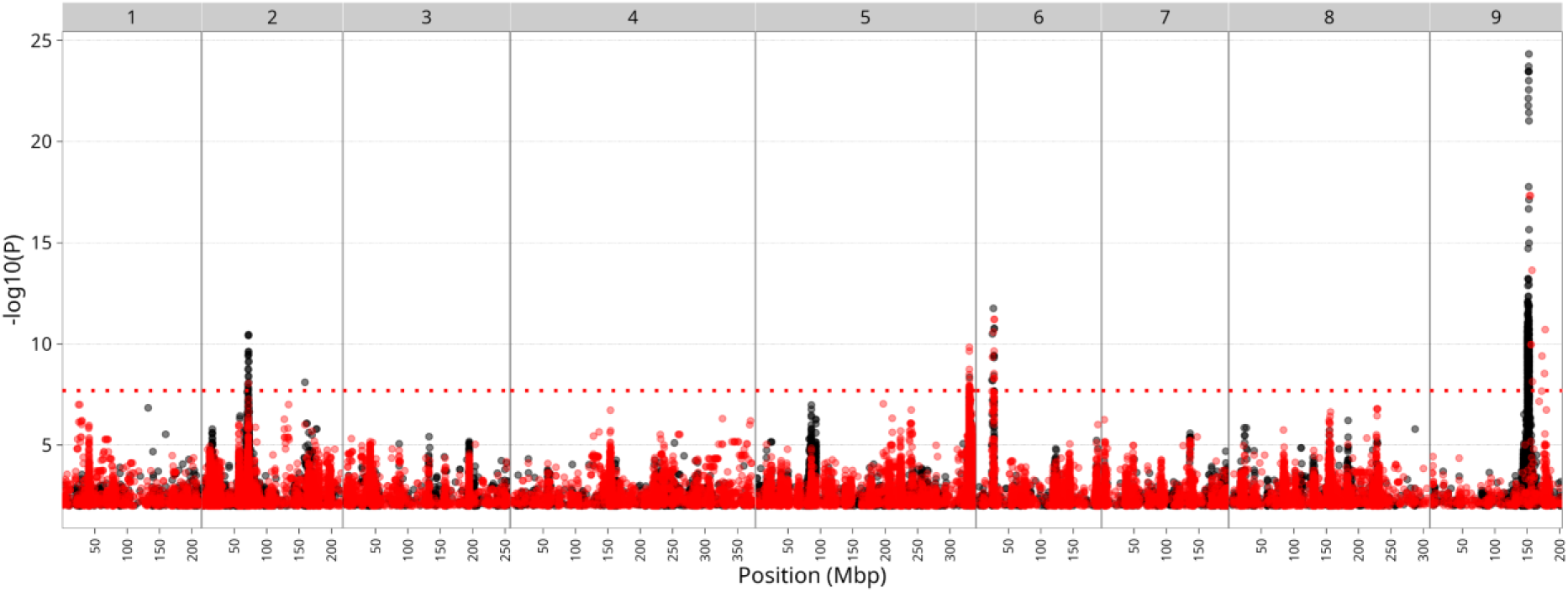
GWAS result for ARI BLUP_Temp_ in green *L. sativa* accessions. The Manhattan plot depicts SNPs associated with the ARI BLUP_Temp_ phenotype of green accessions. Significance is plotted as –log₁₀(p-value) on the y-axis. The x-axis shows genomic position in mega base pairs, with chromosome numbers indicated above. The red horizontal line shows the Bonferroni corrected significance threshold (–log₁₀(P) > 7.69). The coloring corresponds to iterative GWAS, where black is the result of the first iteration, and red is the result of the second iteration, where the top SNP of chromosome 9 was added as a fixed effect.

In addition, two novel QTLs on chromosome 5 (QTL_Chr5) and chromosome 6 (QTL_Chr6.1) were identified for BLUP_Temp_ of ARI in green varieties (**Table 1)**. QTL_Chr5 maps to 329.9-331Mbp, harboring 29 genes. A potential candidate is LOC111915094, a predicted E3 UBIQUITIN PROTEIN LIGASE DRIP1. In *Arabidopsis thaliana, DRIP1* is a regulator of Dehydration-responsive element-binding protein 2B (*DREB2B)*, a transcription factor controlling drought inducible gene expression [42]. As a second candidate, LOC111915113 encodes a histone H2A protein homologous to *AtH2A.Z* variants HTA8;9;11, which are well described for mediating responses to changes in temperature, as well as repressing anthocyanin accumulation through epigenetic regulation [43]. QTL_Chr6.1 maps to 25.2-27.9 Mbp, which includes 70 genes. A candidate is LOC111895179 (*LsERF088*), homologous to *AtSHN* transcription factors that are involved in wax biosynthesis, also conferring drought tolerance [44, 45].

### Field-related QTLs discovered by dimensionality reduction, BLUPs and single traits

We combined QTLs identified across single, PC, and BLUP-based phenotypes to provide an overview of all genomic regions associated with spectral trait variation of lettuce grown in this field trial. Over all timepoints, 155 distinct QTLs were identified (**Figure 14A**). Eighteen QTLs overlapped across all three GWAS sets, pointing to robust genetic loci consistently associated with phenotypic variation measured by different analytical approaches (**Figure 14B**). Among those, several QTLs were identified that could point to previously uncharacterized regions involved in stress responses. An overlapping QTL on chromosome 5 (277-279.1 Mbp) was associated with PC1 in green plants, the BLUP_Time_ of wavelength 894.5 nm, and 38 single traits. These included reflectance between 1570-1698 nm and two water-related vegetation indices (MSI and NDII), both involving ratios of wavelengths at 820 and 1600 nm. Together, this suggests a link to water content at this QTL. Out of 25 genes that map to this locus, LOC111888514 is an interesting candidate, as it is homologous to *AtERF043*, a Dehydration-responsive element-binding (DREB) transcription factor.

**Figure 14:**
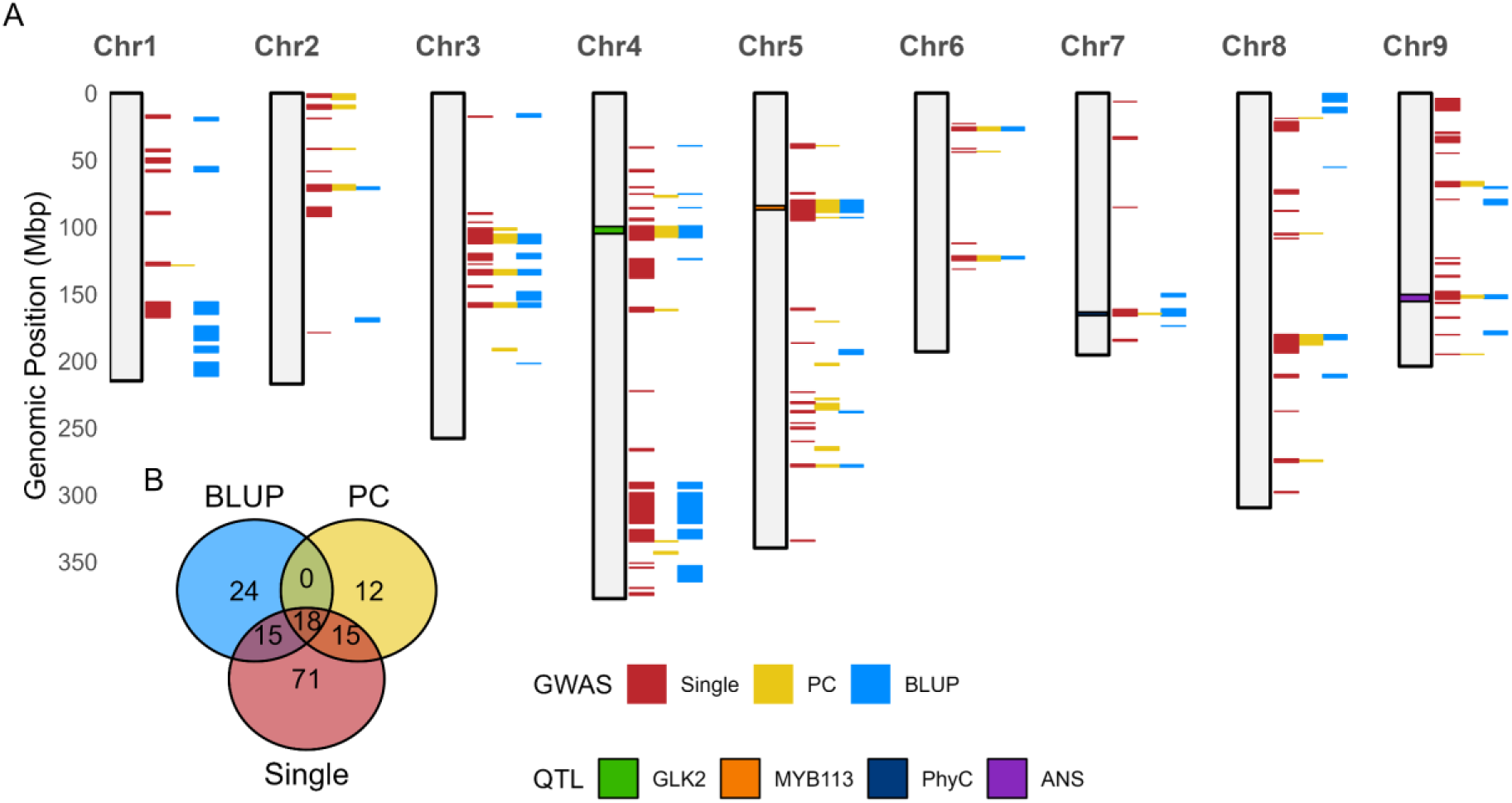
QTLs identified by GWAS using single phenotypes, principal components (PCs), and BLUPs. **A:** QTL positions are shown per chromosome, colored by GWAS type (red = single phenotypes, yellow = PC phenotypes, blue = BLUP phenotypes). The y-axis indicates genomic position in megabases (Mbp). Each QTL spans the region between the first and last significantly associated SNP (–log₁₀(P) > 7.69; Bonferroni-corrected). Only QTLs supported by at least three significant SNPs are shown. Well-characterized QTLs are highlighted within chromosomes: green = GLK2 (pale leaf locus), orange = MYB113, purple = ANS (anthocyanin accumulation), and dark blue = PhyC (flowering time). **B**: Venn diagram showing the overlap of QTLs across the three GWAS types, using the same color scheme as in panel A. All detected QTLs are included, including those with fewer than three significant SNPs.

Two overlapping QTLs on chromosome 6, QTL_Chr6.1 and QTL_Chr6.2, were identified ARI GWAS (**Table 1**) and observed exclusively in green accessions. Both QTLs were specific to timepoints 06-04 to 06-11, and associated with VIs Datt4 and Datt6, single wavelengths of 650 to 690nm, and BLUP_Temp_ traits of these wavelengths, all related to chlorophyll a+b content [46, 47]. QTL_Chr6.2 was also associated with the VI SIPI2, which measures the ratio between Chlorophyll a and carotenoids [48]. These findings suggest that QTL_Chr6.1 and QTL_Chr6.2 may play a role in pigment-linked responses to temperature-induced stress.

## Discussion

Spectral phenotyping of field-grown lettuce can reveal the genetic factors shaping field performance, but extracting meaningful signals from complex, environment- and time-dependent hyperspectral data remains challenging. Because spectral bands are highly correlated, dimensionality reduction helps uncover dominant patterns of phenotypic variation tied to plant physiology. In this study, PC1 and PC2 were strongly driven by water-absorption and chlorophyll bands, a pattern also reported for lettuce and other crop and ornamental species, where such bands have been proposed as markers for early stress monitoring [49–51]. Kumar et al. showed for lettuce grown in the greenhouse under drought stress that PC1 of hyperspectral imaging data primarily reflected water stress progression [51]. Although higher PCs explained only ∼20% of spectral variation, several QTLs mapped to them, indicating they can capture biologically meaningful signals beyond the dominant patterns. Higher PCs of hyperspectral data should therefore not be dismissed, even if most variation is captured by the lower components.

A caveat of PCA is that it assumes linear relationships among variables, so non-linear dynamics underlying spectral responses may be overlooked. Alternative approaches, including machine learning models, offer a valuable complement by extracting additional signals. In lettuce, deep learning models have successfully predicted key phenotypic traits such as soluble solids content and pH from hyperspectral reflectance, capturing spectral features beyond those identified with standard methods [11].

Spectral phenotypes captured both stable genetic signals and dynamic responses to field conditions. Variance decomposition revealed that genotype explained most spectral variance, with a higher contribution in the visible spectrum compared to NIR and SWIR, which has also been reported by Furlanetto et al [52]. This is consistent with the strong population structure in lettuce and clearly distinguishable morphology of horticultural types. Broad-sense heritability across the spectrum matched the variance-decomposition results, as both use comparable mixed-model frameworks to assess the genotypic fraction of variance. Phenotypic plasticity differed by horticultural type, suggesting spectral phenotyping might differentiate cultivars not only by static traits but by their temporal or stress-response dynamics. More importantly, hyperspectral monitoring may reveal early physiological responses to environmental stress before visible symptoms appear, offering a predictive window for selecting resilient varieties [53].

While genotype-by-environment interactions are often characterized across multiple years or sites to encompass broad environmental variation, these approaches usually rely on single timepoint phenotyping [10]. By incorporating temporal resolution, our study captured dynamic genotype responses to fluctuating field conditions, providing a complementary view of G×E that static trials cannot resolve, though further replication across seasons would enhance robustness.

GWAS across spectral phenotypes provided complementary insights into lettuce field-trait genetics. Single-wavelength GWAS linked QTLs to physiologically relevant absorption features, allowing interpretation of what a wavelength reflects when the QTL function is known, such as wavelengths that underly the known pigmentation QTLs in lettuce. The reverse is less straightforward: knowing the physiology behind a spectral feature suggests, but does not confirm, the function of the underlying QTL. PC-GWAS captured both major and minor axes of variation, with PC1-3 confirming associations with water content, chlorophyll, and anthocyanin [16, 18, 51]. Tracking PCs over time revealed shifts in spectral axes and QTLs, with late-stage PC1 and PC2 mapping to the bolting-associated locus *PhyC*, illustrating how combining dimensionality reduction with longitudinal data can uncover links between complex spectral variation and trait genetics, even when the precise interpretation of a PC changes.

BLUP-GWAS revealed temperature- and development-responsive QTLs. For example, QTL_ANS on chromosome 9 was not detected using mean reflectance at the anthocyanin absorption peak (556 nm) but became apparent when reflectance was modeled as a temperature-responsive, plastic phenotype. This demonstrates that integrating environmental dynamics can uncover subtle signals in spectral phenotypes. BLUP_Time_ GWAS revealed a remarkably broad QTL at the end of chromosome 1, spanning nearly 40Mbp, possibly linked to change in water content or cell structure. The size of this QTL indicates a broad LD structure, which can be caused by the presence of a large structural variation [54]. In lettuce, this region coincides with a large inversion present only in Crisp varieties [55]. Such inversions can be used in crop breeding to control recombination rates and stabilize alleles of favorable traits [56]. The association may reflect a horticultural type-specific haplotype rather than a direct effect on water content, although the inversion could still contribute to Crisp-specific structural traits.

Together, these results demonstrate that integrating longitudinal hyperspectral phenotyping with genetic and environmental analyses enables high-throughput characterization of both stable and dynamic components of trait genetics. In lettuce and other field crops, such approaches can accelerate the discovery of adaptive loci and inform breeding for resilience, developmental timing, and product quality. As hyperspectral phenotyping technologies continue to advance, integrating them with genomic tools will further improve our ability to connect genotype, phenotype, and environment in field conditions.

## Acknowledgements

We thank the Netherlands Plant Eco-phenotyping Centre (NPEC) for their help with operating the hyperspectral camera platform (TraitSeeker) and first steps in data exchange.

## Author contributions

GvdA obtained funding for the project. BLS and SM conceptualized the study. SM, AZ, MdH, and BLS extracted the data and performed the analysis. SM wrote the manuscript with input from all co-authors.

## Funding

This publication is part of the LettuceKnow project (with project number 1.2 of the research Perspective Program P19-17 which is (partly) financed by the Dutch Research Council (NWO; TTW) and the breeding companies BASF, Bejo Zaden B.V., Limagrain, Enza Zaden Research & Development B.V., Rijk Zwaan Breeding B.V., Syngenta Seeds B.V., Takii and Company Ltd., and the Foundation for Food and Agriculture Research (FFAR).

## Supplement

**Figure S1:**
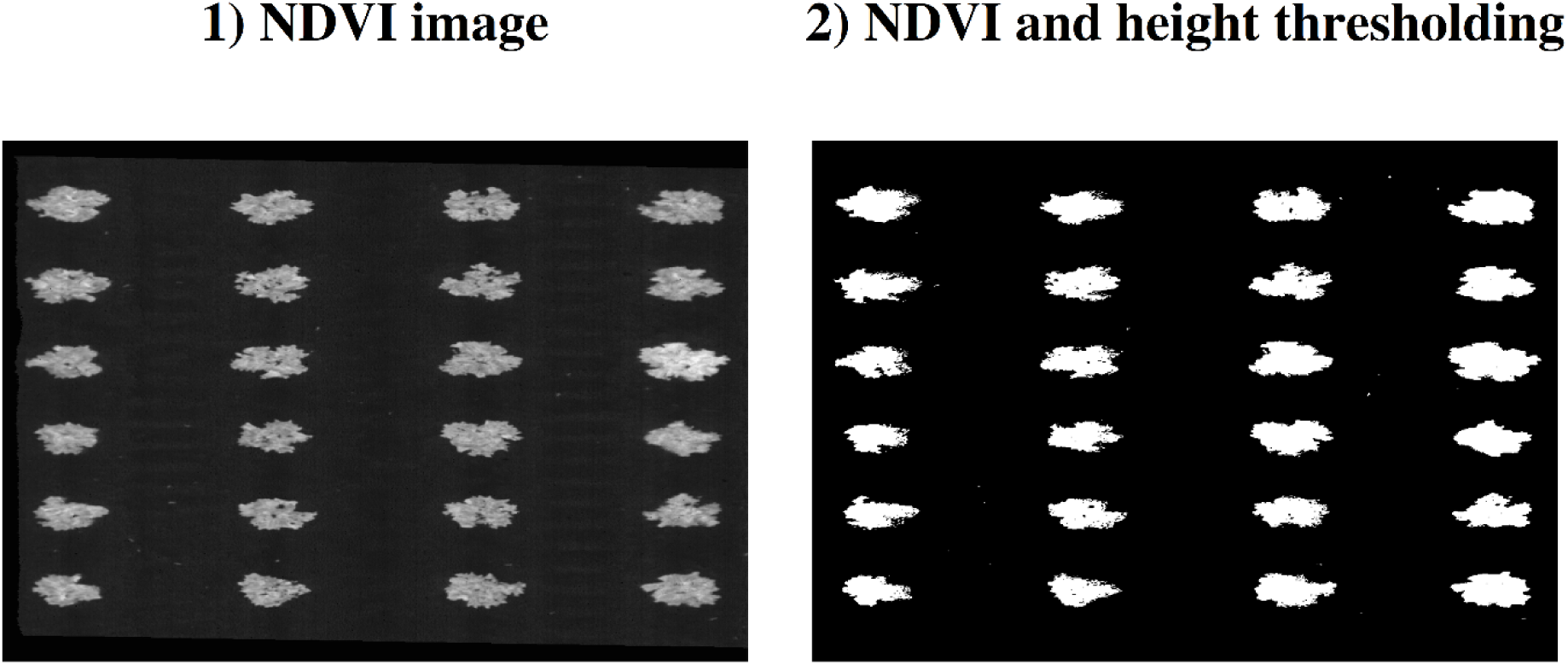
Image segmentation to extract plant pixels from images taken by FX10. Image segmentation process of FX10 generated images for plot IL550 at the first timepoint (20210521). To exemplify this process, images of the intermediary steps are shown. Step 1) is a generated NDVI, Step 2) is the fully segmented image after NDVI and heigh thresholding.

**Figure S2:**
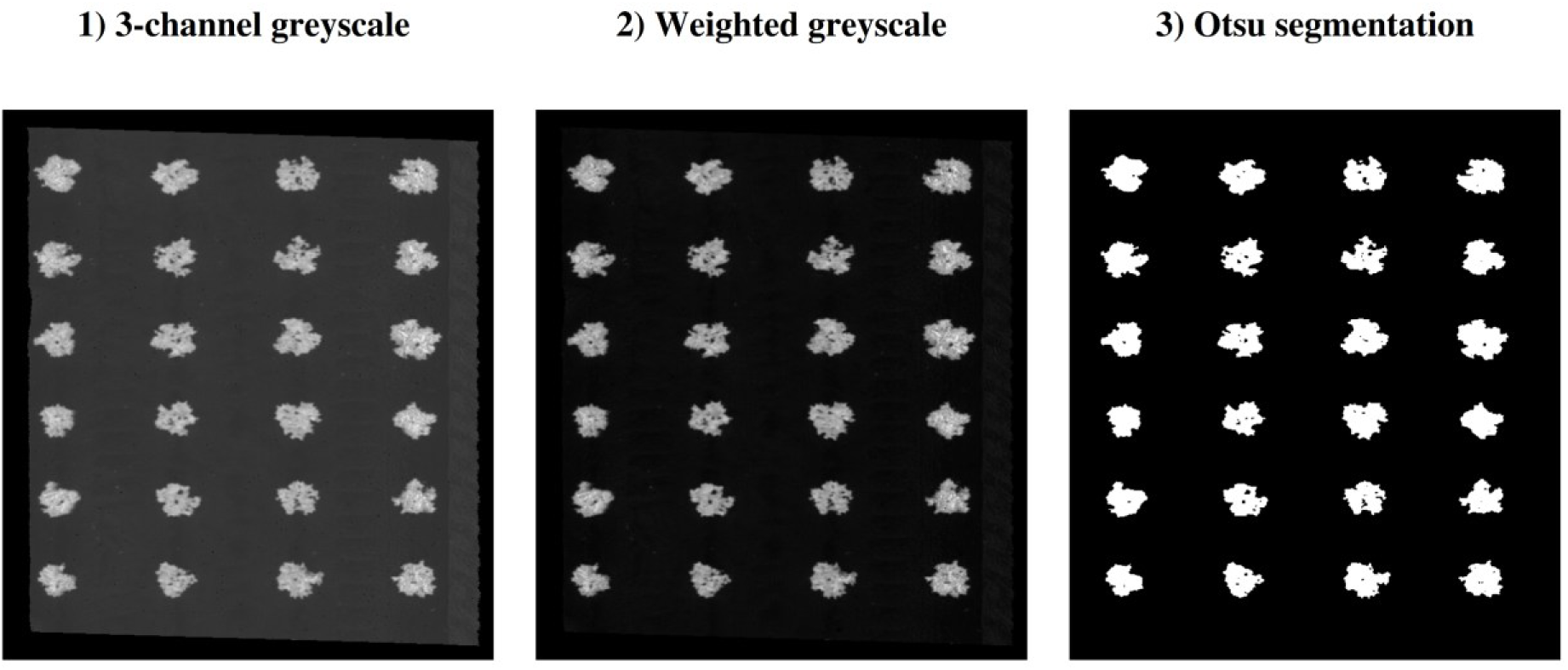
Image segmentation to extract plant pixels from images taken by FX17. Image segmentation process of FX17 generated images for plot IL550 at the first timepoint (20210521). To exemplify this process, images of the intermediary steps are shown. Step 1) is a 3-channel greyscale image, where each channel represents the log2-transformed ratio of pixel values measured at 967nm and 1457nm. Step 2) is the weighted conversion of the 3-channel greyscale to single-channel greyscale. Step 3) is the fully segmented image after employing Otsu thresholding.

**Figure S3:**
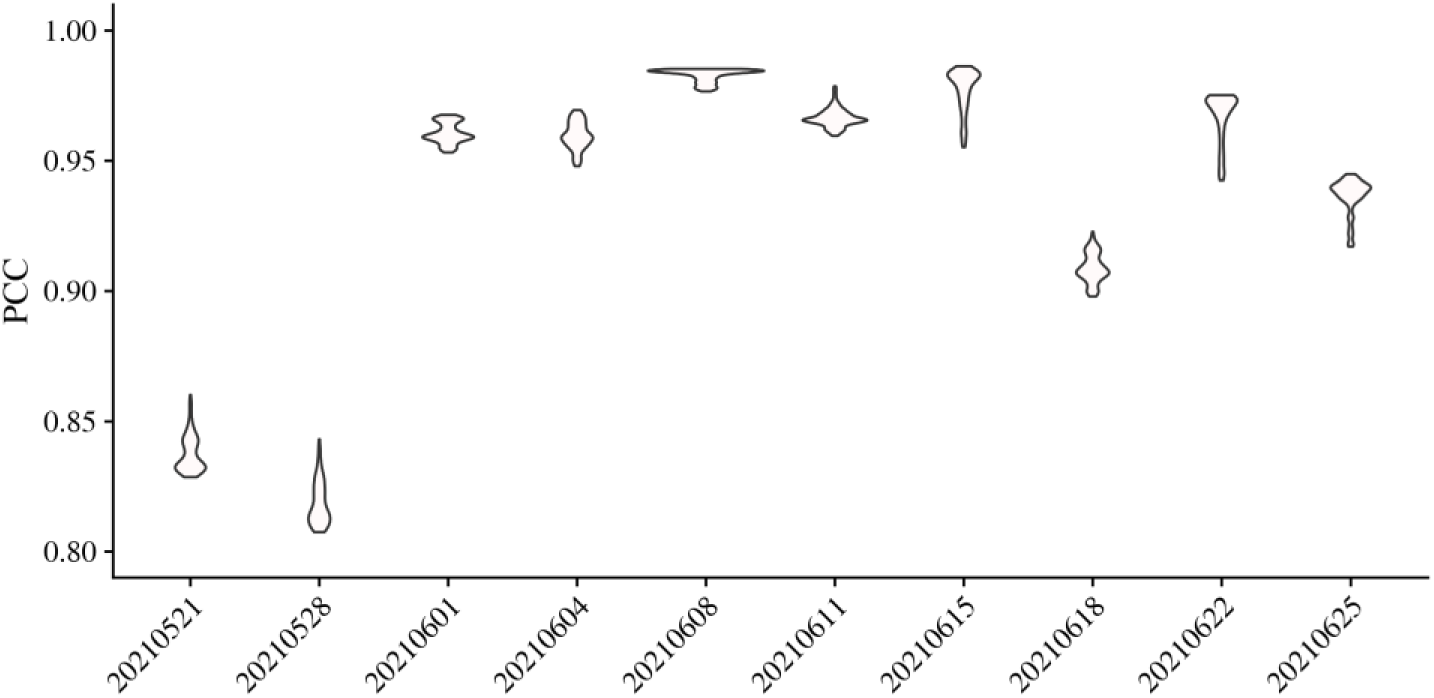
Correlation of overlapping light spectra between FX10 and FX17 images over time. Violin plots display the Pearson correlation coefficients (PCCs) between images of spectral bands (935–1004 nm) captured by both FX10 and FX17 cameras across time points. Pairwise Pearson correlations were computed using the plot-level reflectance averaged over the two replicate plots for each spectral band.

**Figure S4:**
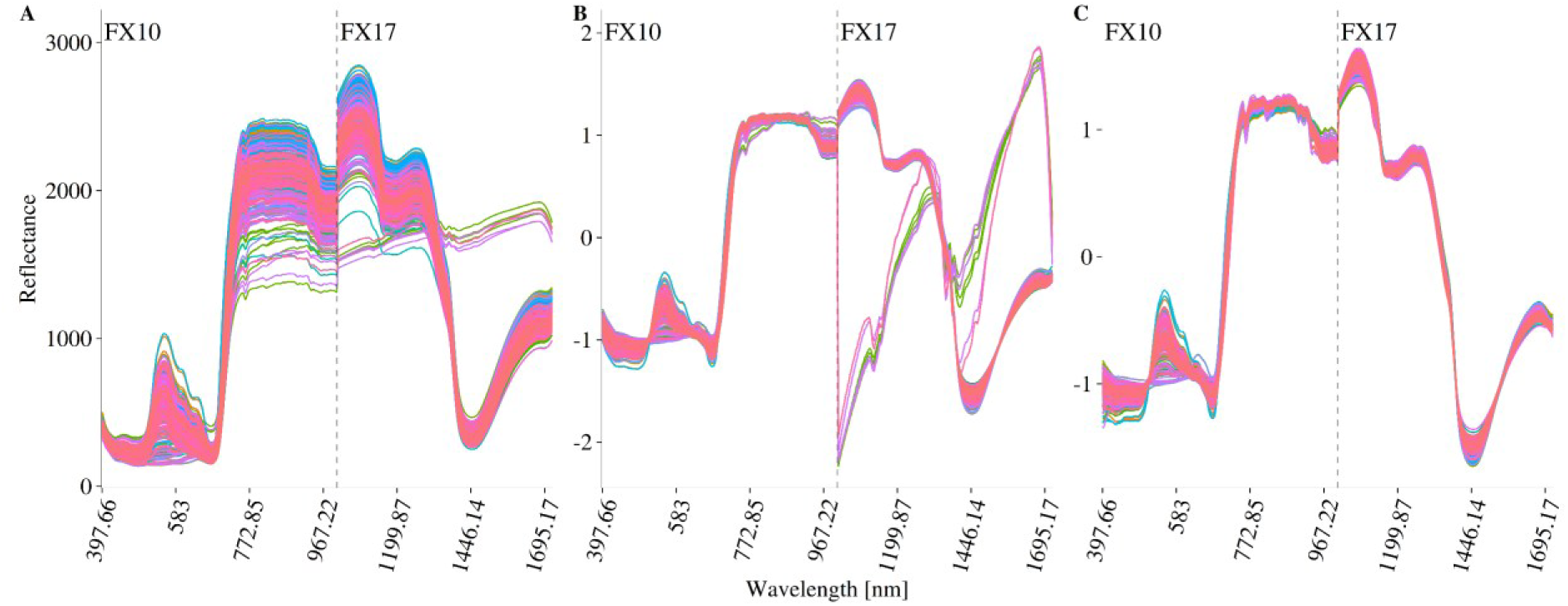
Process of SNV transformation and outlier removal for timepoint 3 (2021-06-01). Plot-averaged spectral curves of all replicates measured at timepoint 3. Line color is used to distinguish plots (accessions and replicates). Vertical line separates FX10 from FX17 measured wavelengths, as labelled in each plot. The x-axis is discretized and does not show all wavelengths that were measured. **A**: Spectral curves of replicates before data cleaning. The y-axis represents the measured reflectance values. **B**: Spectral curves of replicates after SNV transformation. Y-axis indicates transformed reflectance values. **C**: Spectral curves of replicates after SNV transformation with FX17 outlier values removed.

**Figure S5:**
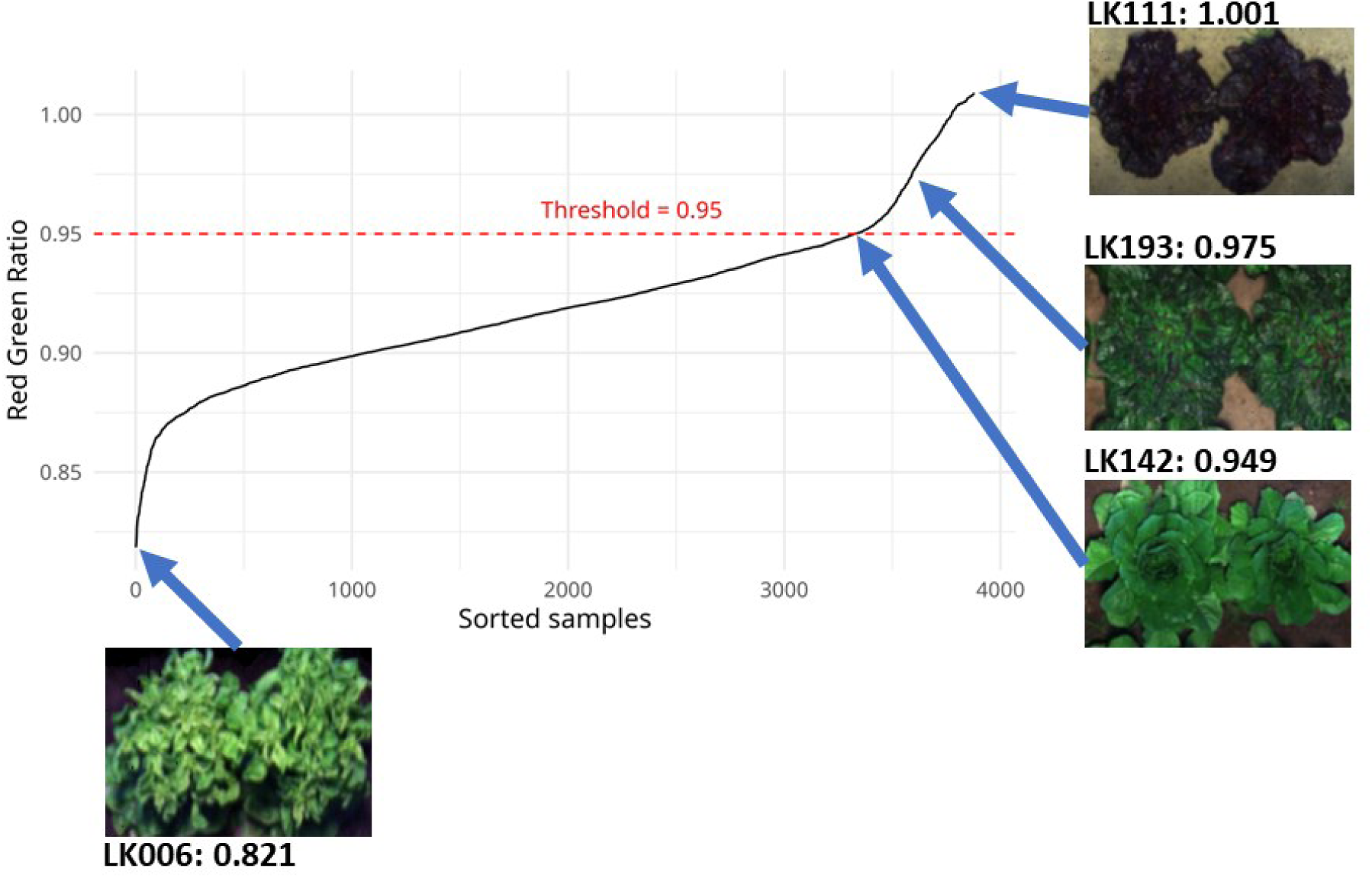
Ranked value plot of Red-Green Ratios of all sampling points. X-axis represents sorted Red-Green Ratio values (Y-axis) across all replicates and timepoints. The horizontal red line marks the threshold of classifying samples as green (< 0.95) or red varieties (≥ 0.95). Exemplary RGB images of varieties LK006, LK111, LK142 and LK193, taken during hyperspectral imaging, illustrate the relationship between the Red-Green ratio and the observed pigmentation. The ratio value is indicated by arrow and image caption.

**Figure S6:**
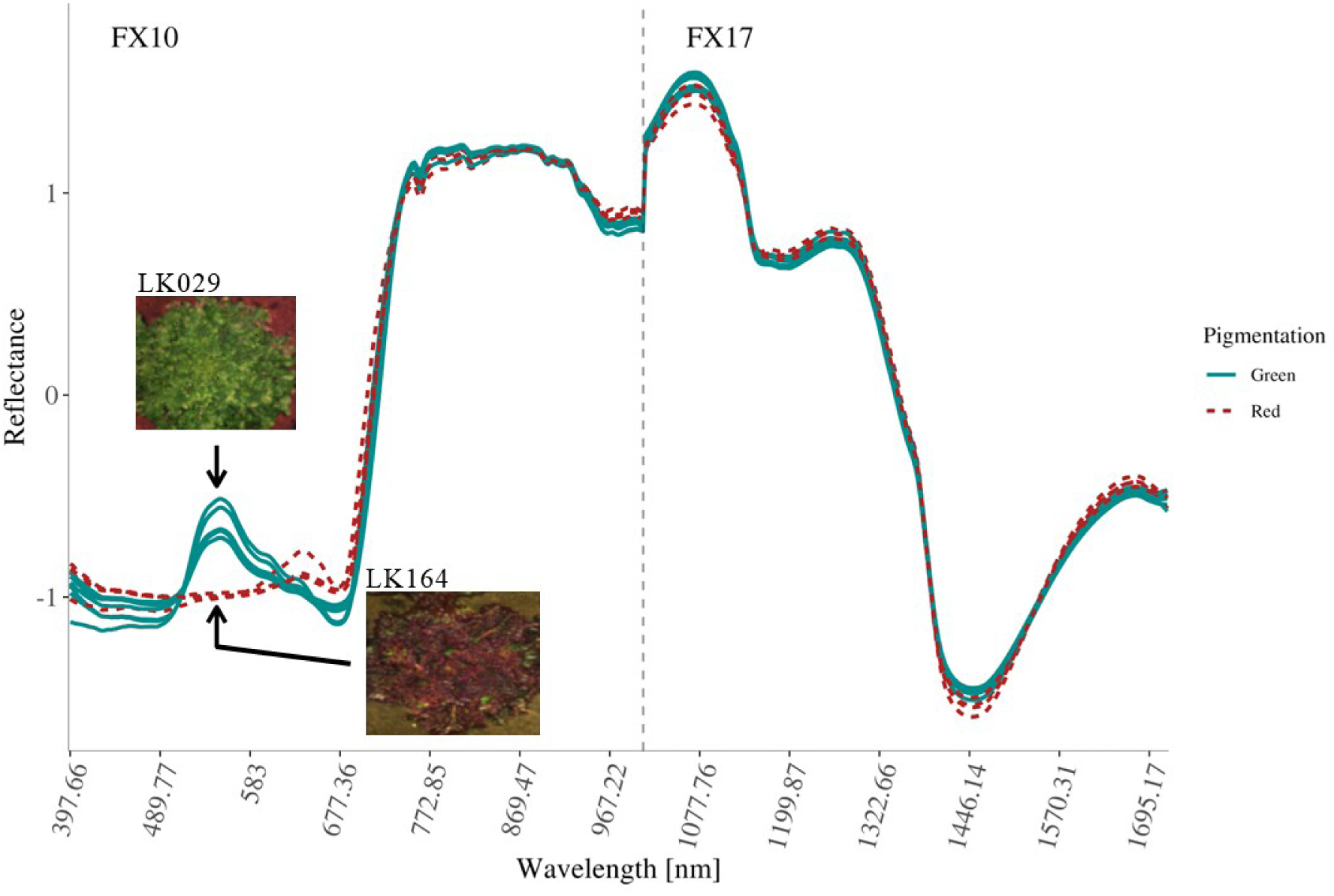
Plot-averaged spectral reflectance curves of red and green *L. sativa* varieties at timepoint 3 (2021-06-01). Line color distinguishes red and green varieties. The x-axis shows discretized measured wavelengths, separated by a vertical line marking the transition from FX10 to FX17 sensor ranges. The y-axis indicates transformed reflectance values. Arrows highlight the spectral shape difference at 584.24 nm. An exemplary RGB image of varieties LK029 (green) and LK164 (red), taken during hyperspectral imaging on the same date, illustrates contrasting plant pigmentation.

**Figure S7:**
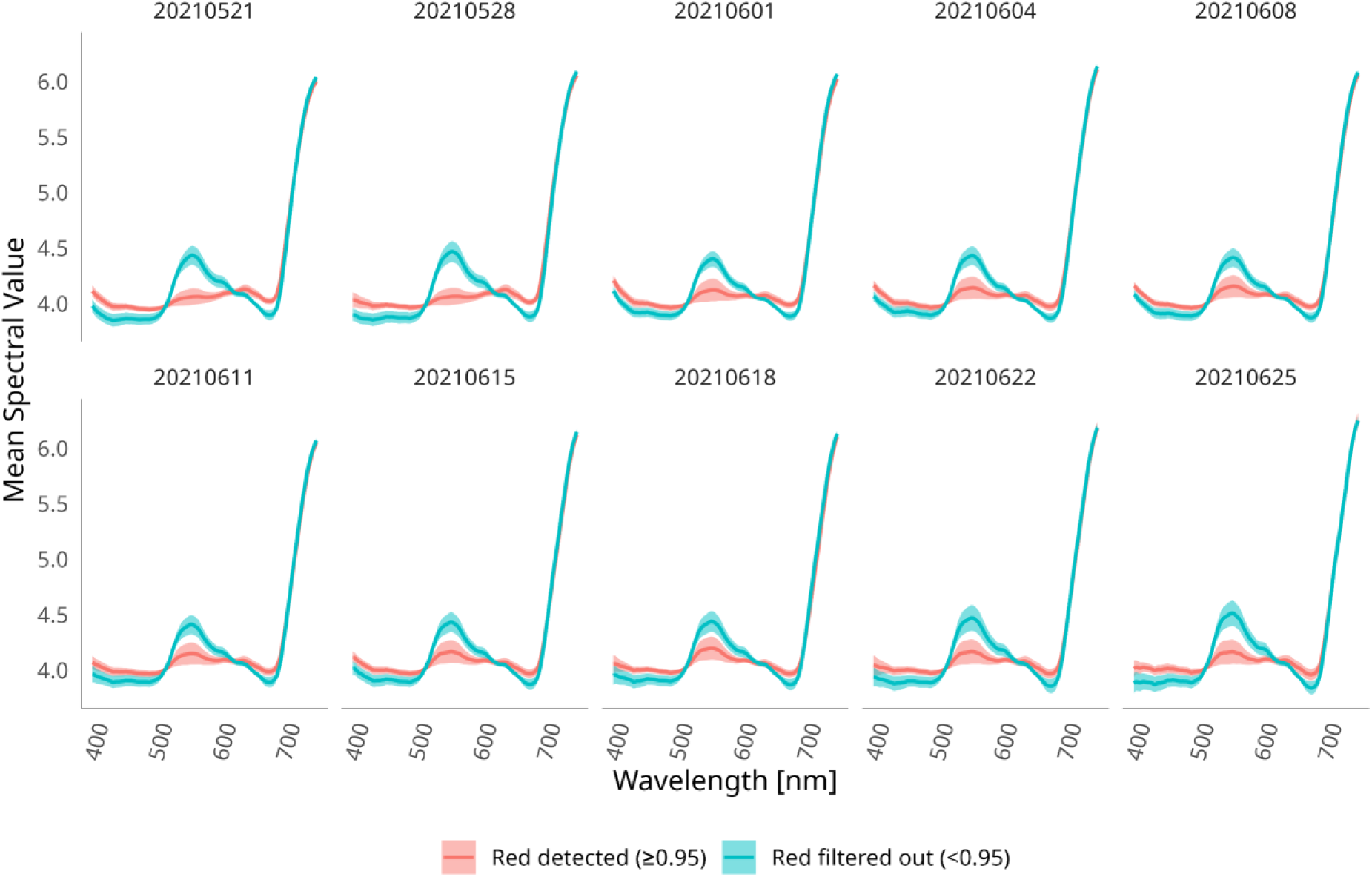
Mean spectral reflectance curves (400–750 nm) across ten timepoints for green and red *L. sativa* accessions. Accessions are classified as green (Red-Green Ratio < 0.95, cyan line color) and red (Red-Green Ratio ≥ 0.95, pink line color). Shaded areas indicate standard deviation.

**Figure S8:**
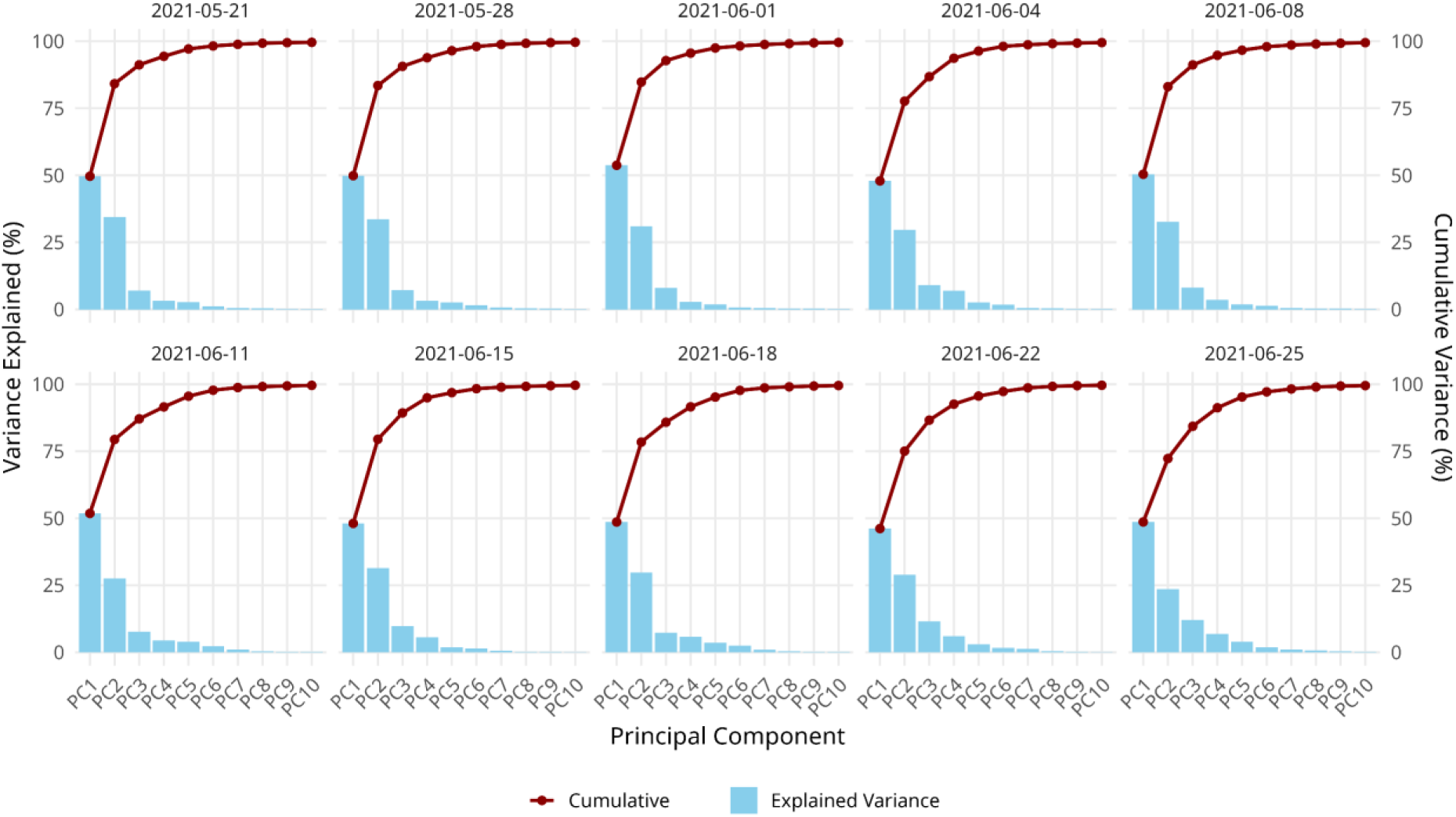
Scree plots describing explained variance of the first 10 principal components of wavelength phenotypes in all *L. sativa* accessions per timepoint. Each subplot represents a timepoint and displays individual percent explained variance (blue bars) and percent cumulative explained variance (red line) per PC.

**Figure S9:**
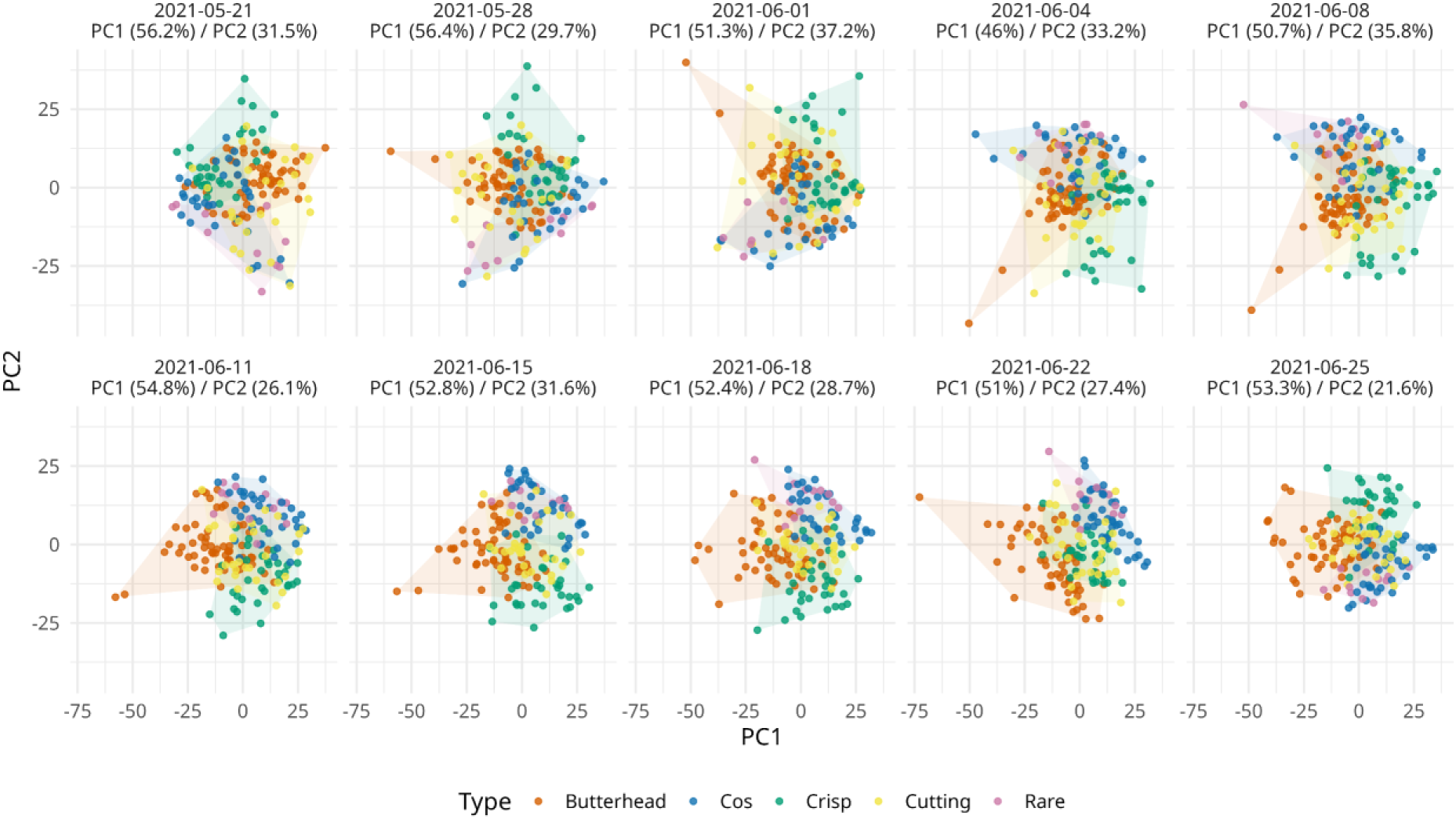
Principal component analysis (PCA) of spectral phenotypes of green *L. sativa accessions* across ten timepoints. Each panel represents a separate timepoint, with PC1 and PC2 capturing most of the variation (percentage of variance explained indicated per plot). Points represent individual genotypes, colored by horticultural type: Butterhead (orange), Cos (blue), Crisp (green), Cutting (yellow), and Rare (purple). Hulls illustrate the spread of each type.

**Figure S10:**
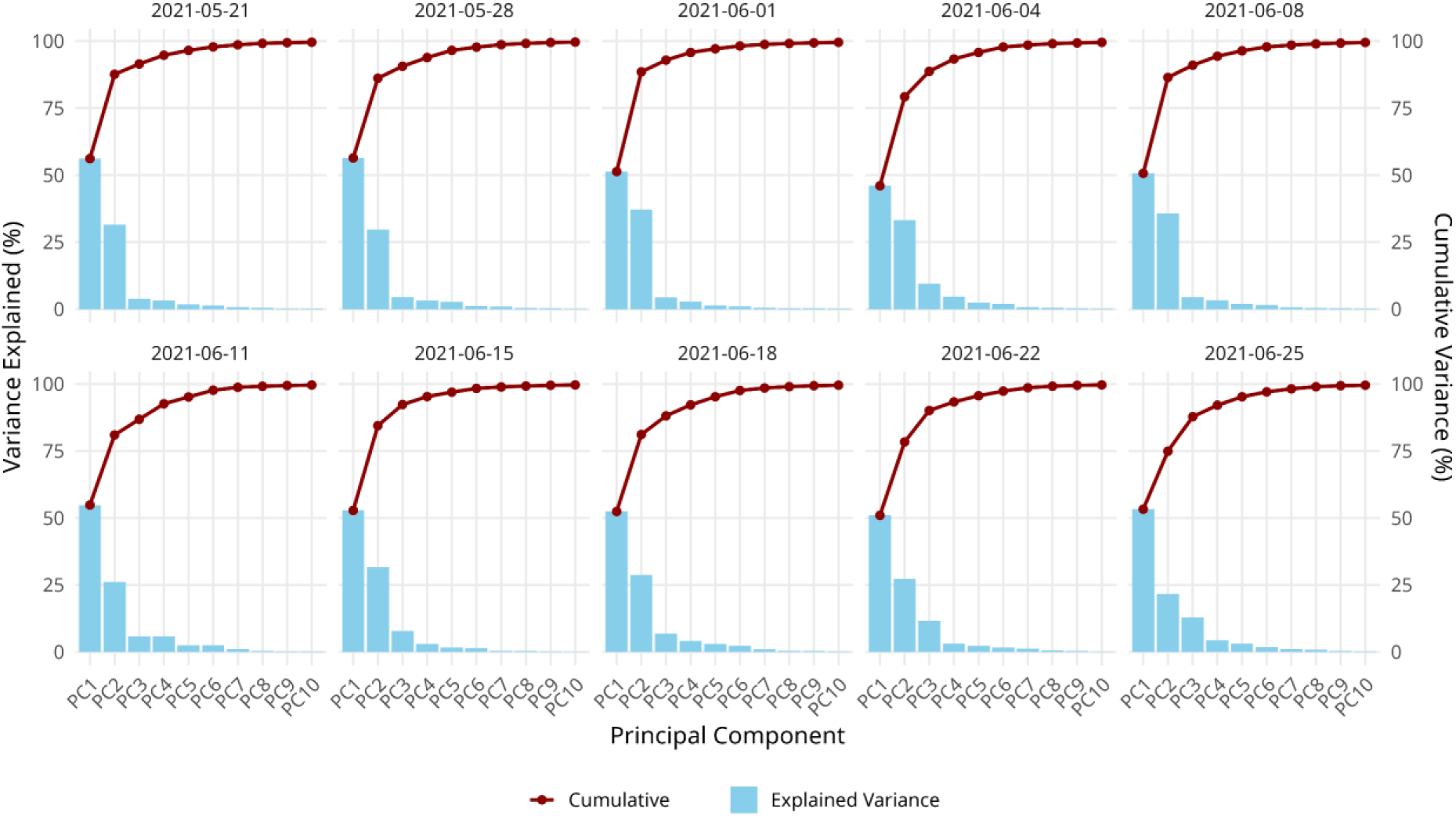
Scree plots describing explained variance of the first 10 principal components of wavelength phenotypes in green *L. sativa* accessions per timepoint. Each subplot represents a timepoint and displays individual percent explained variance (blue bars) and percent cumulative explained variance (red line) per PC.

**Figure S11:**
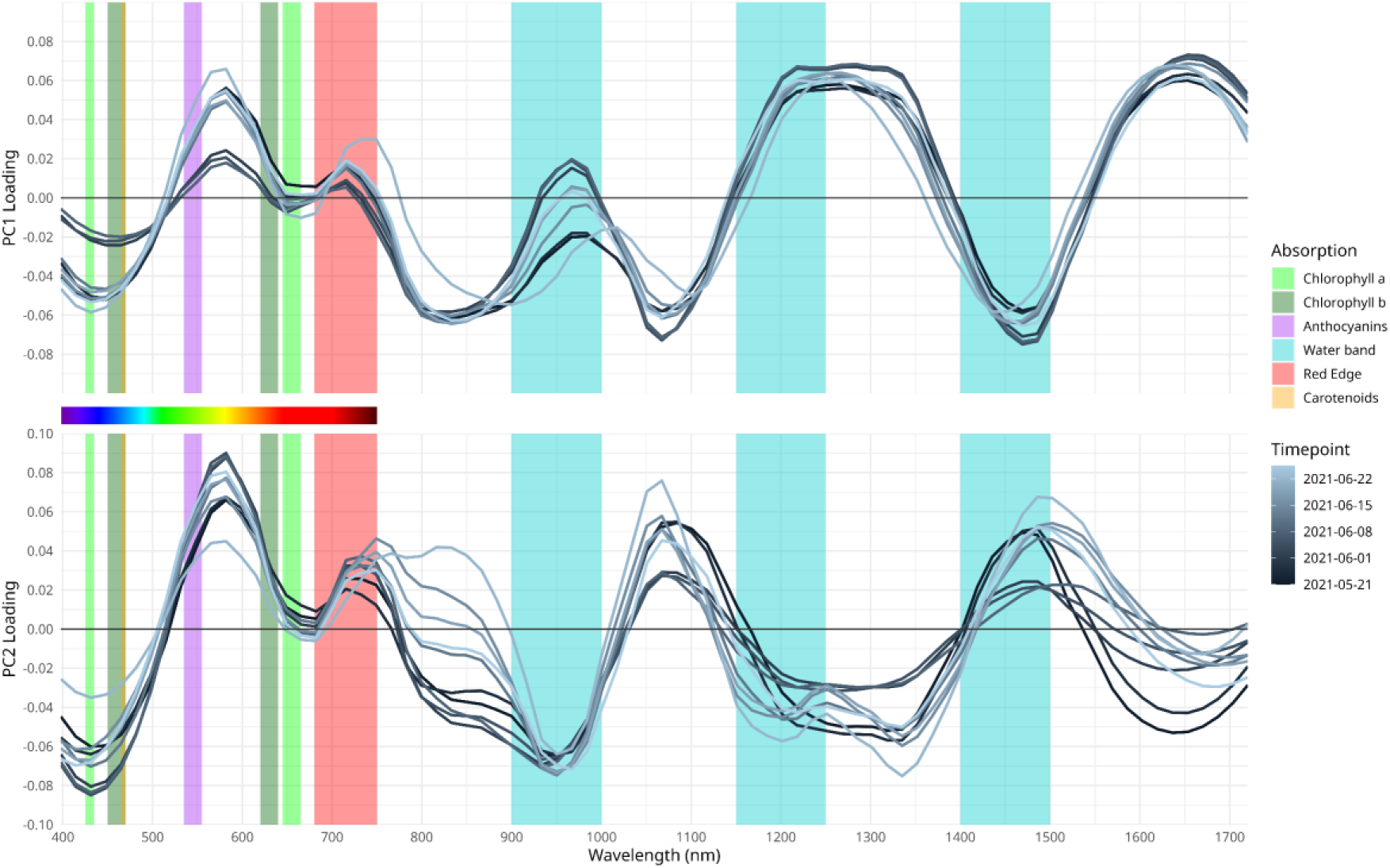
Loadings of wavelengths along the first two PCs over time in green *L. sativa accessions*. Loadings of PC1 (Top) and PC2 (Bottom) are plotted against the respective wavelength (x-axis) using a geom_smooth with the “loes” method with a span=0.2. Line color indicates time progression, with darker lines representing earlier timepoints and lighter lines later ones. Colored background bars indicate absorption spectra of known plant pigments and spectral features of plants. The visible light spectrum is shown as a color gradient between the panels to indicate approximate wavelength positions.

**Figure S12:**
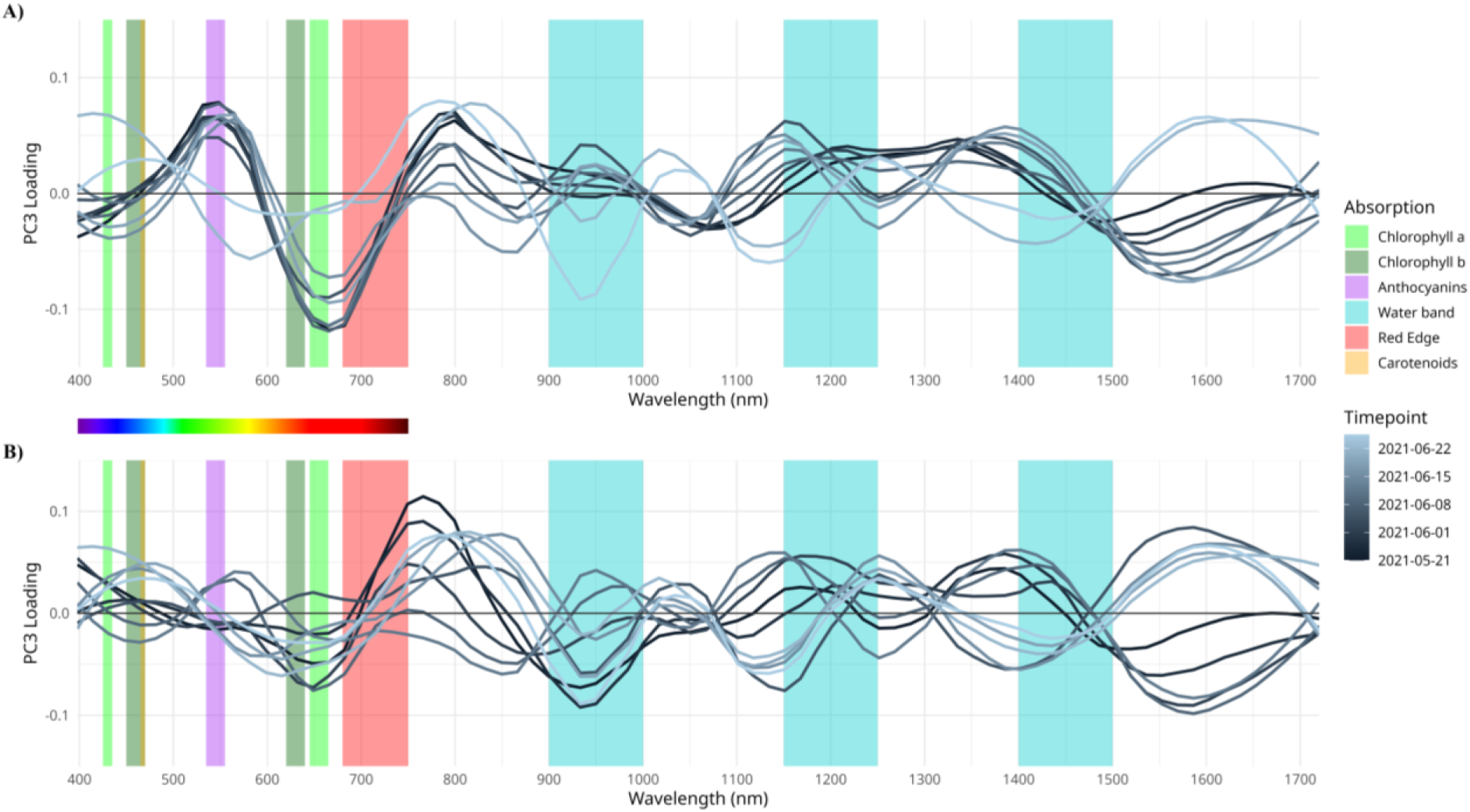
Loadings of wavelengths along the third principal component (PC3) across all *L. sativa* accessions (A) and green-only accessions (B) over ten timepoints. Loadings of PC3 are plotted against the respective wavelength (x-axis) using a geom_smooth with the “loes” method with a span=0.2. Line color indicates time progression, with darker lines representing earlier timepoints and lighter lines later ones. Colored background bars indicate absorption spectra of known plant pigments and spectral features of plants. The visible light spectrum is shown as a color gradient between the panels to indicate approximate wavelength positions.

**Figure S13:**
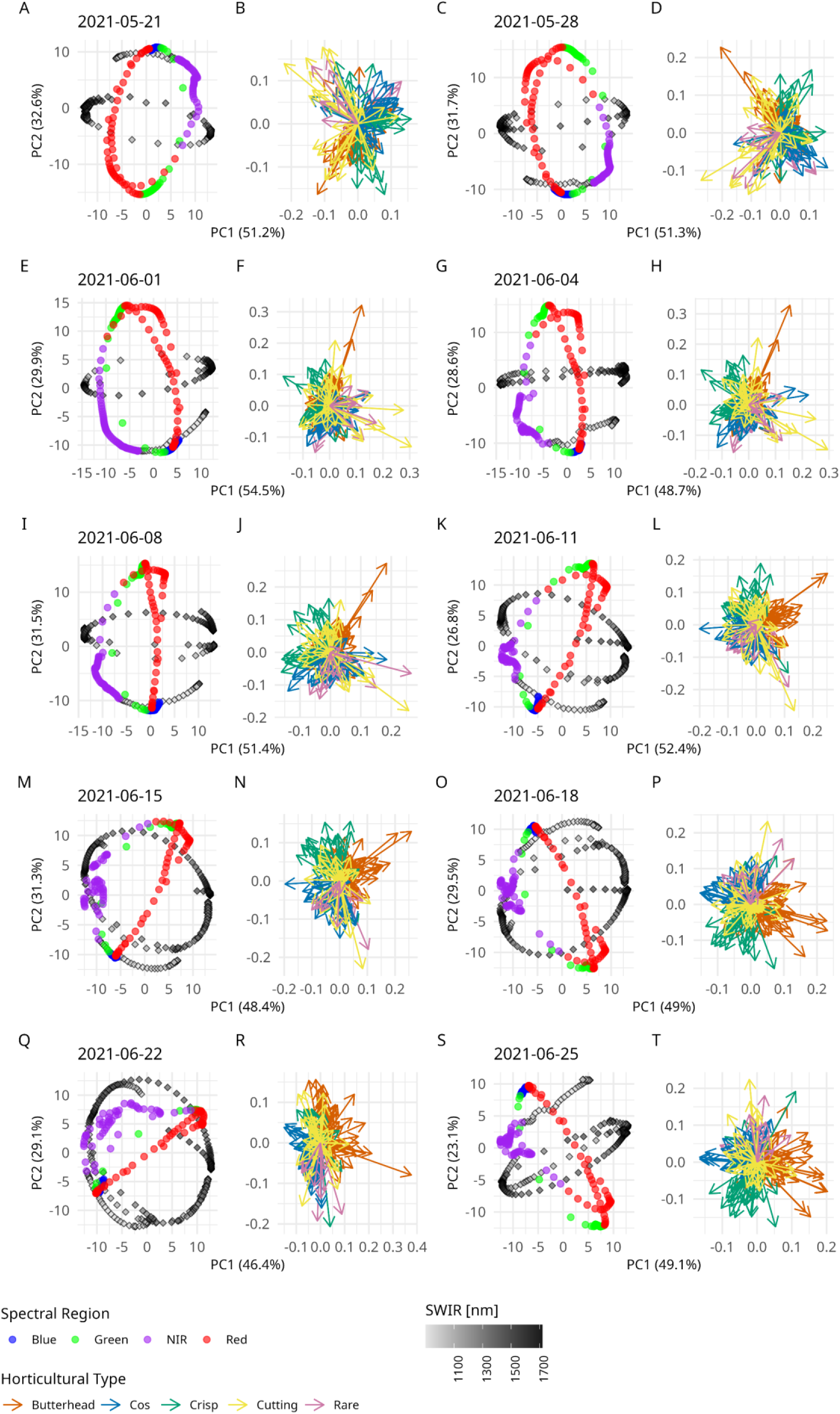
Reciprocal PCA plots of spectral phenotypes for all *L. sativa* accessions across ten timepoints. Each pair of plots corresponds to a timepoint and includes the PCA results based on the first two principal components (left), and the corresponding vector field (right). Axes represent the first (x-axis) and second (y-axis) principal components, with the percentage of variance explained indicated in parentheses. Data points of the PCA are color-coded by spectral region (Blue, Green, Red, purple:VNIR, grey to black: SWIR). Vectors are colored by horticultural type: Butterhead (orange), Cos (blue), Crisp (green), Cutting (yellow), and Rare (purple).

**Figure S14:**
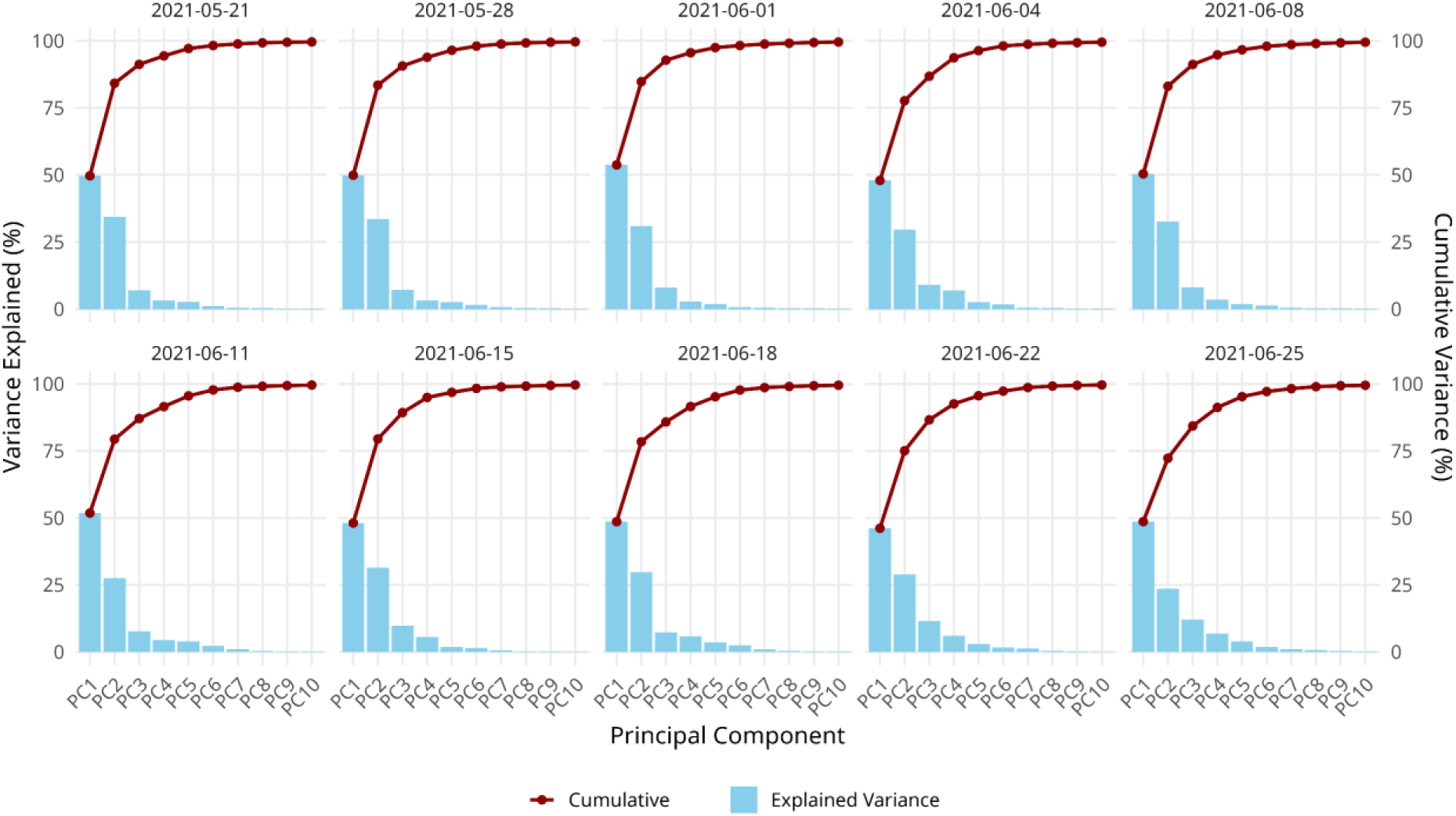
Scree plots showing the explained variance of the first ten principal components from the reciprocal PCA of wavelength phenotypes in all *L. sativa* accessions across timepoints. Each subplot represents one timepoint and displays the individual percent variance explained by each component (blue bars) and the cumulative explained variance (red line).

**Figure S15:**
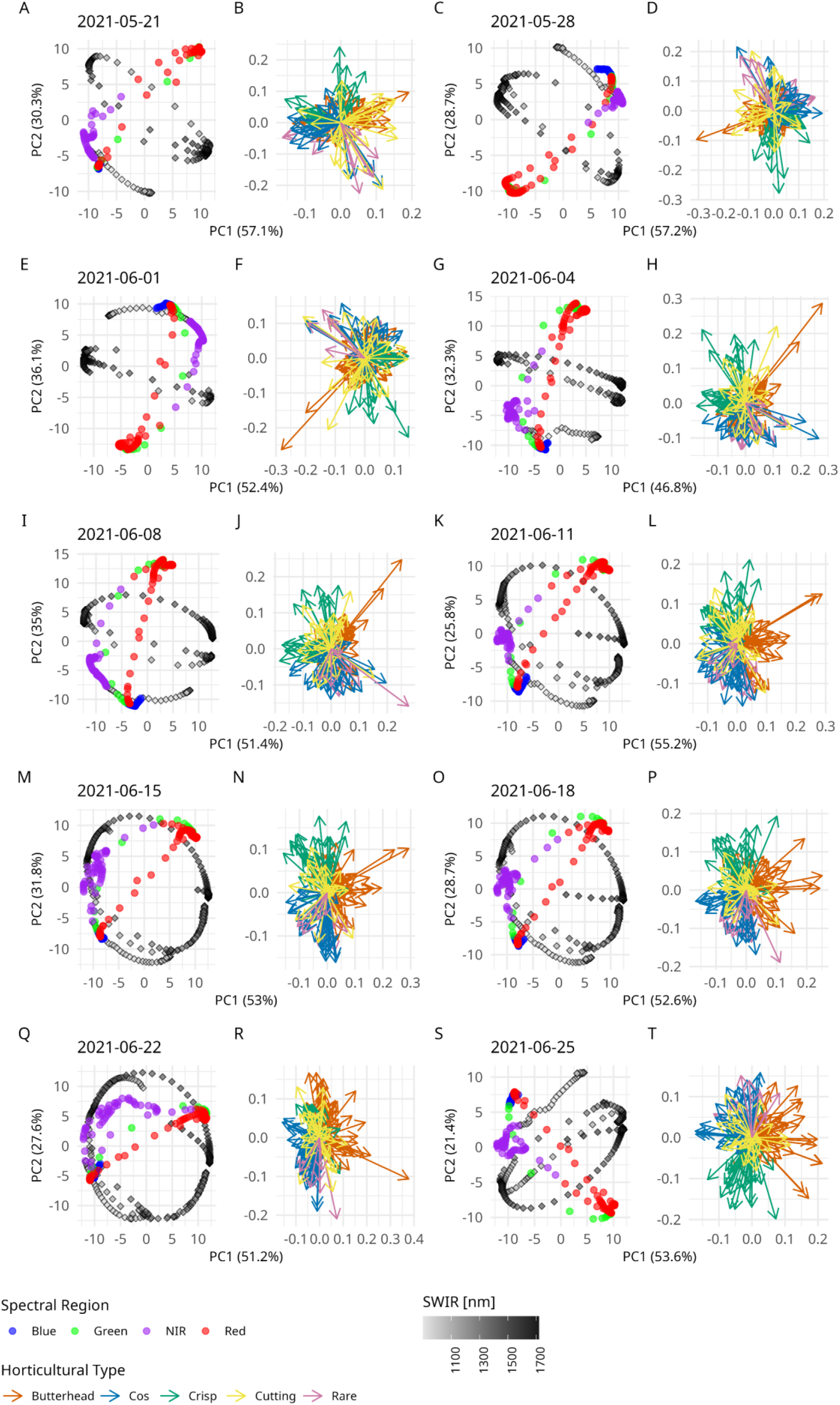
Reciprocal PCA plots of spectral phenotypes for green-only *L. sativa* accessions across ten timepoints. Each pair of plots corresponds to a timepoint and includes the PCA results based on the first two principal components (left), and the corresponding vector field (right). Axes represent the first (x-axis) and second (y-axis) principal components, with the percentage of variance explained indicated in parentheses. Data points of the PCA are color-coded by spectral region (Blue, Green, Red, purple:VNIR, grey to black: SWIR). Vectors are colored by horticultural type: Butterhead (orange), Cos (blue), Crisp (green), Cutting (yellow), and Rare (purple).

**Figure S16:**
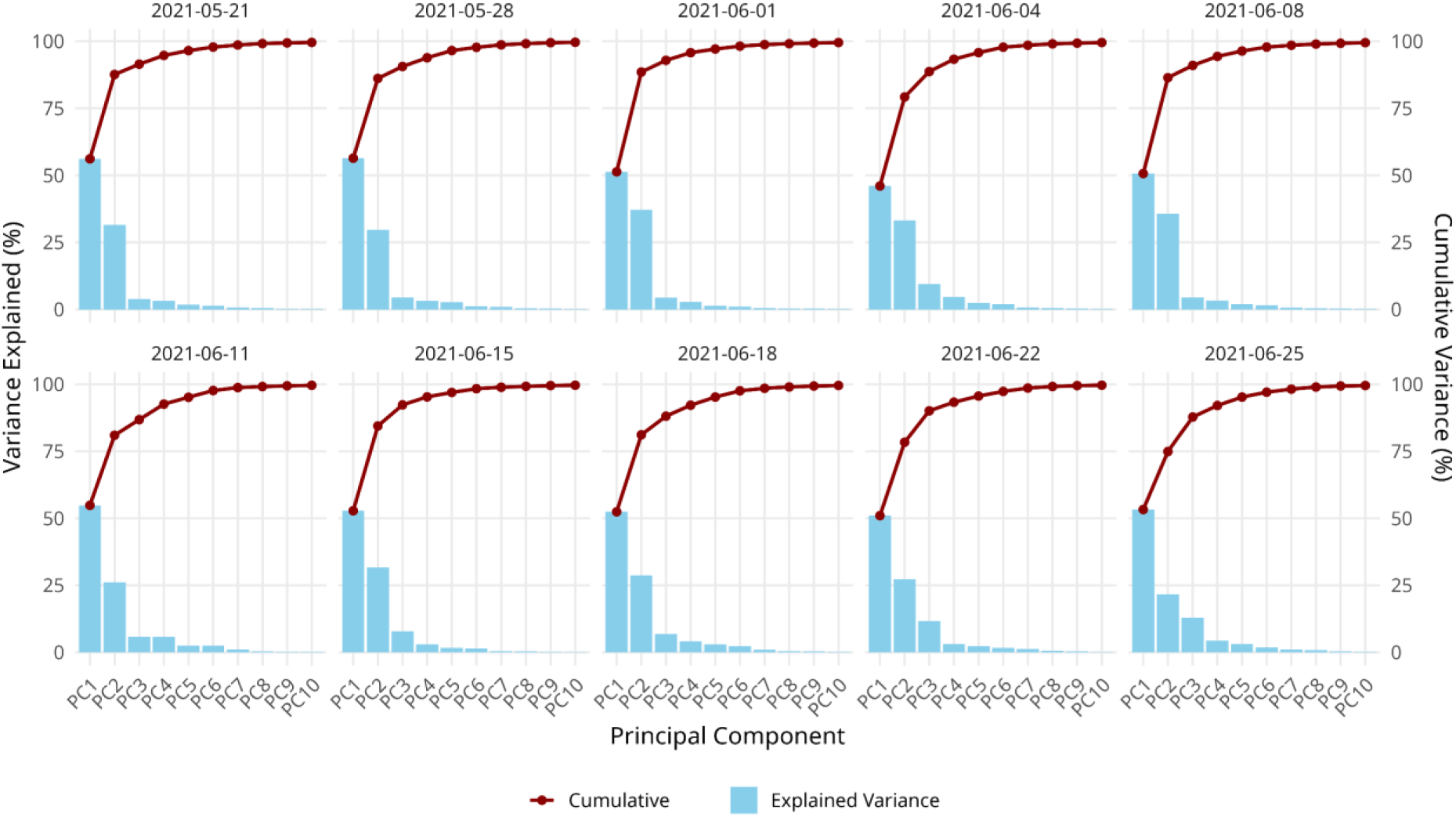
Scree plots showing the explained variance of the first ten principal components from the reciprocal PCA of wavelength phenotypes in green-only *L. sativa* accessions across timepoints. Each subplot represents one timepoint and displays the individual percent variance explained by each component (blue bars) and the cumulative explained variance (red line).

**Figure S17:**
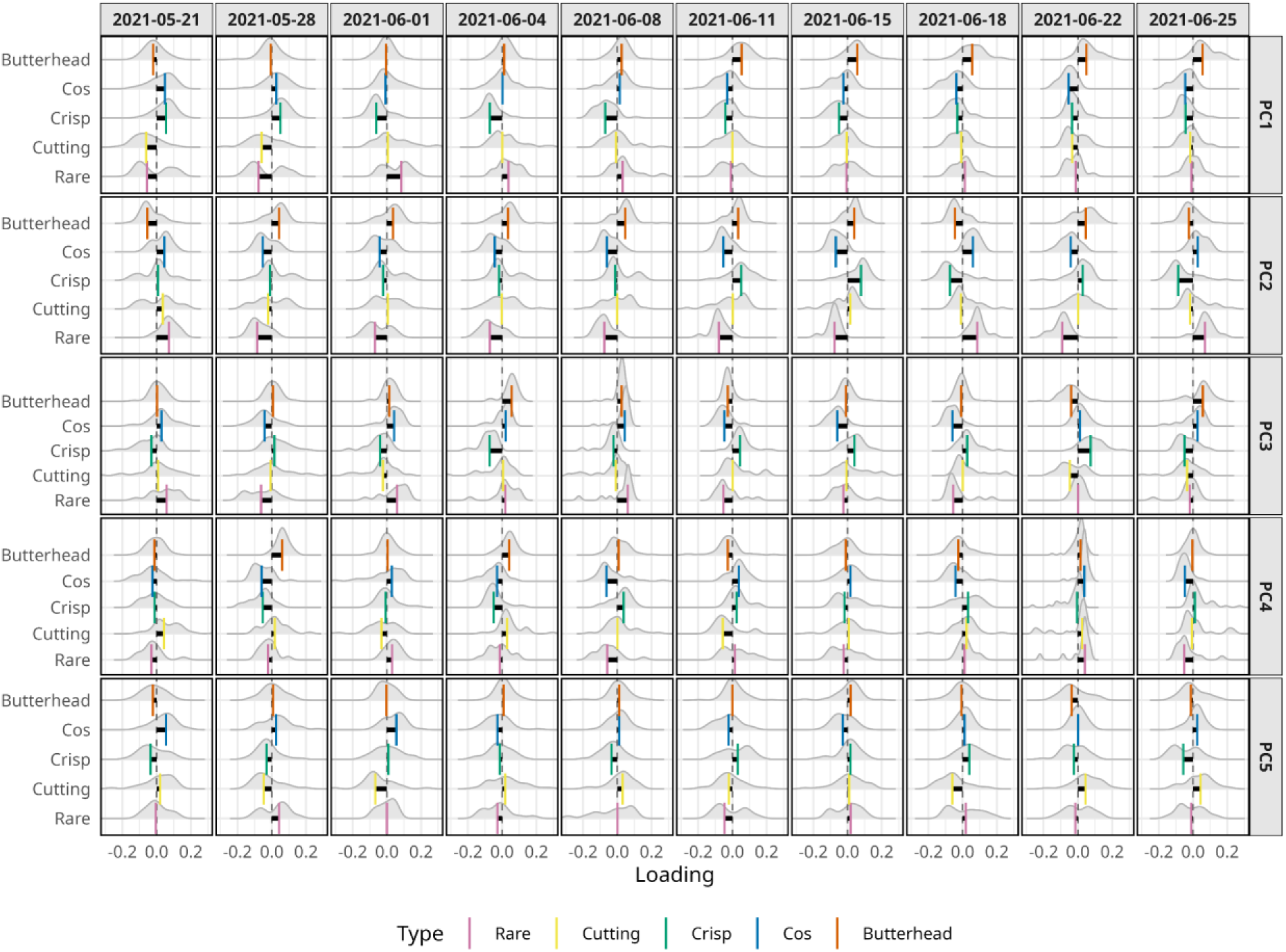
Distribution of reciprocal PCA loadings (PC1 to PC5) across all *L. sativa* accessions grouped by horticultural type. Each row represents one principal component (PC1–PC5), and each column corresponds to a timepoint. Density plots show the distribution of loadings per horticultural type, with mean values indicated by colored bars: Butterhead (orange), Cos (blue), Crisp (green), Cutting (yellow), and Rare (purple).

**Figure S18:**
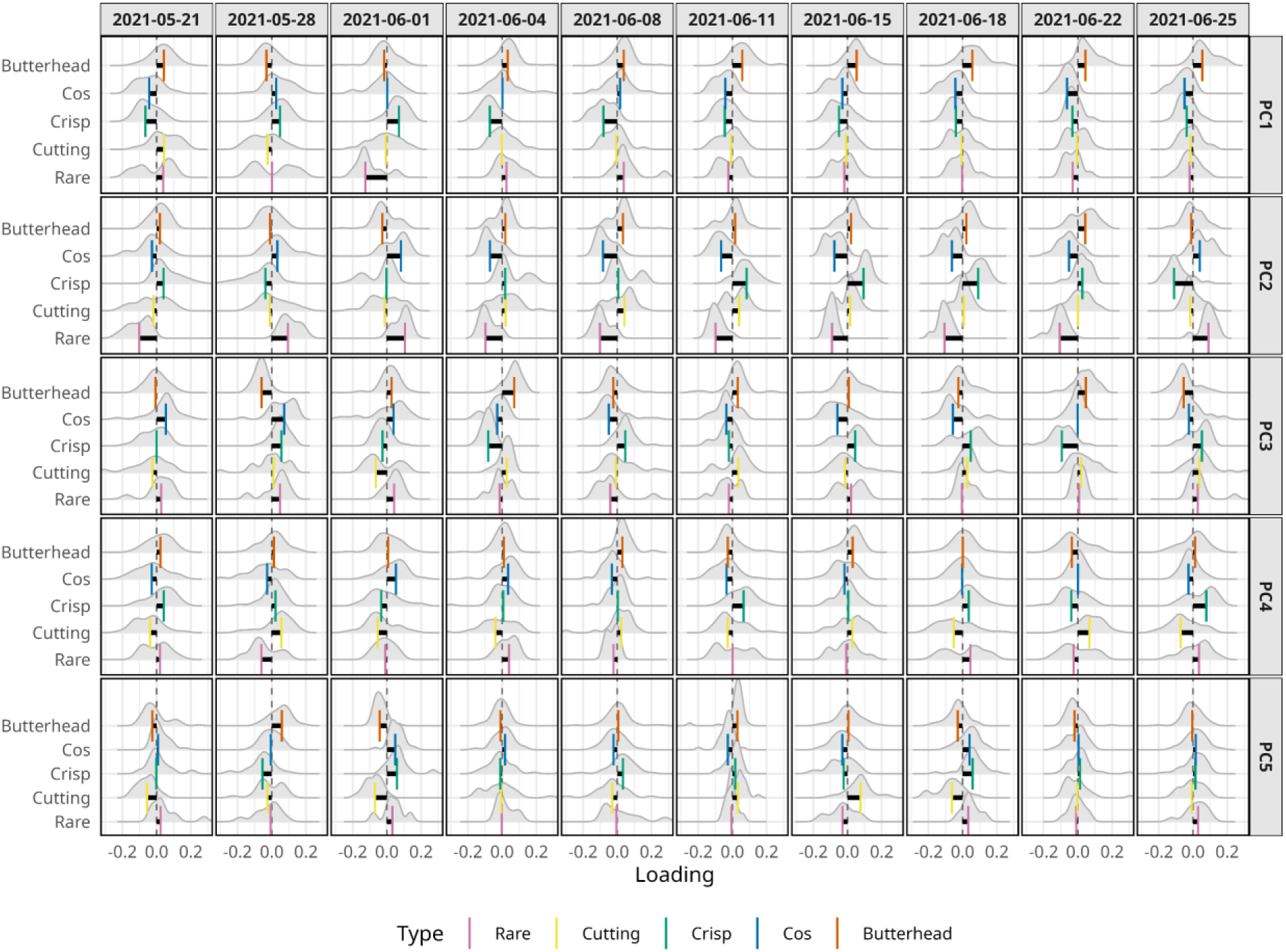
Distribution of reciprocal PCA loadings (PC1 to PC5) across green-only *L. sativa* accessions grouped by horticultural type. Each row represents one principal component (PC1–PC5), and each column corresponds to a timepoint. Density plots show the distribution of loadings per horticultural type, with mean values indicated by colored bars: Butterhead (orange), Cos (blue), Crisp (green), Cutting (yellow), and Rare (purple).

**Figure S19:**
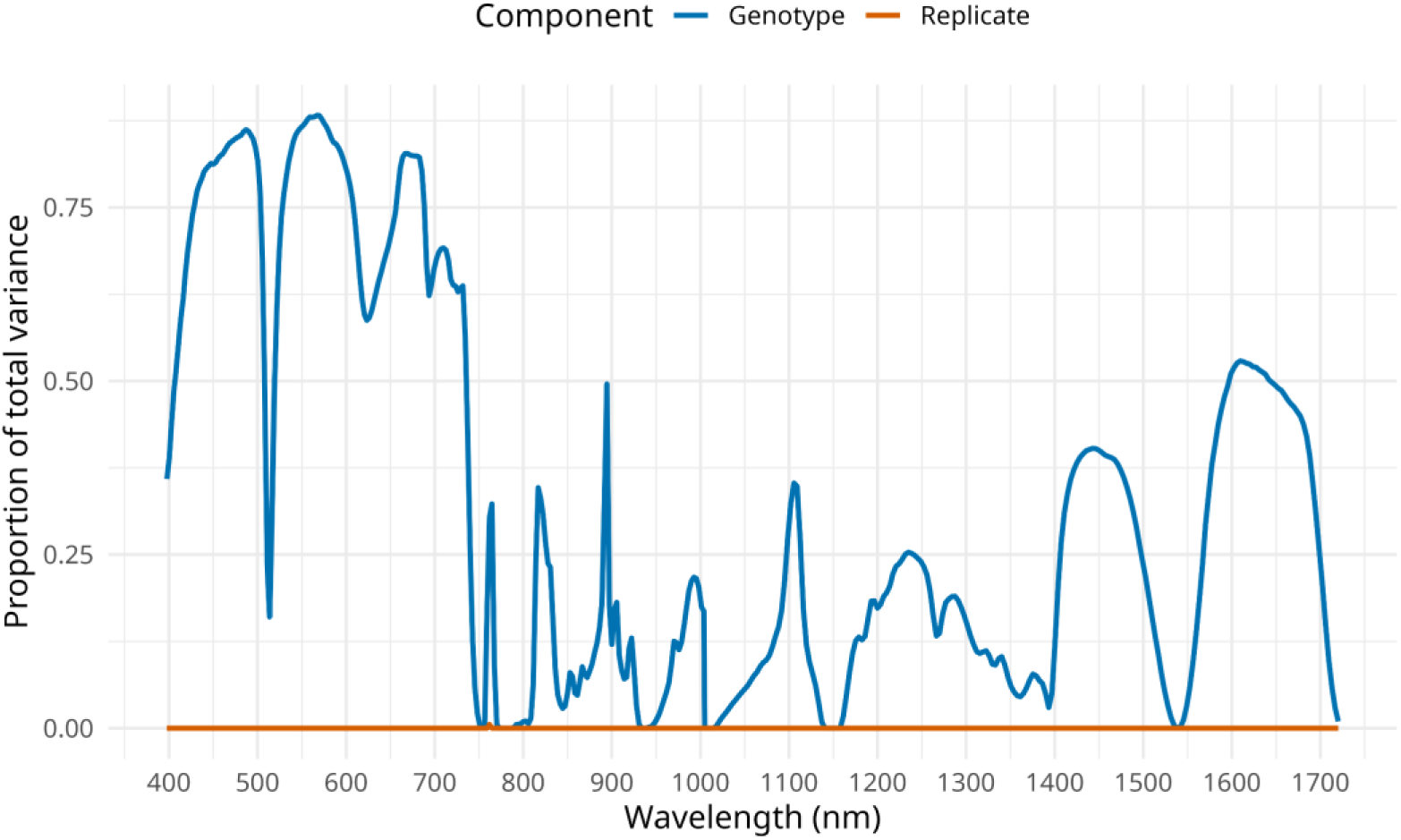
Proportion of variance explained by genotype and replicate. Proportion of variance explained by genotype (blue line) and replicate (orange line) across wavelengths, estimated using a random effects model. The y-axis shows the proportion of variance, and the x-axis represents the individual wavelengths that were modelled.

**Figure S20:**
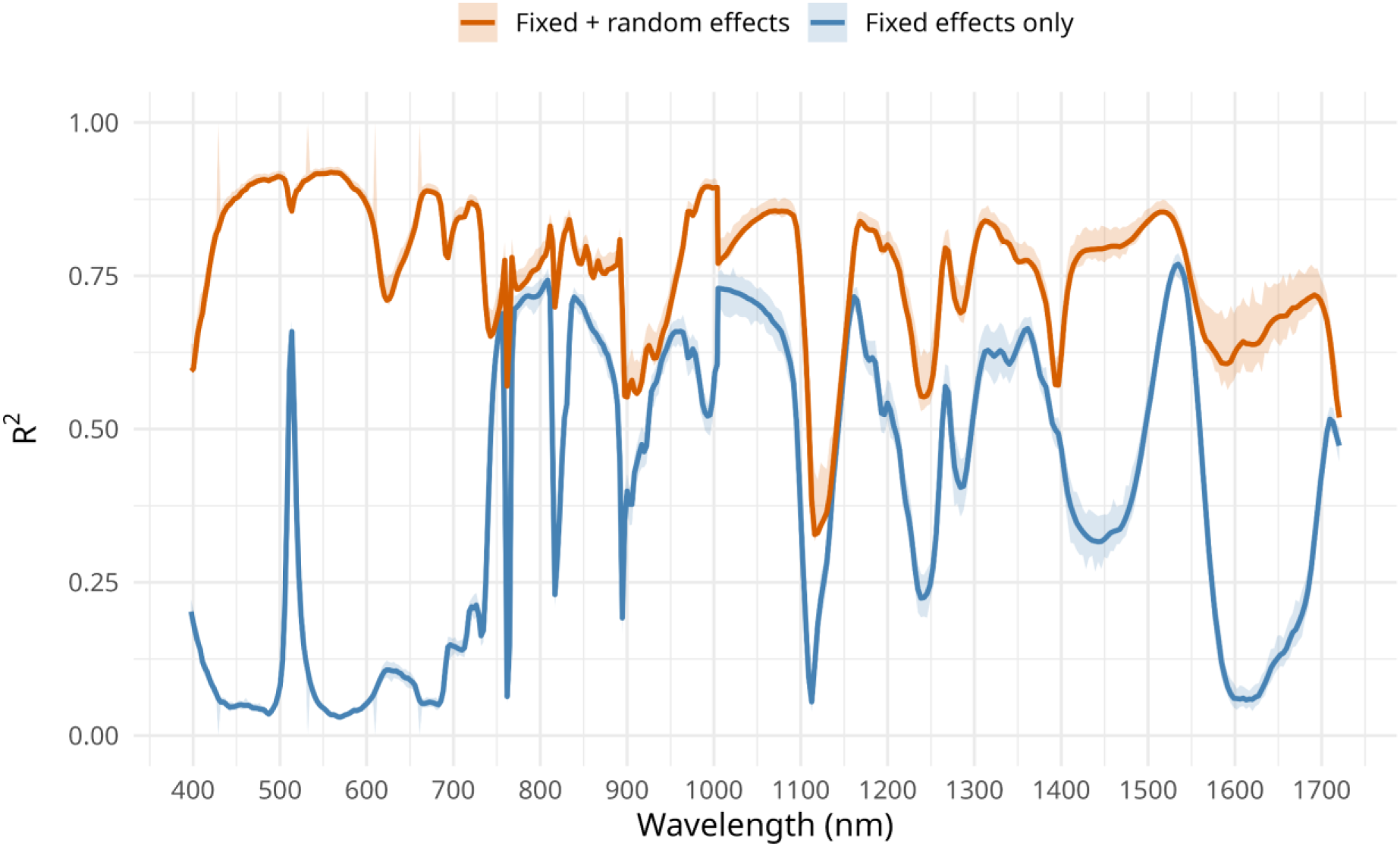
Performance of the full mixed-effects model per wavelength phenotype. The model included time, average air temperature, and cumulative rainfall as fixed effects, with genotype modeled as a random effect. Performance is expressed using Nakagawa’s R², with the marginal R² (blue line) representing variance explained by fixed effects only, and the conditional R² (orange line) representing variance explained by both fixed and random effects combined. Shaded areas indicate 95% confidence intervals derived from bootstrapping (100 iterations). The x-axis shows the wavelength (nm), and the y-axis shows the proportion of variance explained.

**Figure S21:**
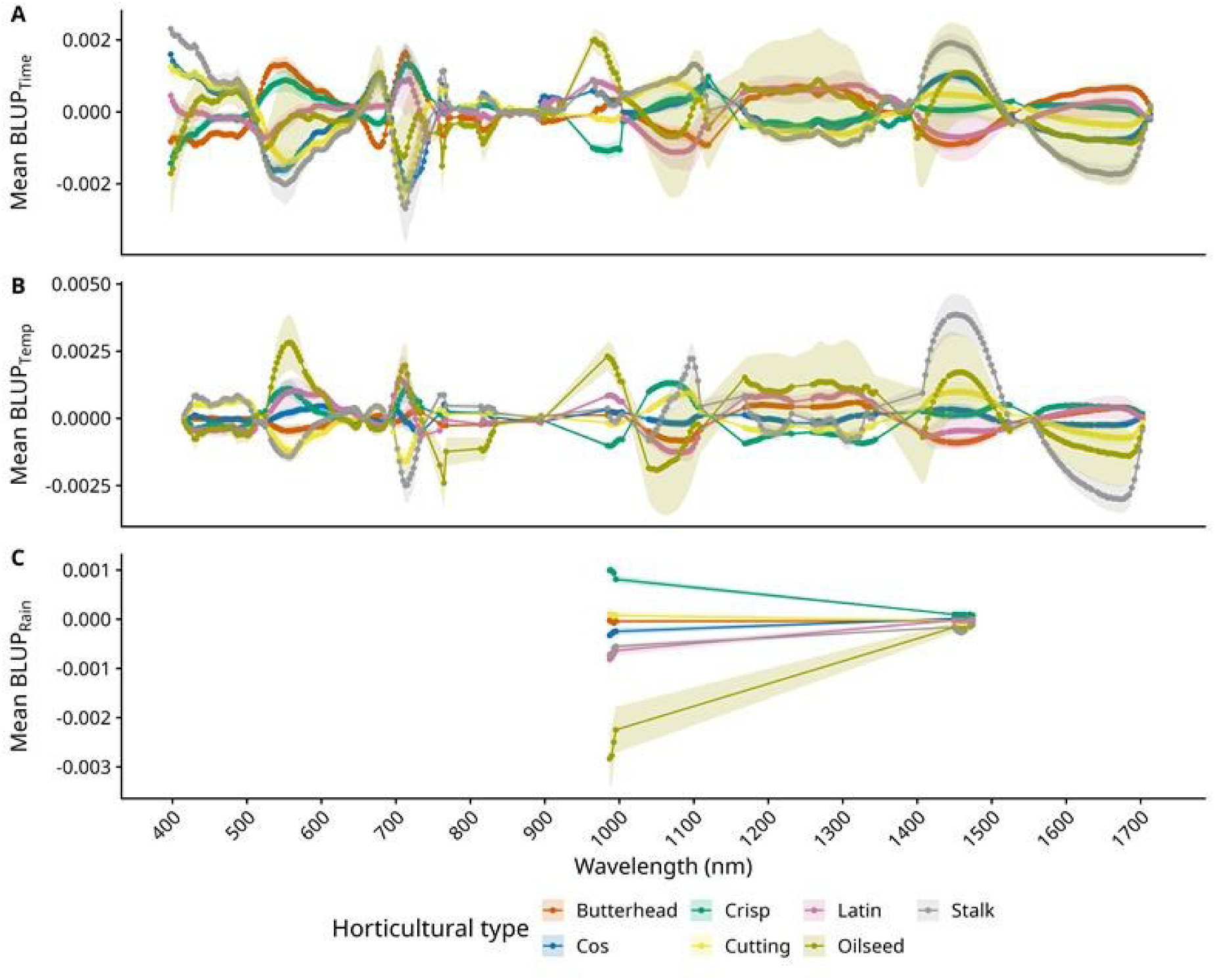
Mean BLUPs (Best Linear Unbiased Predictiors) across wavelengths for genotype-by-environment interaction components time, air temperature and rainfall in green *L. sativa* accessions. BLUPs were extracted from linear mixed models with stable fits (x-axis), then rescaled to reflect the estimated change in reflectance per day (A), per °C (B) or per mm (C) (y-axis). Each colored line represents a different horticultural type of lettuce: Butterhead (red), Crisp (green), Latin (pink), Stalk (grey), Cos (blue), Cutting (yellow), and Oilseed (dark yellow), with shaded areas indicating standard deviation.

**Figure S22:**
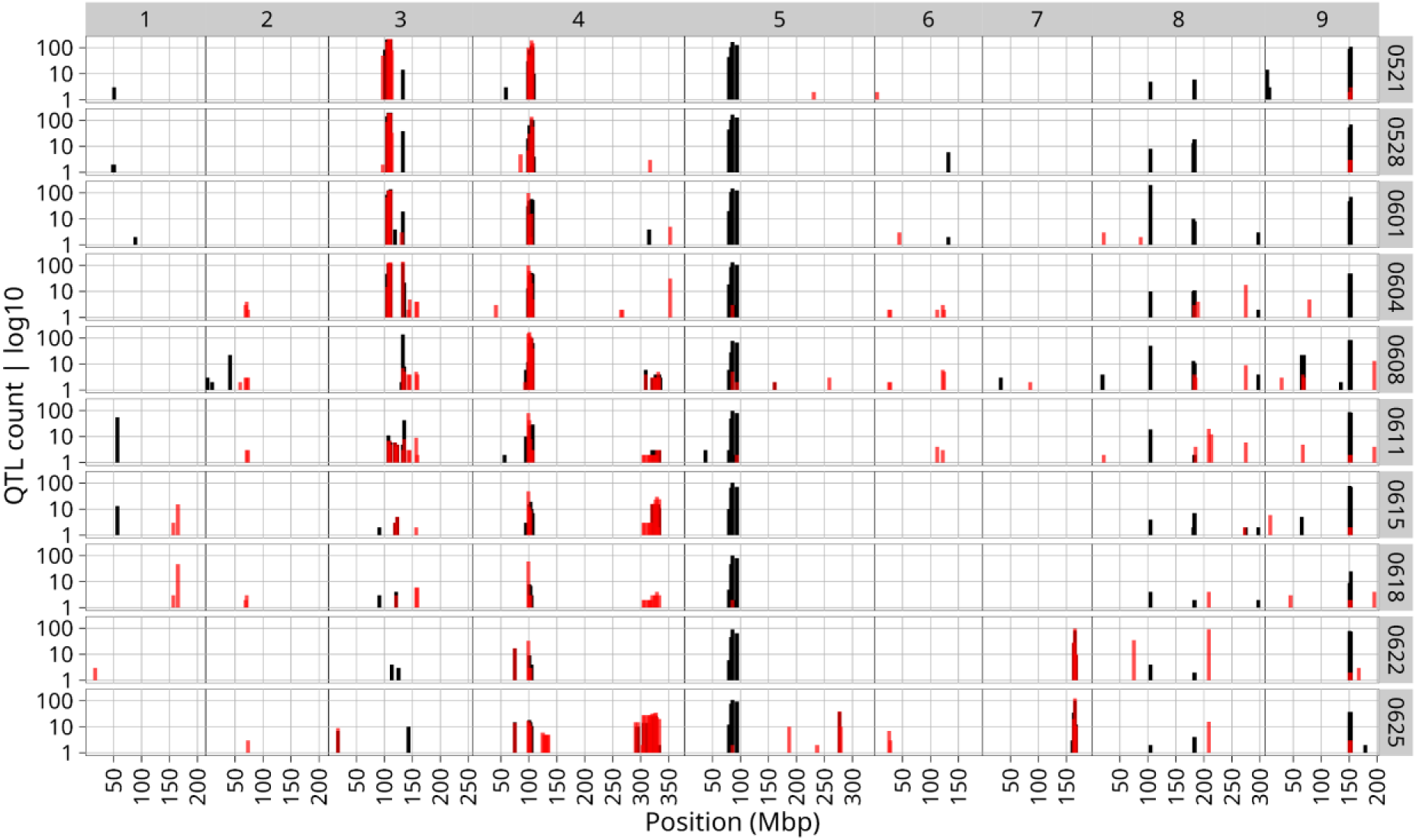
Histogram of QTLs found for all spectral and vegetation index traits and time points. QTLs were defined by the most significant SNP (-log_10_(P) > 7.69, Bonferroni corrected) per 2Mbp per trait per time point. The x-axis shows genomic position in mega base pairs, with chromosome numbers indicated above, the y-axis shows the QTL count. Note that the y-axis is log10 scaled. Timepoints are indicated on the right. Bar color corresponds to QTLs found with GWAS with all accessions (black), and QTLs found with GWAS in green accessions (red).

**Figure S23:**
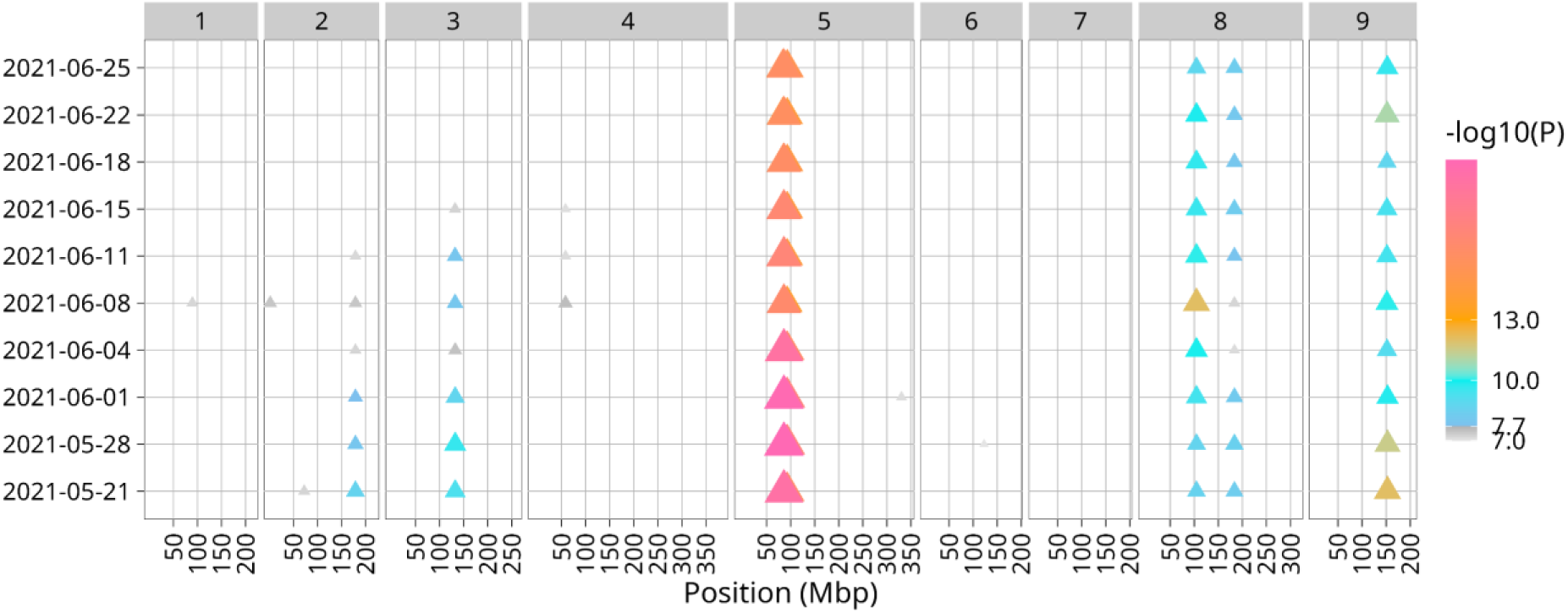
GWAS of anthocyanin reflectance index (ARI) in all *L. sativa* accessions over time. Multi-manhattan depicts QTLs associated with ARI variation per timepoint (left). The x-axis shows genomic position in mega base pairs, with chromosome numbers indicated above. Significant QTLs (–log₁₀(P) > 7.69, Bonferroni-corrected) are marked as triangles, where color and size correspond to the -log10(P) value.

**Figure S24:**
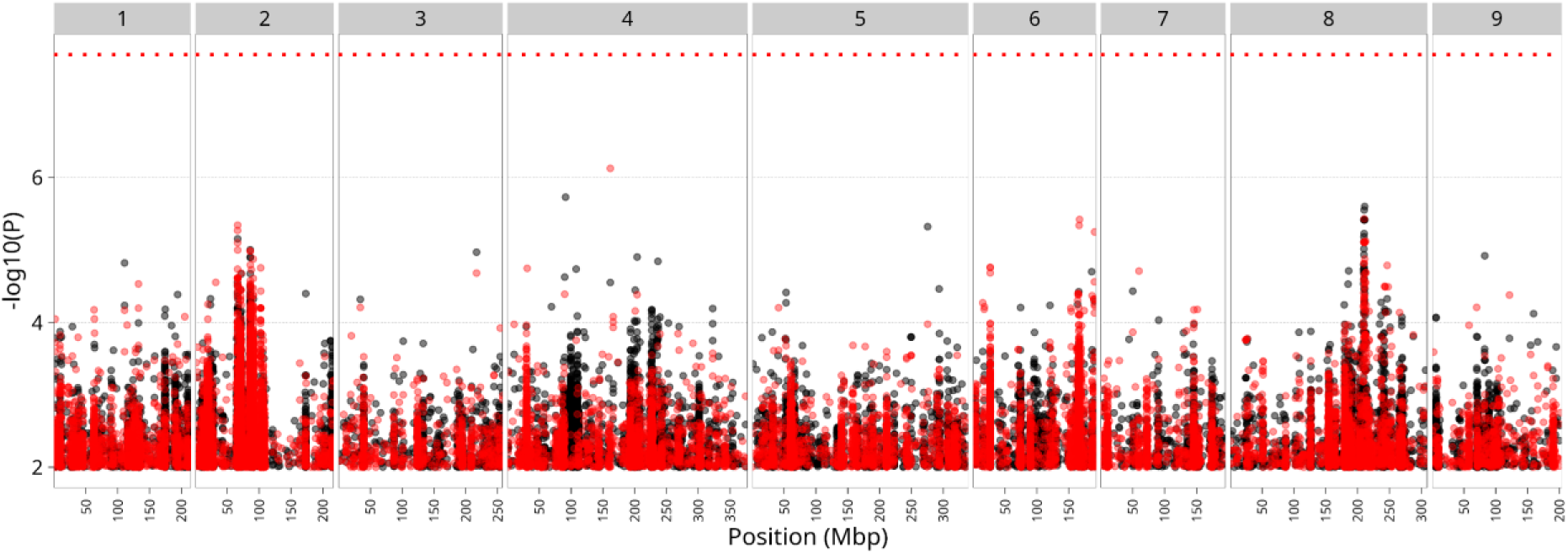
GWAS result for ARI BLUP_Time_ in green *>L. sativa* accessions. The Manhattan plot depicts SNPs associated with the ARI BLUP_Time_ phenotype of green accessions. Significance is plotted as –log₁₀(p-value) on the y-axis. The x-axis shows genomic position in mega base pairs, with chromosome numbers indicated above. The red horizontal line shows the Bonferroni corrected significance threshold (–log₁₀(P) > 7.69). The coloring corresponds to iterative GWAS, where black is the result of the first iteration, and red is the result of the second iteration, where the top SNP of chromosome 9 was added as a fixed effect.

**Figure S25:**
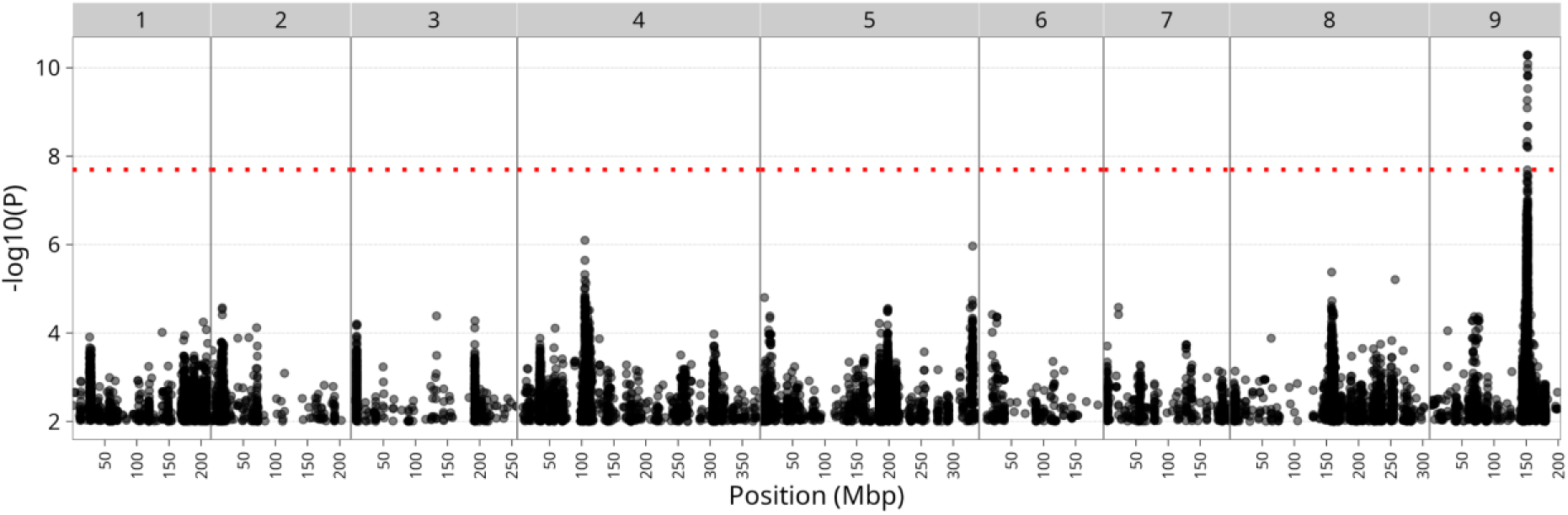
GWAS result for 556.25nm BLUP_Temp_ in green *L. sativa* accessions. The Manhattan plot depicts SNPs associated with the 556.25nm BLUP_Temp_ phenotype of green accessions. Significance is plotted as –log₁₀(p-value) on the y-axis. The x-axis shows genomic position in mega base pairs, with chromosome numbers indicated above. The red horizontal line shows the Bonferroni corrected significance threshold (–log₁₀(P) > 7.69).

**Figure S26:**
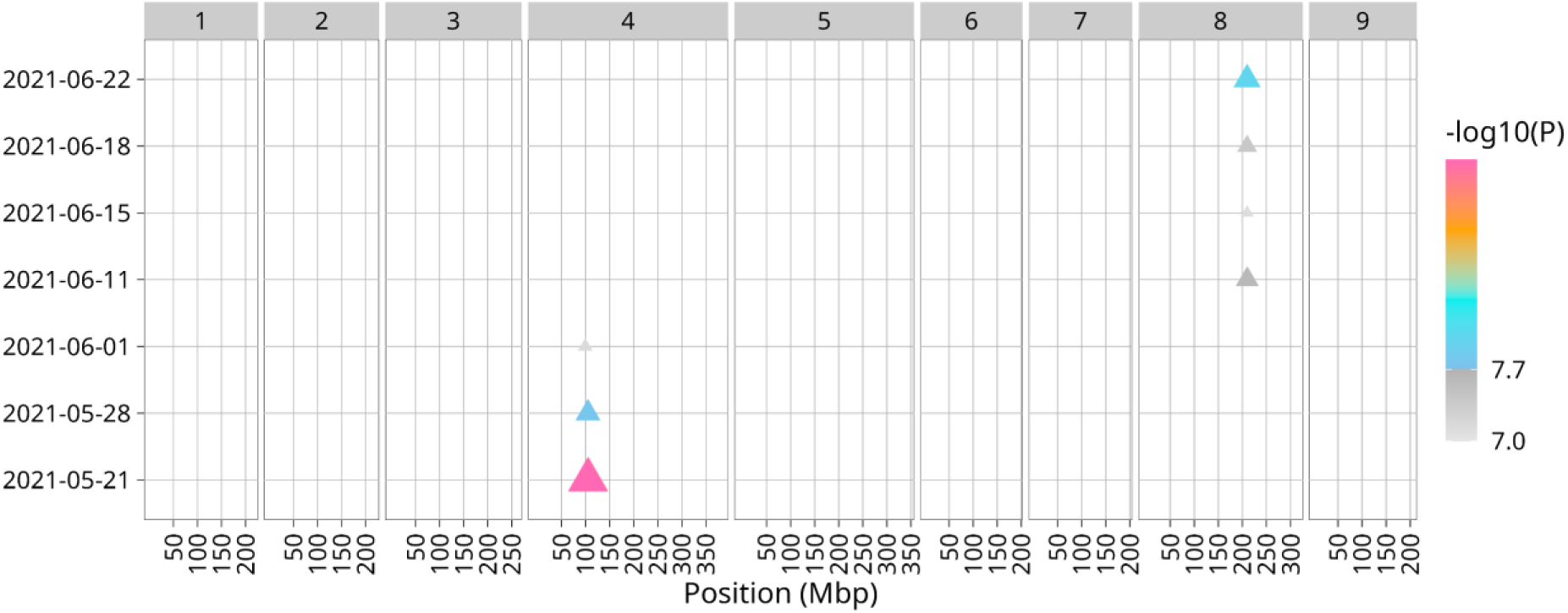
GWAS of reflectance at 556.25nm over time in green *L. sativa* accessions. Multi-manhattan depicts QTLs associated with variation in reflectance at 556.25nm in *L. sativa* accessions over time (indicated left). The x-axis shows genomic position in mega base pairs, with chromosome numbers indicated above. Significant QTLs (–log₁₀(P) > 7.69, Bonferroni-corrected) are marked as triangles, where color and size correspond to the -log10(P) value.

**Figure S27:**
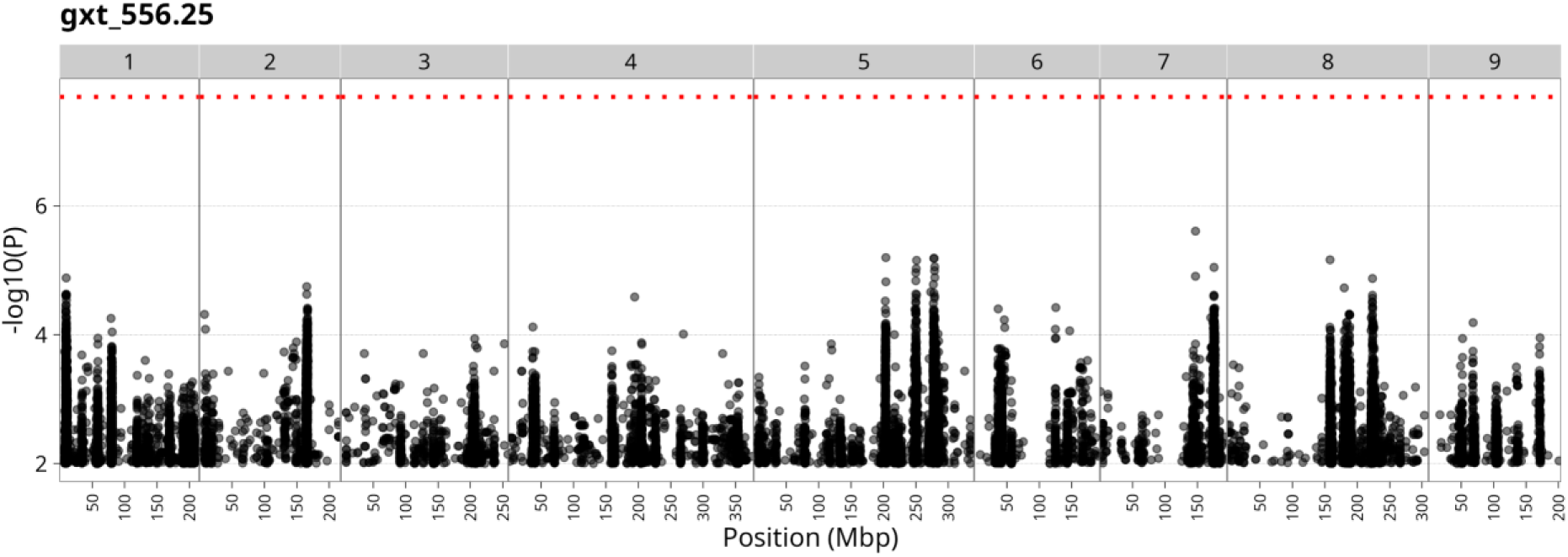
GWAS result for 556.25nm BLUP_Time_ in green *L. sativa* accessions. The Manhattan plot depicts SNPs associated with the 556.25nm BLUP_Time_ phenotype of green accessions. Significance is plotted as –log₁₀(p-value) on the y-axis. The x-axis shows genomic position in mega base pairs, with chromosome numbers indicated above. The red horizontal line shows the Bonferroni corrected significance threshold (–log₁₀(P) > 7.69).

## Notes

https://github.com/SnoekLab/Hyperspec_Mehrem_etal_2025

### Competing Interest Statement

The authors have declared no competing interest.

## References

1. Hultgren, A., et al., Impacts of climate change on global agriculture accounting for adaptation. Nature, 2025. 642(8068): p. 644–652.

2. Hartman, Y., et al., Abiotic stress QTL in lettuce crop–wild hybrids: comparing greenhouse and field experiments. Ecology and Evolution, 2014. 4(12): p. 2395–2409.

3. Yamashita, H., et al., Deciphering transcriptomic signatures explaining the phenotypic plasticity of nonheading lettuce genotypes under artificial light conditions. Plant, Cell & Environment, 2023. 46(12): p. 3971–3985.

4. Zhao, Y., et al., Exploring phenotypic differences and dynamic associations among lettuce types based on high-throughput phenotyping platform. Computers and Electronics in Agriculture, 2025. 236: p. 110454.

5. Chen, H., et al., Dissecting the genetic architecture of key agronomic traits in lettuce using a MAGIC population. Genome Biology, 2025. 26(1): p. 67.

6. Jangra, S., et al., High-Throughput Phenotyping: A Platform to Accelerate Crop Improvement. Phenomics, 2021. 1(2): p. 31–53.

7. Sarić, R., et al., Applications of hyperspectral imaging in plant phenotyping. Trends in Plant Science, 2022. 27(3): p. 301–315.

8. van Eeuwijk, F.A., et al., Modelling strategies for assessing and increasing the effectiveness of new phenotyping techniques in plant breeding. Plant Science, 2019. 282: p. 23–39.

9. Moreira, F.F., et al., Integrating High-Throughput Phenotyping and Statistical Genomic Methods to Genetically Improve Longitudinal Traits in Crops. Frontiers in Plant Science, 2020. Volume 11 - 2020.

10. Hobby, D., A.J. Mbebi, and Z. Nikoloski, Towards genetic architecture and genomic prediction of crop traits from time-series data: Challenges and breakthroughs. Journal of Plant Physiology, 2025. 312: p. 154566.

11. Yu, S., et al., Hyperspectral Technique Combined With Deep Learning Algorithm for Prediction of Phenotyping Traits in Lettuce. Frontiers in Plant Science, 2022. Volume 13 - 2022.

12. Gonçalves, J.V., et al. Mapping of Leaf Pigments in Lettuce via Hyperspectral Imaging and Machine Learning. Horticulturae, 2025. 11, DOI: 10.3390/horticulturae11091077.

13. Lord, E., et al., Early detection of multiple biotic stressors in lettuces using hyperspectral imaging and deep learning. 2025. p. 139–146.

14. Dijkhuizen, R.F., et al., From aerial drone to quantitative trait locus: leveraging next-generation phenotyping to reveal the genetics of color and height in field-grown Lactuca sativa. The Plant Journal, 2025. 123(3): p. e70405.

15. Wei, T., et al., Whole-genome resequencing of 445 Lactuca accessions reveals the domestication history of cultivated lettuce. Nature Genetics, 2021. 53(5): p. 752–760.

16. Zhang, L., et al., RNA sequencing provides insights into the evolution of lettuce and the regulation of flavonoid biosynthesis. Nature Communications, 2017. 8(1): p. 2264.

17. Su, W., et al., Characterization of four polymorphic genes controlling red leaf colour in lettuce that have undergone disruptive selection since domestication. Plant Biotechnology Journal, 2020. 18(2): p. 479–490.

18. Zhang, L., et al., Alternative splicing triggered by the insertion of a CACTA transposon attenuates LsGLK and leads to the development of pale-green leaves in lettuce. The Plant Journal, 2022. 109(1): p. 182–195.

19. R Development Core Team, R: A language and environment for statistical computing. 2022, R Foundation for Statistical Computing: Vienna, Austria.

20. Barthelme, S., imager: Image Processing Library Based on ‘CImg*’*.

21. Vallejos, R., F. Osorio, and M. Bevilacqua, Spatial relationships between two georeferenced variables. 1 ed. 2020, Cham, Switzerland: Springer Nature. 194.

22. Tuszynski, J., caTools: Tools: Moving Window Statistics, GIF, Base64, ROC AUC, etc.

23. Barnes, R.J., M.S. Dhanoa, and S.J. Lister, Standard Normal Variate Transformation and De-Trending of Near-Infrared Diffuse Reflectance Spectra. Applied Spectroscopy, 1989. 43(5): p. 772–777.

24. Montero, D., et al., A standardized catalogue of spectral indices to advance the use of remote sensing in Earth system research. Scientific Data, 2023. 10(1): p. 197.

25. Gu, Z., Complex heatmap visualization. iMeta, 2022. 1(3): p. e43.

26. Bates, D., et al., Fitting Linear Mixed-Effects Models Using lme4. Journal of Statistical Software, 2015. 67(1): p. 1–48.

27. Lüdecke, D., et al., *performance: An R Package for Assessment*, Comparison and Testing of Statistical Models. Journal of Open Source Software, 2021. 6(60): p. 3139.

28. Arnold, P.A., L.E.B. Kruuk, and A.B. Nicotra, How to analyse plant phenotypic plasticity in response to a changing climate. New Phytologist, 2019. 222(3): p. 1235–1241.

29. van Workum, D.M., et al., Dataset: Structural, functional and evolutionary characterisation of genes in Lactuca sp. reference genomes in the context of eudicots. 2024, 4TU.ResearchData.

30. Poplin, R., et al., Scaling accurate genetic variant discovery to tens of thousands of samples. bioRxiv, 2018.

31. Ziyatdinov, A., et al., lme4qtl: linear mixed models with flexible covariance structure for genetic studies of related individuals. BMC Bioinformatics, 2018. 19(1): p. 68.

32. Tange, O., GNU Parallel - The Command-Line Power Tool. The USENIX Magazine, 2011. 36(1): p. 42–47.

33. Kuang, K., Q. Kong, and F. Napolitano, pbmcapply: Tracking the Progress of Mc*pply with Progress Bar. 2022.

34. Mehrem, S.L., G. van den Ackerveken, and B.L. Snoek, Natural variation in seed coat color in lettuce and wild Lactuca species. bioRxiv, 2024: p. 2024.06.27.600409.

35. van Workum, D.J.M., et al., Lactuca super-pangenome reduces bias towards reference genes in lettuce research. BMC Plant Biology, 2024. 24(1): p. 1019.

36. Mehrem, S., G. Ackerveken, and B. Snoek, Phenotypic variation across Lactuca species and genome-wide association analysis in L. sativa and L. serriola. Euphytica, 2026. 222.

37. Gao, B., NDWI—A normalized difference water index for remote sensing of vegetation liquid water from space. Remote Sensing of Environment, 1996. 58(3): p. 257–266.

38. Haboudane, D., et al., Integrated narrow-band vegetation indices for prediction of crop chlorophyll content for application to precision agriculture. Remote sensing of environment, 2002. 81(2-3): p. 416–426.

39. Gitelson, A.A., M.N. Merzlyak, and O.B. Chivkunova, Optical properties and nondestructive estimation of anthocyanin content in plant leaves. Photochemistry and Photobiology, 2001. 74(1): p. 38–45.

40. Jing, Y. and R. Lin, The VQ Motif-Containing Protein Family of Plant-Specific Transcriptional Regulators. Plant Physiol, 2015. 169(1): p. 371–8.

41. Li, Y., et al., H(+)-ATPases in Plant Growth and Stress Responses. Annual Review of Plant Biology, 2022. 73: p. 495–521.

42. Qin, F., et al., Arabidopsis DREB2A-interacting proteins function as RING E3 ligases and negatively regulate plant drought stress-responsive gene expression. Plant Cell, 2008. 20(6): p. 1693–707.

43. Cai, H., et al., Epigenetic regulation of anthocyanin biosynthesis by an antagonistic interaction between H2A.Z and H3K4me3. New Phytologist, 2019. 221(1): p. 295–308.

44. Aharoni, A., et al., The SHINE clade of AP2 domain transcription factors activates wax biosynthesis, alters cuticle properties, and confers drought tolerance when overexpressed in Arabidopsis. Plant Cell, 2004. 16(9): p. 2463–80.

45. Park, S., et al., Genome-wide characterization and evolutionary analysis of the AP2/ERF gene family in lettuce (Lactuca sativa). Scientific Reports, 2023. 13(1): p. 21990.

46. Datt, B., Remote Sensing of Chlorophyll a, Chlorophyll b, Chlorophyll a+b, and Total Carotenoid Content in Eucalyptus Leaves. Remote Sensing of Environment, 1998. 66(2): p. 111–121.

47. Kume, A., T. Akitsu, and K.N. Nasahara, Why is chlorophyll b only used in light-harvesting systems? Journal of Plant Research, 2018. 131(6): p. 961–972.

48. Blackburn, G.A., Spectral indices for estimating photosynthetic pigment concentrations: A test using senescent tree leaves. International Journal of Remote Sensing, 1998. 19(4): p. 657–675.

49. Yang, Y., et al. Rapid and Nondestructive Evaluation of Wheat Chlorophyll under Drought Stress Using Hyperspectral Imaging. International Journal of Molecular Sciences, 2023. 24, DOI: 10.3390/ijms24065825.

50. Patiluna, V., et al. Using Hyperspectral Imaging and Principal Component Analysis to Detect and Monitor Water Stress in Ornamental Plants. Remote Sensing, 2025. 17, DOI: 10.3390/rs17020285.

51. Kumar, P., et al., Molecular Mapping of Water-Stress Responsive Genomic Loci in Lettuce (Lactuca spp.) Using Kinetics Chlorophyll Fluorescence, Hyperspectral Imaging and Machine Learning. Front Genet, 2021. 12: p. 634554.

52. Furlanetto, R.H., et al., Hyperspectral reflectance imaging to classify lettuce varieties by optimum selected wavelengths and linear discriminant analysis. Remote Sensing Applications: Society and Environment, 2020. 20: p. 100400.

53. Cotrozzi, L. and J.J. Couture, Hyperspectral assessment of plant responses to multi-stress environments: Prospects for managing protected agrosystems. PLANTS, PEOPLE, PLANET, 2020. 2(3): p. 244–258.

54. Salson, M., et al., Interplay between large low-recombining regions and pseudo-overdominance in a plant genome. Nature Communications, 2025. 16(1): p. 6458.

55. Cao, S., N. Sawettalake, and L. Shen, Lactuca super-pangenome provides insights into lettuce genome evolution and domestication. Nature Communications, 2025. 16(1): p. 7257.

56. Rönspies, M., et al., CRISPR–Cas-mediated chromosome engineering for crop improvement and synthetic biology. Nature Plants, 2021. 7(5): p. 566–573.

